# The long non-coding RNA *GHSROS* reprograms prostate cancer cell lines toward a more aggressive phenotype

**DOI:** 10.1101/682203

**Authors:** Patrick B. Thomas, Penny L. Jeffery, Manuel D. Gahete, Eliza J. Whiteside, Carina Walpole, Michelle L. Maugham, Lidija Jovanovic, Jennifer H. Gunter, Elizabeth D. Williams, Colleen C. Nelson, Adrian C. Herington, Raúl M. Luque, Rakesh N. Veedu, Lisa K. Chopin, Inge Seim

## Abstract

It is now appreciated that long non-coding RNAs (lncRNAs) are important players in the orchestration of cancer progression. In this study we characterized *GHSROS*, a human lncRNA gene on the opposite DNA strand (antisense) to the ghrelin receptor gene, in prostate cancer. The lncRNA was upregulated by prostate tumors from different clinical datasets. Consistently, transcriptome data revealed that *GHSROS* alters the expression of cancer-associated genes. Functional analyses *in vitro* showed that *GHSROS* mediates tumor growth, migration, and survival and resistance to the cytotoxic drug docetaxel. Increased cellular proliferation of *GHSROS*-overexpressing PC3, DU145, and LNCaP prostate cancer cell lines *in vitro* was recapitulated in a subcutaneous xenograft model. Conversely, *in vitro* antisense oligonucleotide inhibition of the lncRNA reciprocally regulated cell growth and migration, and gene expression. Notably, *GHSROS* modulates the expression of PPP2R2C, the loss of which may drive androgen receptor pathway-independent prostate tumor progression in a subset of prostate cancers. Collectively, our findings suggest that *GHSROS* can reprogram prostate cancer cells toward a more aggressive phenotype and that this lncRNA may represent a potential therapeutic target.

## INTRODUCTION

The human genome yields a multitude of RNA transcripts with no obvious protein-coding ability, collectively termed non-coding RNAs (ncRNAs)^1^. A decade of intensive research has revealed that many ncRNAs greater than 200 nucleotides in length have expression patterns and functions as diverse as protein-coding RNAs^1, 2^. These long non-coding RNAs (lncRNAs) have emerged as important regulators of gene expression, acting on nearby (*cis*) or distant (*trans*) protein-coding genes^2^. Although the vast majority of lncRNAs remain uncharacterized, it is clear that they play key regulatory roles in development, normal physiology, and disease.

We previously^3^ identified *GHSROS* (also known as *AS-GHSR*), a 1.1-kb capped and polyadenylated lncRNA gene antisense to the intronic region of the ghrelin receptor gene (*GHSR*) (Fig. 1a). *GHSROS* harbors a putative human-specific promoter in a transposable element^3^, a pattern frequently found in promoters of lncRNAs with high tissue specificity and low expression levels^4, 5^. It is now appreciated that many lncRNAs are equivalent to classical oncogenes or tumor suppressors and drive similar transcriptional programs in diverse cancer types^2^. Indeed, our earlier study showed that *GHSROS* is overexpressed in lung cancer and that its forced overexpression increases migration in lung adenocarcinoma cells lines^3^. We speculated that *GHSROS* plays a role in other cancers. Prostate cancer is a disease diagnosed in nearly 1.5 million men worldwide annually^6^. Intriguing recent studies have revealed that, like breast cancer, prostate cancer is a heterogeneous disease with multiple molecular phenotypes^7, 8, 9^. The identification of genes that drive or mediate these distinct phenotypes is crucial. Although a number of lncRNAs have been reported in prostate cancer, few have been functionally characterized or assessed as therapeutic targets^10^. Here, we report that *GHSROS* is highly expressed in a subset of prostate tumors. We provide evidence that this lncRNA reprograms prostate cancer cells toward a more aggressive phenotype, possibly by repressing the expression of the tumor suppressor PPP2R2C to allow androgen-independent growth.

**Figure 1.**
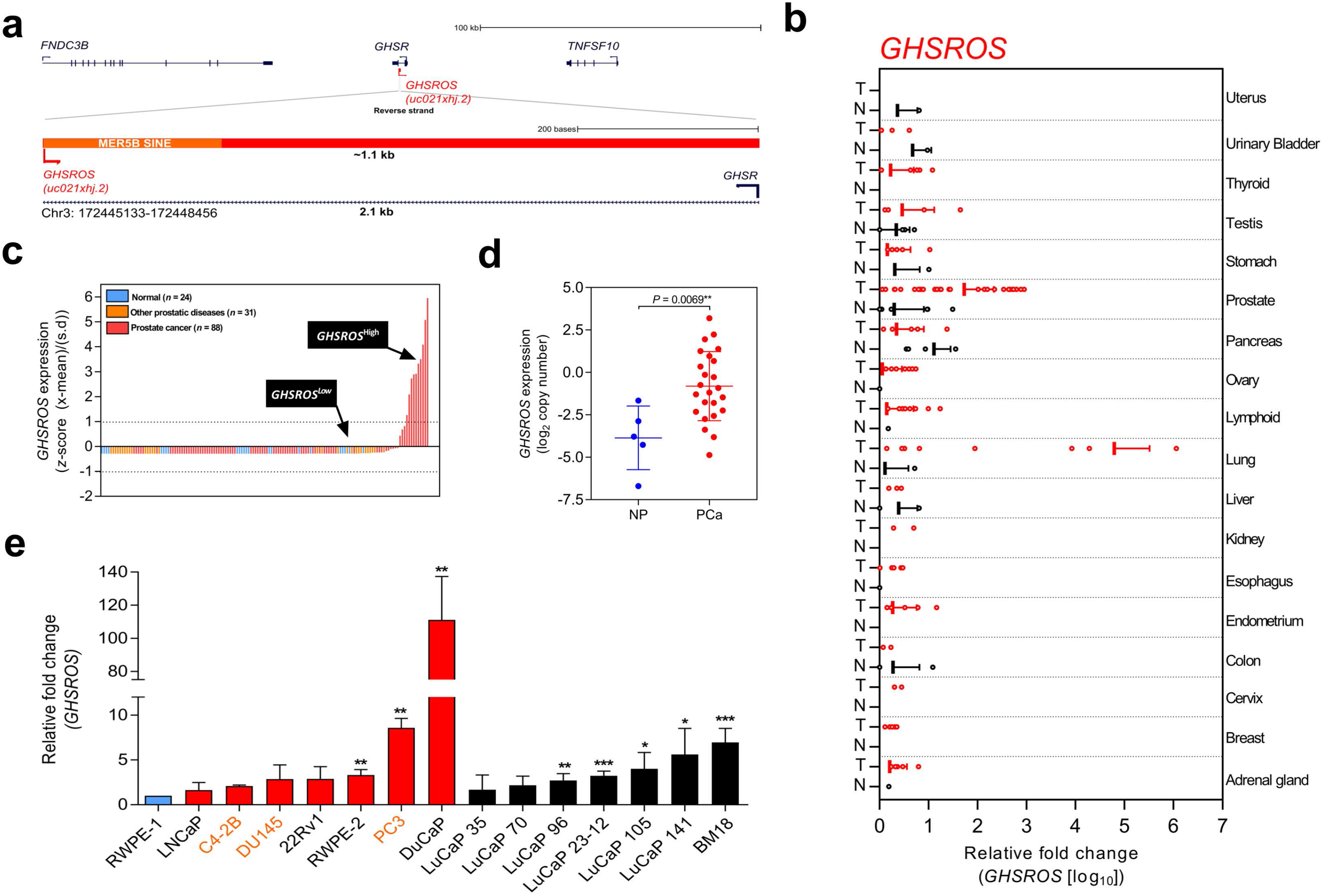
Overview of the lncRNA *GHSROS* and its expression in cancer. **(a)** Overview of the *GHSR* and *GHSROS* gene loci. *GHSR* exons (black), *GHSROS* exon (red), repetitive elements (orange), introns (lines). **(b)** *GHSROS* expression in 19 cancers (TissueScan Cancer Survey Tissue qPCR panel). N (black) denotes normal tissue; T tumor (red). For each cancer, data are expressed as mean fold change using the comparative 2^-ΔΔCt^ method against a non-malignant control tissue. Normalized to β-actin (*ACTB*). **(c)** Relative gene expression of *GHSROS* in OriGene cDNA panels of tissues from normal prostate (*n*=24; blue), primary prostate cancer (*n*=88; red), and other prostatic diseases (*n*=31; orange). Determined by qRT-PCR, normalized to ribosomal protein L32 (*RPL32*), and represented as standardized expression values (*Z*-scores). **(d)** *GHSROS* expression in an Andalusian Biobank prostate tissue cohort. Absolute expression levels were determined by qRT-PCR and adjusted by a normalization factor calculated from the expression levels of three housekeeping genes (*HPRT*, *ACTB*, and *GAPDH*). NP denotes non-malignant prostate. \**P*≤0.05, Mann-Whitney-Wilcoxon test. **(e)** Expression of *GHSROS* in immortalized, cultured cell lines and patient-derived xenograft (PDX) lines. Mean ± s.e.m. (*n*=3). \**P*≤0.05, \*\**P*≤0.01, \*\*\**P*≤0.001, Student’s *t*-test. Normalized as in (b) to the RWPE-1 non-malignant cell line. Androgen-independent lines are labeled in orange.

## RESULTS

### *GHSROS* is expressed in prostate cancer

Microarrays and RNA-sequencing are commonly used to assess the expression of genes. LncRNAs are often expressed at orders of magnitude lower than protein-coding transcripts, however, making them difficult to detect^5, 11, 12, 13, 14, 15^. Interrogation of exon arrays harboring four different strand-specific probes against *GHSROS* demonstrated that the lncRNA is actively transcribed, although expressed at very low levels in cancer cell lines and tissues (Supplementary Fig. S1), consistent with previous observations from Northern blotting and RT-PCR experiments^3^. The low expression across the *GHSROS* and *GHSR* loci in RNA-seq datasets is illustrated in Supplementary Fig. S2. Collectively, these data demonstrate that it is not currently possible to detect *GHSROS* in public genome-wide gene expression datasets.

We next evaluated *GHSROS* expression in a qRT-PCR tissue array of 18 cancers. This analysis revealed particularly high *GHSROS* expression in lung tumors, as previously reported^3^, and elevated expression in prostate tumors (Fig. 1b). Analysis of additional prostate tissue-derived cDNA arrays revealed that *GHSROS* could be detected in approximately 41.7% of all normal prostate tissues (*n*=24), 55.7% of tumors (*n*=88), and 58.1% of other prostatic diseases (e.g. prostatitis; *n*=31) (Supplementary Table S1). *GHSROS* was highly expressed by a subset of prostate tumors (∼11.4%; *Z*-score >1) (Fig. 1c) and elevated in tumors with Gleason scores 8-10 (Supplementary Fig. S3; Supplementary Table S1; Mann-Whitney-Wilcoxon test *P*=0.0021). To expand on these observations, we examined an independent cohort of eight normal prostate tissue specimens and 28 primary tumors with high Gleason scores (18 of which had metastases at biopsy). Similarly, *GHSROS* expression was significantly elevated in tumors compared to normal prostate tissue (Mann-Whitney-Wilcoxon test, *P*=0.0070) (Fig. 1d; Supplementary Fig. S4; Supplementary Table S2).

As the functional thresholds of long non-coding RNAs are difficult to gauge and likely cell-context specific^16^, we identified cell lines with a range of endogenous *GHSROS* expression. Compared to the RWPE-1 benign prostate-derived cell line, higher expression was observed in the PC3 (*P*=0.00040, Student’s *t-*test) (Fig. 1e) and DuCaP prostate cancer cell lines (*P*=0.0024), and expression was similar to RWPE-1 in the DU145 (*P*=0.29) and LNCaP prostate cancer cell lines (*P*=0.49). We also assessed the expression of *GHSROS* in patient-derived xenografts (PDXs). Compared to RWPE-1, *GHSROS* was significantly upregulated (*P≤*0.05) in 4/6 of the LuCaP series of PDX lines^17^ and in the BM18 femoral metastasis-derived androgen-responsive PDX line^18^ (*P=*0.0005) (Fig. 1e).

### *GHSROS* promotes growth and motility of prostate cancer cells *in vitro*

To gain insights into *GHSROS*, we assessed its function in three prostate-derived cell lines by stably overexpressing the lncRNA in PC3, DU145, and LNCaP cells (denoted PC3-GHSROS, DU145-GHSROS, and LNCaP-GHSROS) (Supplementary Fig. S5). Cell proliferation over 72 hours (measured by a xCELLigence real-time cell analysis instrument) was increased in PC3 (*P*=0.029, Student’s t-test) and DU145 (*P*=0.026) *GHSROS*-overexpressing cells (Fig. 2a). LNCaP cells did not attach well to the gold electrodes of the xCELLigence instrument (data not shown), and we therefore utilized a WST-1 assay to assess this cell line. Similar to PC3 and DU145 cells overexpressing *GHSROS*, proliferation was also increased in LNCaP-GHSROS cells at 72 hours (*P*=0.040) (Fig. 2b). *GHSROS* overexpression also increased the rate of cell migration of PC3 (*P*=0.0064, Student’s t-test), DU145 (*P*=0.017), and LNCaP cells (*P*=0.00020) over 24 hours (Fig. 2c) (where LNCaP was assessed by a standard transwell migration assay; PC3 and DU145 by an xCELLigence instrument). To confirm the *in vitro* functional effects of *GHSROS*, we designed locked nucleic antisense oligonucleotides (LNA-ASOs) to strand-specifically silence endogenous *GHSROS* expression (Fig. 2d; Supplementary Fig. S6). Two LNA-ASOs targeting distinct regions of *GHSROS*, RNV124 and RNV104L, independently reduced the expression of *GHSROS* (percentage knockdown of ∼63% and ∼71%, respectively) in native PC3 cells 48 hours post transfection compared to scrambled control (*P*=0.0002 and *P*=0.0001, Student’s *t*-test) (Fig. 2e). Moreover, *GHSROS* knockdown attenuated cell proliferation (RNV124, *P*=0.049; RNV104L, *P*=0.030) (Fig. 2f) and migration of PC3 cells over 18 hours (RNV124, *P*=0.0042) (Fig. 2g) – the reciprocal effects observed when *GHSROS* was forcibly overexpressed.

**Figure 2.**
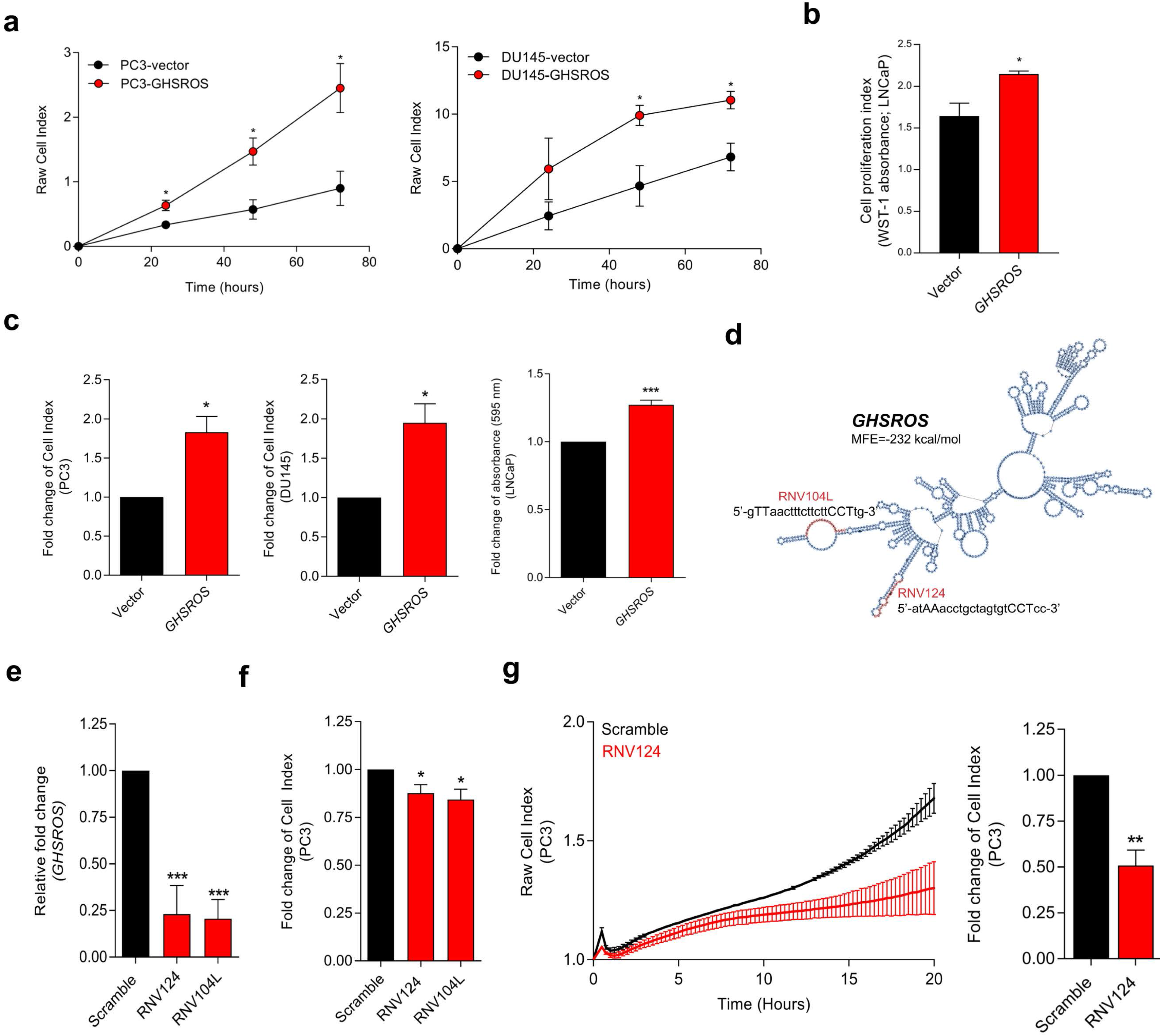
*GHSROS* promotes human prostate cancer cell line growth and motility *in vitro*. **(a, b)** Increased proliferation by *GHSROS*-overexpressing cells. PC3 and DU145 cells were assessed using an xCELLigence real-time cell analyzer for 72 hours; LNCaP using a WST-1 assay at 72 hours. Vector denotes empty control plasmid. Mean ± s.e.m. (*n*=3). \**P*≤0.05, \*\**P*≤0.01, \*\*\**P*≤0.001, Student’s *t*-test. **(c)** Increased migration by *GHSROS*-overexpressing cells. PC3 and DU145 cells were assessed using an xCELLigence real-time cell analyzer for 24 hours; LNCaP using a transwell assay (at 24 hours; *n*=3). Parameters and annotations as in (a). **(d)** *GHSROS* RNA secondary structure prediction. The location of locked nucleic antisense oligonucleotides (LNA-ASOs) that target the lncRNA are shown in red. MFE denotes minimum free energy. **(e)** LNA ASOs reduced *GHSROS* expression by PC3 cells (measured 48 hours post-transfection). Fold-enrichment of *GHSROS* normalized to *RPL32* and compared to scrambled control (*n*=3). Parameters and annotations as in (a). **(f)** *GHSROS* knockdown reduces PC3 proliferation (*n*=3). Parameters and annotations as in (a). **(g)** *GHSROS* knockdown reduces PC3 migration. Left panel: representative plot of cell index impedance measurements from 0 to 20 hours after transfection of LNA-ASO RNV124 (*n*=3). Right panel: RNV124 reduced cell migration at 18 hours (*n*=3). Parameters and annotations as in (c).

### *GHSROS* is associated with cell survival and resistance to the cytotoxic drug docetaxel

Knockdown experiments also revealed that *GHSROS* protected PC3 prostate cancer cells from death by serum starvation (Supplementary Fig. S7). This observation led us to examine whether *GHSROS* contributes to cell survival following chemotherapy. The current treatment of choice for advanced, castration-resistant prostate cancer (CRPC; the fatal final stage of the disease) after the failure of hormonal therapy is the cytotoxic drug docetaxel, a semi-synthetic taxoid that induces cell cycle arrest. At the half maximal inhibitory concentration (IC_50_) of docetaxel (5 nM for LNCaP^19^), survival was significantly increased in *GHSROS*-overexpressing LNCaP cells (*P*≤0.05, Student’s *t*-test) (Fig. 3a) after 96 hours. A similar, less pronounced response was observed in LNCaP cells treated with enzalutamide, a hormonal therapy used to target the androgen receptor in metastatic, castration-resistant tumors^20^ (Fig. 3a).

**Figure 3.**
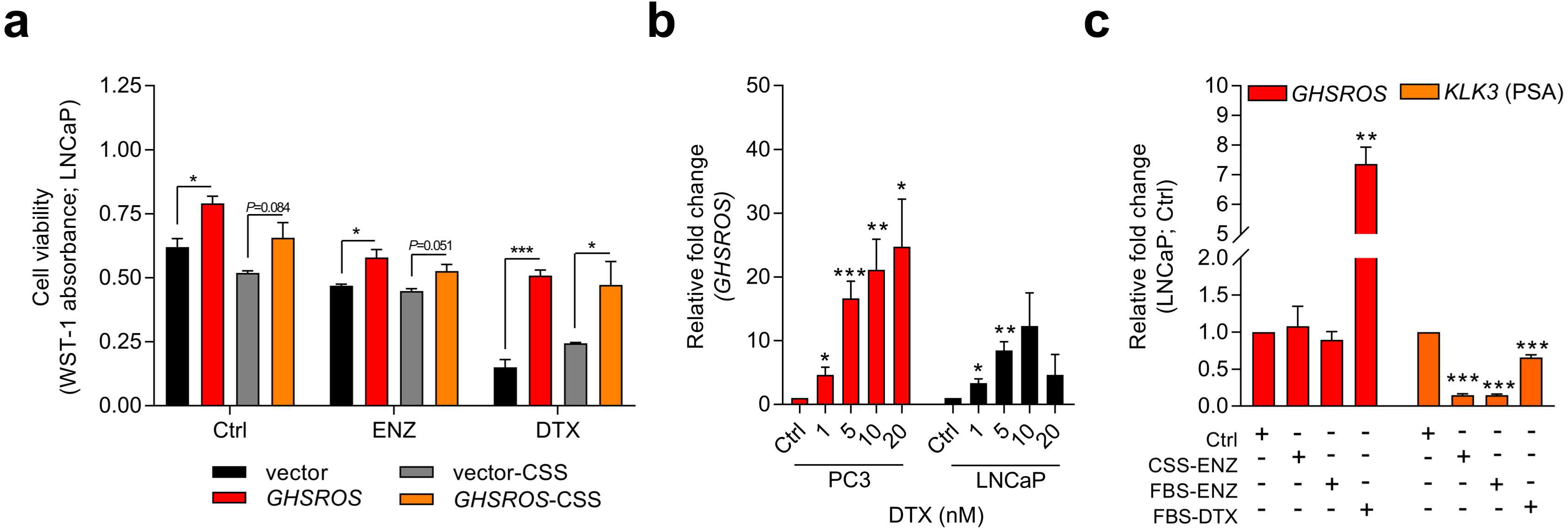
*GHSROS* mediates cell survival and resistance to the cytotoxic drug docetaxel. **(a)** Viability of *GHSROS*-overexpressing LNCaP cells under different culture conditions. Cell number was assessed using WST-1. Cells were treated with enzalutamide (ENZ; 10 µM) or docetaxel (DTX; 5 nM) for 96 hours and grown in either 2% FBS or 5% charcoal stripped serum (CSS) RPMI-1640 media (*n*=3). Mean ± s.e.m. \**P*≤0.05, \*\*\**P*≤0.001, Student’s *t*-test. **(b)** *GHSROS* expression of native PC3 and LNCaP cells treated with docetaxel. Cells were grown in RPMI-1640 media with 2% FBS and treated with 1-20 nM docetaxel (DTX) for 48 hours (*n*=3). Fold-enrichment of *GHSROS* normalized to *RPL32* and compared to empty vector control. Parameters and annotations as in (a). **(c)** *GHSROS* and PSA (*KLK3*) expression of native LNCaP cells treated with ENZ (10 µM in 2% FBS or 5% CSS RPMI-1640) or DTX (5 nM in 2% FBS RPMI-1640) for 48 hours (*n*=3). Parameters and annotations as in (a).

Survival pathways are induced after docetaxel treatment in prostate cancer^21, 22^, and resistance may develop after chemotherapy (acquired resistance) or exist in treatment-naïve patients (innate resistance)^21^. The pronounced survival following docetaxel treatment in *GHSROS*-overexpressing LNCaP cells led us to speculate that endogenous *GHSROS* expression also contributed to drug resistance. Docetaxel significantly increased *GHSROS* expression in native LNCaP and PC3 cells – in a dose-dependent manner and at concentrations both above and below their respective IC_50_ values (Fig. 3b). The lncRNA was not differentially expressed in charcoal stripped serum (CSS), used to simulate androgen deprivation therapy, or following treatment with enzalutamide (Fig. 3c). In agreement with previous reports^23, 24^, the gene coding for prostate specific antigen (PSA; *KLK3*) was downregulated by docetaxel and enzalutamide in LNCaP cells (−6.6-fold, *P*=0.00070, Student’s *t*-test) (Fig. 3c). Taken together, these data suggest that *GHSROS* mediates tumor survival and resistance to the cytotoxic chemotherapy docetaxel.

### *GHSROS* potentiates tumor growth *in vivo*

In order to firmly establish a role for *GHSROS* in tumor growth, we established subcutaneous *GHSROS*-overexpressing androgen-independent (PC3 and DU145) and androgen-responsive (LNCaP) cell line xenografts in NOD/SCIDIL2Rγ (NSG) mice. Subcutaneous graft sites allow easy implantation and monitoring of tumor growth (using calipers)^25^ – ideal for exploring the role of a new gene such as *GHSROS in vivo*. Overexpression of *GHSROS* in xenografts was confirmed post-mortem by qRT-PCR (Supplementary Fig. S8). Compared to vector controls, xenograft tumor volumes were significantly greater at day 25 in PC3-GHSROS mice (*P*=0.0040, Mann-Whitney-Wilcoxon test) and at day 35 in DU145-GHSROS mice (*P*=0.0011) (Fig. 4a). While xenograft tumors were not palpable in LNCaP-GHSROS mice *in vivo*, tumors were significantly larger by weight post-mortem (at 72 days) (*P*=0.042, Student’s *t*-test) (Fig. 4b) – with a size increase similar to that seen for DU145-GHSROS xenografts (Fig. 4c). LNCaP-GHSROS tumors invaded the muscle of the flank and the peritoneum (data not shown) and were more vascularized than control tumors (observed grossly and estimated by CD31^+^ immunostaining) (Fig. 4d). Representative Ki67 immunostaining for proliferating xenograft tumor cells is shown in Fig. 4e.

**Figure 4.**
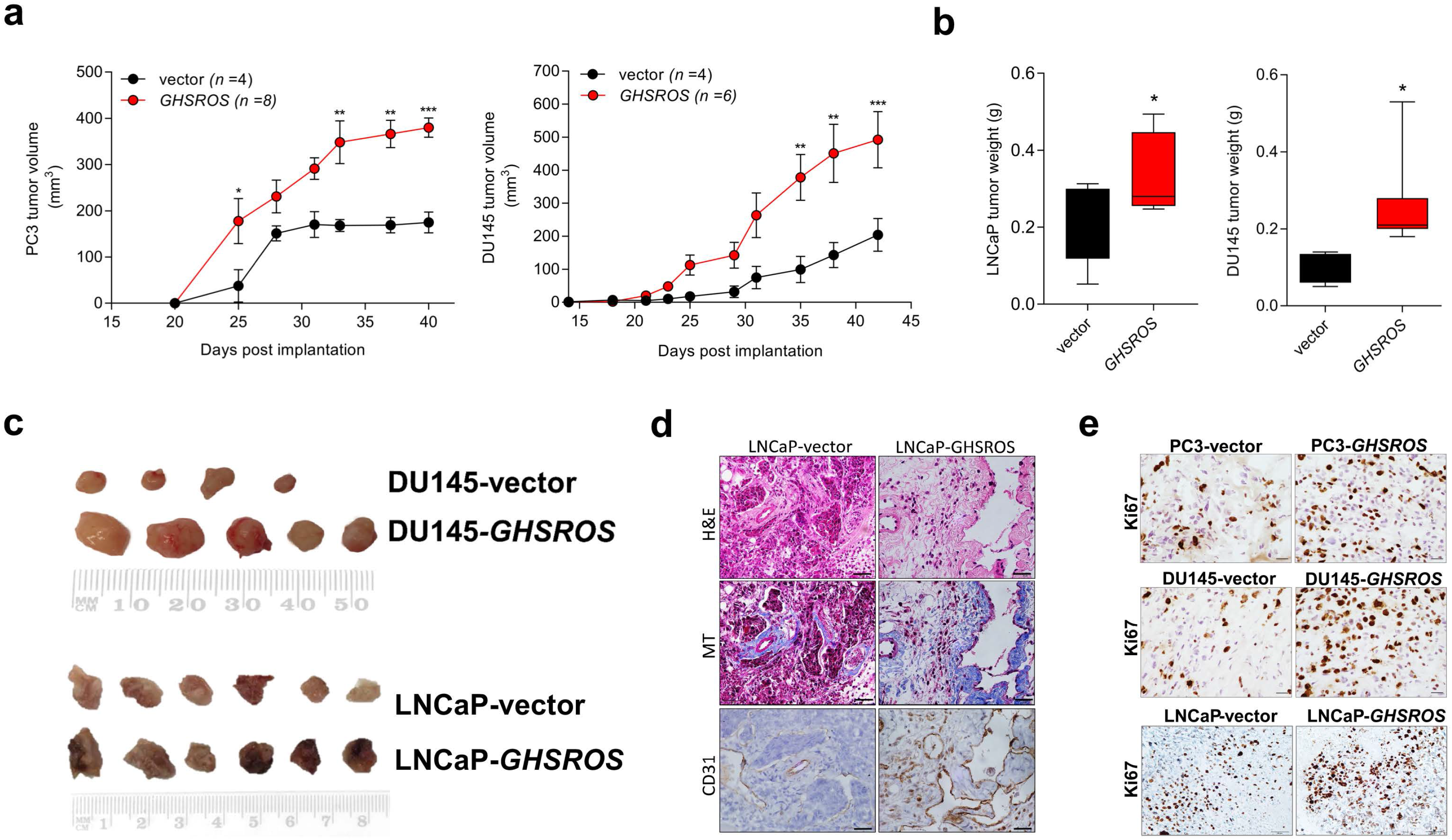
*GHSROS* promotes human prostate cancer cell line growth *in vivo.* **(a)** Left panel: time course for PC3-GHSROS (*n*=8) and vector control (*n*=4) xenograft tumor volumes. Right panel: DU145-GHSROS (*n*=6) and vector control (*n*=4). Mean ± s.e.m. \**P*≤0.05, \*\**P*≤0.01, \*\*\**P*≤0.001, two-way ANOVA with Bonferonni’s *post hoc* analysis. Tumors were measured with digital calipers. **(b)** Tumor weights of LNCaP (left panel; *GHSROS*-overexpressing *n*=9, vector *n*=8) or DU145 (right panel; see (a)). \**P*≤0.05, Mann-Whitney-Wilcoxon test. **(c)** Size comparisons of DU145 (top panel) and LNCaP (bottom panel) xenografts overexpressing *GHSROS* or empty vector. **(d)** Representative morphology of LNCaP xenografts overexpressing *GHSROS* or empty vector. Tissue was stained with hematoxylin and eosin (H&E), Masson’s Trichrome (MT; collagen; blue) and CD31 (endothelial marker; brown immunoreactivity). Scale bar = 20 μm. **(e)** Representative Ki67 immunostaining of PC3 xenografts (top), DU145 xenografts (middle), and LNCaP xenografts (bottom). Scale=20 µm.

### *GHSROS* modulates the expression of cancer-associated genes

Having established that *GHSROS* plays a role in regulating hallmarks of cancer – including cell proliferation, invasion, and migration^26^ – we sought to determine the genes likely to mediate its function by examining the transcriptomes of cultured PC3 cells and LNCaP xenografts overexpressing this lncRNA.

High-throughput RNA-seq of cultured PC3-GHSROS cells (∼50M reads) revealed that 400 genes were differentially expressed (168 upregulated, 232 downregulated, moderated *t*-test; cutoff set at log_2_ fold-change ± 1.5, *Q*≤0.05) (Supplementary Table S3) compared with empty vector control cells. In support of our functional data, gene ontology analysis using DAVID showed enrichment for cancer, cell motility, cell migration, and regulation of growth (Supplementary Tables S4 and S5). Given that *GHSROS* is not readily detectable by high-throughput sequencing and array technologies, we queried the 400 genes differentially expressed in PC3-GHSROS cells using Oncomine concept map analysis^27^. Enriched Oncomine concepts included poor clinical outcome and metastatic progression (Fig. 5a; Supplementary Table S6).

**Figure 5.**
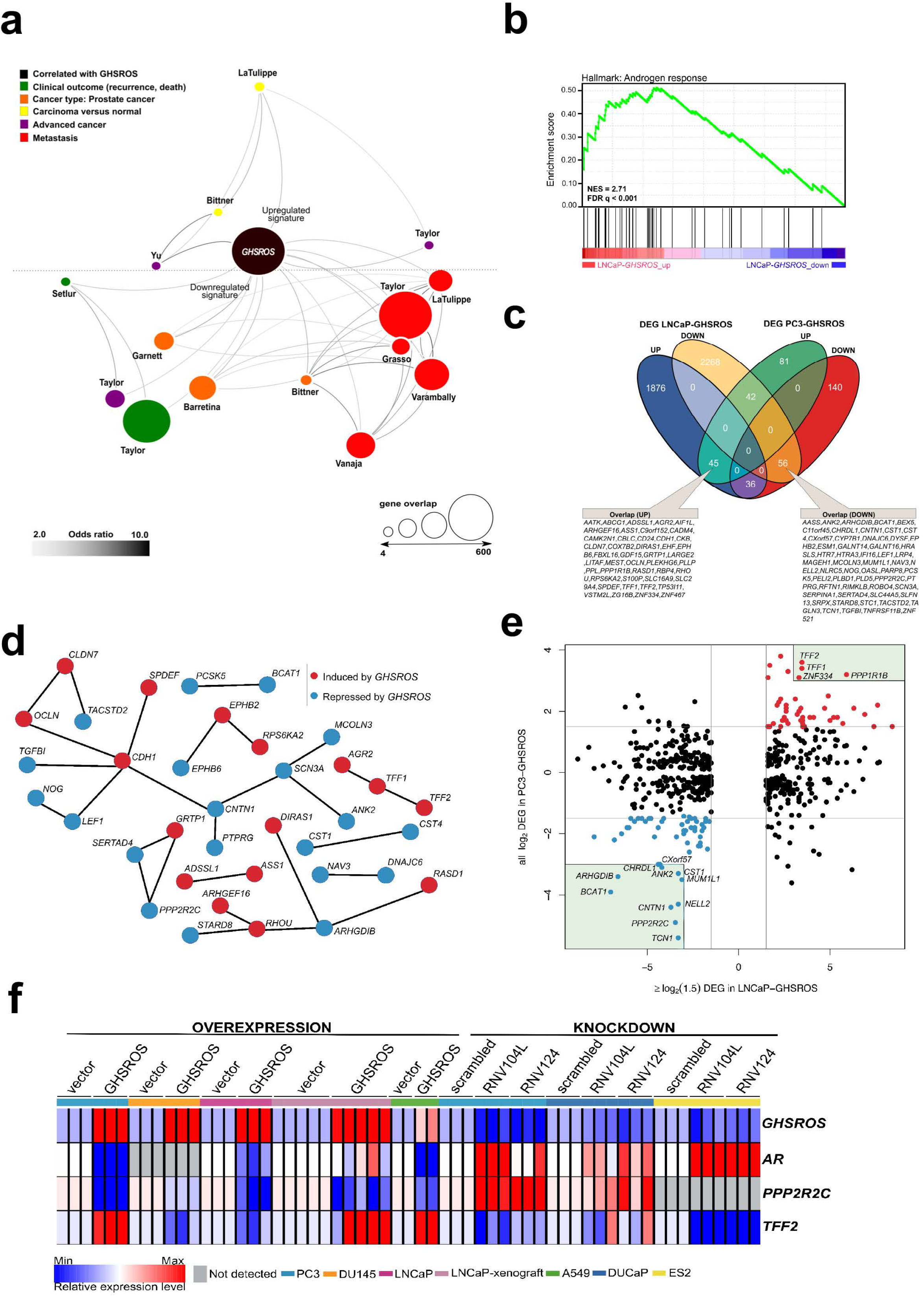
*GHSROS* overexpression modulates the expression of cancer-associated genes. **(a)** Oncomine network representation of genes differentially expressed by cultured PC3-GHSROS cells visualized using Cytoscape. Node sizes (gene overlap) reflect the number of genes per molecular concept. Nodes are colored according to concept categories indicated in the left corner. Edges connect enriched nodes (odds ratio ≥3.0) and darker edge shading indicates a higher odds ratio. **(b)** Gene set enrichment analysis (GSEA) of genes differentially expressed by LNCaP-GHSROS xenografts reveals enrichment for the androgen response. The normalized enrichment score (NES) and GSEA false-discovery corrected *P*-value (*Q*) are indicated. **(c)** Venn diagram of differentially expressed genes (DEG) in LNCaP-GHSROS and PC3-GHSROS cells. Symbols of 101 overlapping genes are indicated in text boxes. **(d)** Interaction of 101 genes differentially expressed in PC3-GHSROS and LNCaP-GHSROS cells (see (c)). Lines represent protein-protein interaction networks from the STRING database. Genes induced (red) or repressed (blue) by *GHSROS*-overexpression are indicated. **(e)** Gene expression scatter plot comparing *GHSROS*-overexpressing PC3 and LNCaP cells. Differentially expressed genes (DEGs) in both datasets shown in red (induced) and blue (repressed); of which ≥8-fold (log_2_ cutoff at −3 and 3) DEGs are highlighted by a green box. **(f)** Heat map of gene expression in *GHSROS*-perturbed cells. Each row shows the relative expression of a single gene and each column a sample (biological replicate). Fold-enrichment of each gene normalized to *RPL32* and compared to empty vector control (overexpression) or scrambled control (LNA-ASO knockdown). Fold-changes were log_2_-transformed and are displayed in the heat map as the relative expression of a gene in a sample compared to all other samples (*Z*-score).

Complementary lower-coverage (∼30M reads) RNA-seq data from LNCaP-GHSROS xenografts demonstrated that a surprisingly large number of genes were differentially expressed (1 961 upregulated, 2 372 downregulated, moderated *t*-test; cutoff set at log_2_ fold^-^ change ± 1.5, *Q*≤0.05) (Supplementary Dataset 1). Selected genes with low expression counts were validated by qRT-PCR (Supplementary Fig. S9). In LNCaP-GHSROS xenografts, *GHSROS*-regulated genes were enriched for the androgen response (gene set enrichment analysis; NES = 2.71, *Q*≤0.001) (Fig. 5b), and included PSA (*KLK3*) (750.9-fold, *Q*=3.6 × 10^-6^) and transmembrane protease serine 2 (*TMPRSS2*) (335.4-fold, *Q*=4.5 × 10^-6^) (Supplementary Dataset 2). We also observed downregulation of numerous genes associated with cell migration and adhesion, epithelial–mesenchymal transition (EMT) (including *ZEB1;* −97.0-fold, *Q*=1.5 × 10^-5^), and angiogenesis and vasculature development (Supplementary Dataset 3). As mentioned above, subcutaneous LNCaP-GHSROS xenografts infiltrated muscle of the flank and the peritoneum and were more vascularized at 72 days post injection in NSG mice, which may indicate that these tumors had completed EMT and angiogenesis at this time.

It is appreciated that the bone metastasis-derived, androgen-independent PC3 and the lymph node metastasis-derived, androgen-responsive LNCaP prostate cancer cell lines represent genetically and presumably metabolically distinct subtypes^28^. They are therefore useful for revealing broad, functional gene expression changes associated with aggressive disease in forced overexpression and knockdown experiments. Despite the differences between these cell lines, a quarter (25.3%; 101 genes) of genes differentially expressed by PC3 cells overexpressing *GHSROS* were also differentially expressed by LNCaP-*GHSROS* cells (Fig. 5c) (*P*=0.000020, hypergeometric test). These genes represent candidate mediators of *GHSROS* function.

We interrogated the STRING database^29^ to reveal protein interactions between the 101 genes regulated by *GHSROS* in both cell lines. A number of genes associated with cell-cell adhesion, migration, and growth were connected, indicating functional enrichment of these proteins in *GHSROS*-overexpressing prostate cancer cells (Fig. 5d). This included increased expression of epithelial cadherin (*CDH1*), occludin (*OCLN*), and claudin-7 (*CLDN7*); and decreased contactin 1 (*CNT1*), noggin (*NOG*), and transforming growth factor beta induced (*TGFBI*) in *GHSROS*-overexpressing cells. Of note, increased *CDH1* expression is associated with exit from EMT and growth of aggressive, metastatic prostate tumors^25^. A second, interesting upregulated network included anterior gradient 2 (*AGR2*) and trefoil factors 1 and 2 (*TFF1* and *TFF2*). Trefoil factors are small proteins associated with mucin glycoproteins. Their expression is increased in castration-resistant prostate cancer (CRPC) and may facilitate the acquisition of hormone independence^31, 32^. Similarly, *AGR2* has been associated with the propensity of a number of aggressive tumor types to metastasize, including prostate cancer^33, 34^.

Ten out of the 101 genes were differentially expressed in metastatic tumors compared to primary tumors in two clinical prostate datasets: Grasso^35^ (59 localized and 35 metastatic prostate tumors) and Taylor^36^ (123 localized and 27 metastatic prostate tumors) (Supplementary Tables S7 and S8) (*Q*≤0.25, moderated *t*-test). *DIRAS1*, *FBXL16*, *TP53I11*, *TFF2*, and *ZNF467* were upregulated in both metastatic tumors and *GHSROS*-overexpressing PC3 and LNCaP cells, while *AASS*, *CHRDL1*, *CNTN1*, *IFI16*, and *MUM1L1* were downregulated. We investigated whether the expression of these genes contributes to adverse disease outcome by assessing survival in two independent datasets: the Taylor dataset and TCGA-PRAD. The latter a dataset of localized prostate tumors generated by The Cancer Genomics Atlas (TCGA) consortium^37^. As overall survival data was available for a small number of patients in these datasets, we assessed disease-free survival (relapse). Relapse is a suitable surrogate for overall survival in prostate cancer given that recurrence of disease would be expected to contribute significantly to mortality, and metastatic disease is incurable. Unsupervised *k*-means clustering was employed to divide each dataset into two groups based on gene expression alone. Two genes, zinc finger protein 467 (*ZNF467*; which was induced by forced *GHSROS*-overexpression) and chordin-like 1 (*CHRDL1*; which was repressed), correlated with relapse in both datasets (Supplementary Table S9). Chordin-like 1 is a negative regulator of bone morphogenetic protein 4-induced migration and invasion in breast cancer^38^. It was downregulated in *GHSROS*-overexpressing cell lines and in metastatic tumors compared to localized tumors in the Taylor and Grasso datasets. Interrogation of the Chandran prostate cancer dataset (60 localized tumors and 63 adjacent, normal prostate)^39^ suggests that *CHRDL1* is downregulated by prostate tumors in general. *CHRDL1* expression stratified the Taylor (*N*=150; 27 metastatic tumors) dataset into two groups with a significant, 438-day difference in overall disease-free survival (relapse; Cox *P*=0.0062, absolute hazard ratio (HR) = 2.5). A statistically significant, yet clinically negligible difference in relapse (9 days; Cox *P*=0.0071, absolute HR = 1.8) was observed in the TCGA-PRAD dataset (*N*=489; no metastatic tumors) (Supplementary Table S9). Survival analysis *P*-values (Kaplan-Meier and Cox proportional-hazard) and hazard ratios indicate whether there is a significant difference between two groups, but not the degree of difference. Evaluating statistically significant differences in survival (e.g. in days) between groups is therefore subjective. Given these data, we propose that *CHRDL1* may play an important role in metastatic tumors.

In contrast to *CHRDL1*, *ZNF467* stratified patients into clusters with an obvious difference in overall median survival (relapse) between groups in both the Taylor (697 days; Cox *P*=0.0039, HR = 2.7) and TCGA-PRAD datasets (139 days; Cox *P*=0.000026, HR = 2.5) (Supplementary Table S9). ZNF467 has not been functionally characterized, however, a recent study suggests that it is a transcription factor which clusters in close proximity to the androgen receptor in a network associated with breast cancer risk^40^, indicating that ZNF467 and AR regulate similar pathways. Clustering of patients into groups of either low or high *ZNF467* expression revealed that elevated expression of the gene associated with a worse relapse outcome (Supplementary Fig. S10a-c). In agreement, *ZNF467* gene expression can distinguish low (≤ 6) from high (≥ 8) Gleason score prostate tumors in a Fred Hutchinson Cancer Research Center prostate cancer dataset (381 localized and 27 metastatic prostate tumors)^41^. *ZNF467* expression is also elevated in chemotherapy-resistant ovarian cancer^42^ and breast cancer^43^ cell lines.

The 101 *GHSROS*-regulated genes were visualized in a scatter plot to reveal genes with particularly distinct (≥ 8-fold) differential expression in *GHSROS*-overexpressing prostate cancer cell lines – putative fundamental drivers of the observed tumorigenic phenotypes. This revealed that *PPP2R2C* (Fig. 5e), a gene encoding a subunit of the holoenzyme phosphatase 2A (PP2A)^44, 45^, was downregulated by forced overexpression of *GHSROS*. In the PC3-GHSROS RNA-seq dataset, *PPP2R2C* was the third most downregulated gene (−29.9-fold, moderated *t*-test *Q*=3.4 × 10^-10^) (Supplementary Table S3). Consistently, forced overexpression or knockdown of *GHSROS* in prostate cancer cell lines reciprocally regulated endogenous *PPP2R2C* expression (Fig. 5f; Supplementary Figs. S9 and S11).

We observed that *GHSROS* was also able to reciprocally regulate androgen receptor (*AR*) expression in some prostate cancer cell lines (downregulated upon *GHSROS* overexpression in PC3 and LNCaP; upregulated upon *GHSROS* knockdown in DUCaP) (Fig. 5f). LNCaP-GHSROS xenografts showed a variable *AR* expression pattern, which may be linked to differences in available androgen, however, *PPP2R2C* expression was still significantly repressed *in vivo* (−3.7-fold, Student’s *t*-test *P*=7.9 × 10^-3^) (Supplementary Fig. S9). Similarly, while *AR* could not be detected in DU145 cells, *GHSROS*-overexpression decreased *PPP2R2C* expression in this cell line (Fig. 5f). The androgen receptor is also expressed by ovarian and lung cancer tumors and cell lines^46, 47^. Forced overexpression of *GHSROS* in the A549 lung adenocarcinoma cell line decreased *AR* and *PPP2R2C* expression (Student’s *t*-test, *P*≤0.0001). *GHSROS* knockdown in the ES-2 ovarian clear cell carcinoma cell line, which does not express *PPP2R2C*, increased the expression of *AR* (Student’s *t*-test, *P*=0.0029 and *P*=0.0022) (Fig. 5f; Supplementary Fig. S11).

## DISCUSSION

Very recent work suggests that a small proportion (∼3%) of long non-coding RNA genes are dysregulated in tumors and mediate cell growth^48^. Herein, we demonstrate that the lncRNA *GHSROS* is one such gene. *GHSROS* expression is elevated across many different cancers, suggesting that it is a so-called pan-cancer lncRNA^49, 50^. In prostate cancer *GHSROS* is detectable in normal tissue and expressed at higher levels in a subset (∼10%) of tumors. We have yet to narrow down on particular prostate tumor strata with elevated *GHSROS*, however.

From assessing the function of *GHSROS* in immortalized prostate cancer cell lines, the following observations were made: Forced overexpression of *GHSROS* enhances *in vivo* tumor growth, and *in vitro* cell viability and motility. We also demonstrate that forced overexpression of *GHSROS* facilitates survival and recalcitrance to the cytotoxic chemotherapy drug docetaxel. Critically, we show that endogenous *GHSROS* is elevated following docetaxel treatment. Docetaxel is commonly prescribed for late-stage, metastatic CRPC patients, but large, randomized trials suggest that it is also effective against recently-diagnosed, localized prostate tumors^51^. These data suggest that *GHSROS* acts as a cell survival factor in prostate cancer. While the underlying mechanisms are unknown, two genes associated with chemotherapy resistance, *ZNF467* and *PPP1R1B* (also known as *DARPP-32*), were upregulated in PC3 and LNCaP cells overexpressing *GHSROS*. *PPP1R1B* is a potent anti-apoptotic gene which confers resistance in cancer cell lines to several chemotherapeutic agents when overexpressed^52^.

The expression and function of *GHSROS* in prostate cancer suggests that it belongs to a growing list of lncRNAs that function as *bona fide* oncogenes. Notable examples associated with aggressive cancer and adverse outcomes include *HOTAIR* (HOX transcript antisense RNA), which is upregulated in a range of cancers^2^, and the prostate cancer-specific *SCHLAP1* (SWI/SNF Complex Antagonist Associated With Prostate Cancer 1)^53^. To better understand how *GHSROS* mediates its effects in prostate cancer, we examined transcriptomes of prostate cancer cell lines with forced *GHSROS* overexpression: PC3 cells in culture (*in vitro*) and subcutaneous LNCaP xenografts in mice (*in vivo*). The 101 common differentially expressed genes included several transcription factors with established roles in prostate cancer and genes associated with metastasis and poor prognosis. Our study not only highlights genes modulated by *GHSROS*, but also genes (such as *ZNF467*, *CHRDL1*, and *PPP2R2C*) that may be generally relevant to prostate cancer progression.

Reactivation of the androgen receptor (*AR*) has long been considered a seminal event; supporting renewed tumor growth in a majority of metastatic CRPC patients^54, 55^. However, it is now increasingly recognized that, similar to other endocrine-related cancers, several subtypes of prostate cancer exist^7, 8, 9^. These include subtypes characterized by androgen pathway-independent growth^44, 56^. In this context, our results on *PPP2R2C,* a gene which encodes a PP2A substrate-binding regulatory subunit, is of interest. We demonstrate that *PPP2R2C* expression in prostate cancer cell lines is repressed by forced *GHSROS* overexpression and increased by *GHSROS* knockdown. There is emerging evidence that inactivation of PP2A mediates CRPC in a subset of patients who display resistance to AR-targeting therapies^44, 45^. Loss of *PPP2R2C* expression alone is thought to reprogram prostate tumors towards AR pathway-independent growth and survival^44^. Several independent lines of evidence suggest that *PPP2R2C* is a critical tumor suppressor involved in many cancers. Loss of *PPP2R2C* expression has been attributed to esophageal adenocarcinoma tumorigenesis^57^, and *PPP2R2C* downregulation by distinct microRNAs positively correlates with increased proliferation of cultured cancer cells derived from the prostate^58^, nasopharynx^59^, and ovary^60^. *PPP2R2C* also has a classical growth-inhibiting tumor suppressor role in brain cancers^61^. A subtype of medulloblastoma, pediatric brain tumors, are characterized by high expression of the chemokine receptor CXCR4 and concordant suppression of *PPP2R2C*^62^. Similarly, the gene is ablated in A2B5^+^ glioma stem-like cells, a population which mediates a particularly aggressive chemotherapy-resistant glioblastoma phenotype^63^. Although seemingly paradoxical, *GHSROS* repression of *AR* and *PPP2R2C* in prostate cancer cell lines can be rationalized. Knockdown of *PPP2R2C* using small interfering RNA in cultured LNCaP and VCaP cells did not alter the expression of *AR*^44^. In contrast, *AR* knockdown in androgen-independent LP50 cells^64^ (a cell line derived from LNCaP) markedly decreased *PPP2R2C* expression (Supplementary Fig. S12) – suggestive of an adaptive response to loss of androgen receptor expression (and function). Precisely how *GHSROS* mediates *PPP2R2C* downregulation and its effects on tumor growth remains to be determined, however, *GHSROS* is the first lncRNA shown to downregulate this critical tumor suppressor, suggesting a role in adaptive survival pathways and CRPC development. Taken together, we speculate that *GHSROS* prime prostate tumors for androgen receptor-independent growth.

In this study, the growth of *GHSROS*-overexpressing prostate cancer cell lines was assessed using subcutaneous prostate cancer cell lines xenografts. We appreciate that other models (including orthotopic xenografts) are critical for firmly establishing roles for a gene in cancer processes, including invasion and metastasis^25^, and we will assess these in a future study. The interaction between *GHSROS* and genomic regions, proteins, and other RNA transcripts also requires further elucidation. While this study firmly establishes that *GHSROS* plays a role in prostate cancer, the mechanism by which it reprograms gene expression remains unknown. LncRNAs are now considered critical components of the cellular machinery^1^. Unlike protein-coding genes, which typically require sequence conservation to maintain function, the mechanisms of action of lncRNAs are usually not obvious and uncovering their precise, sometimes subtle, function remains a challenge^1^. For example, some lncRNAs modulate the epigenetic regulation of gene expression and interact with chromatin, acting as scaffolds to guide other molecules (including RNA, proteins, and epigenetic enzymes) to influence gene expression^1, 2^.

Although cancers are highly heterogeneous diseases and few therapies target molecular phenotypes, lncRNAs provide a largely untapped source for new molecular targets^2^. Here, we developed antisense oligonucleotides targeting *GHSROS* and assessed them in cultured cancer cells. We are in the process of refining our oligonucleotides for targeting *in vivo* xenografts. Targeting *GHSROS* may present an opportunity for clinical intervention, however, it is appreciated that translational and regulatory challenges exist for oligonucleotide therapies^65^.

In summary, we propose that *GHSROS* is an oncogene that regulates cancer hallmarks and the expression of a number of genes, including the tumor suppressor *PPP2R2C –* the loss of which is an emerging alternative driver of prostate cancer. Further studies are needed to elucidate the expression and function of *GHSROS* in more detail and to determine whether pharmacological targeting of this lncRNA could prove useful for treating cancer.

## MATERIALS AND METHODS

### Assessment of *GHSROS* transcription in public high-throughput datasets

To expand on Northern blot and qRT-PCR analyses which suggest that the lncRNA *GHSROS* is expressed at low levels^3^, we interrogated ∼4,000 oligonucleotide microarrays with probes for known and predicted exons (Affymetrix GeneChip Exon 1.0 ST). For illustrative purposes, an RNA-sequencing dataset averaging ∼160M reads from metastatic castration-resistant prostate cancer was also examined. See Supplementary information and Supplementary Table S10.

### Cell culture and treatments

Prostate-derived cell lines (PC3, DU145, LNCaP, C4-2B^66^, 22Rv1, DUCaP^67^, RWPE-1, and RWPE-2), the ES-2 ovarian cancer cell line, and the A549 lung cancer cell line were obtained from ATCC (Rockville, MD, USA), except where indicated by a reference. See Supplementary information for details.

### Patient-derived xenografts

Patient-derived xenograft (PDX) lines were obtained in-house (see Supplementary information).

### Production of *GHSROS* overexpressing cancer cell lines

See Supplementary information for details.

### RNA extraction, reverse transcription, and quantitative reverse transcription Polymerase Chain Reaction (qRT-PCR)

See Supplementary information for details. Primers are listed in Supplementary Table S11.

### Locked Nucleic Acid-Antisense Oligonucleotides (LNA-ASO)

Two distinct LNA ASOs complementary to different regions of *GHSROS*, RNV104L and RNV124 (see Supplementary Fig. S6), were designed in-house (by R.N.V.) and synthesized commercially (Exiqon, Vedbæk, Denmark). See Supplementary information for details.

### *In vitro* cell assays

Proliferation and migration assays were performed using an xCELLigence real-time cell analyzer (RTCA) DP instrument (ACEA Biosciences, San Diego, CA). Cell viability was assessed using a WST-1 cell proliferation assay (Roche, Nonnenwald, Penzberg, Germany). See Supplementary information for details.

### Mouse subcutaneous *in vivo* xenograft models

PC3, DU145, and LNCaP cells overexpressing *GHSROS* (or empty vector control) were injected subcutaneously into the flank of 4-week-old male NSG mice (obtained from Animal Resource Centre, Murdoch, WA, Australia). All mouse studies were carried out with approval from the University of Queensland and the Queensland University of Technology Animal Ethics Committees performed in accordance with relevant guidelines and regulations. See Supplementary information for details.

### RNA-sequencing of *GHSROS* overexpressing PC3 and LNCaP cells

See Supplementary information for details. Raw and processed RNA-sequencing (transcriptome) data have been deposited in Gene Expression Omnibus (GEO) with the accession codes GSE86097 (*GHSROS* overexpression in cultured PC3 cells) and GSE103320 (*GHSROS* overexpression in LNCaP xenografts).

### LP50 prostate cancer cell line *AR* knockdown microarray

We interrogated microarray data (NCBI GEO accession no. GSE22483) from androgen-independent late passage LNCaP cells (LP50) subjected to androgen receptor (*AR*) knockdown by shRNA^64^. See Supplementary information for details.

### Survival analysis in clinical gene expression datasets

Non-hierarchical *k*-means clustering was used to partition patients into groups (*k*=2) of samples with similar gene expression patterns. Kaplan-Meier and Cox proportional-hazard model were utilized to generate survival probabilities and hazard ratios (HRs). See Supplementary information for details.

### Code

Code is available in a repository at https://github.com/sciseim/GHSROS_MS.

## ACKNOWLEDGEMENTS

This work was supported by the National Health and Medical Research Council Australia (1002255 and 1059021; to P.L.J., A.C.H., L.K.C., and I.S.), the Cancer Council Queensland (1098565; to A.C.H., R.N.V., L.K.C., and I.S.), the Australian Research Council (grant no DP140100249; to A.C.H., and L.K.C.), a QUT Vice-Chancellor’s Senior Research Fellowship (to I.S.), the Movember Foundation and the Prostate Cancer Foundation of Australia through a Movember Revolutionary Team Award, the Australian Government Department of Health, and the Australian Prostate Cancer Research Center, Queensland (L.K.C., A.C.H., J. H. G., E.D.W., and C.C.N.), Queensland University of Technology, the Instituto de Salud Carlos III (co-funded by European Union ERDF/ESF, “Investing in your future” grant no. PI13-00651; to R.M.L.), a Miguel Servet grant (CP15/00156; to M.D.G.), Junta de Andalucía (grant no. BIO-0139 to R.M.L.), and CIBERobn (CIBER is an initiative of Instituto de Salud Carlos III, Ministerio de Sanidad, Servicios Sociales e Igualdad, Spain; to R.M.L.). We acknowledge the use of the high-performance computational facilities at the Queensland University of Technology and the technical assistance of the Translational Research Institute Histology core and Biological Resource Facility.

## AUTHOR CONTRIBUTIONS

PBT, IS, PLJ and LKC conceived and designed the study, and interpreted the data. PBT, MM, MDG, CW, LJ and PLJ performed laboratory experiments. IS and PBT performed computational biology analyses. PBT, LKC, PLJ and IS wrote the article. All authors (PBT, PLJ, MDG, EJW, CW, MM, LJ, JHG, EDW, CCN, RML, RNV, LKC and IS) contributed to the conception and design of the study, interpretation of the data and writing of the manuscript.

## Supplementary information

**Supplementary Fig. S1.**
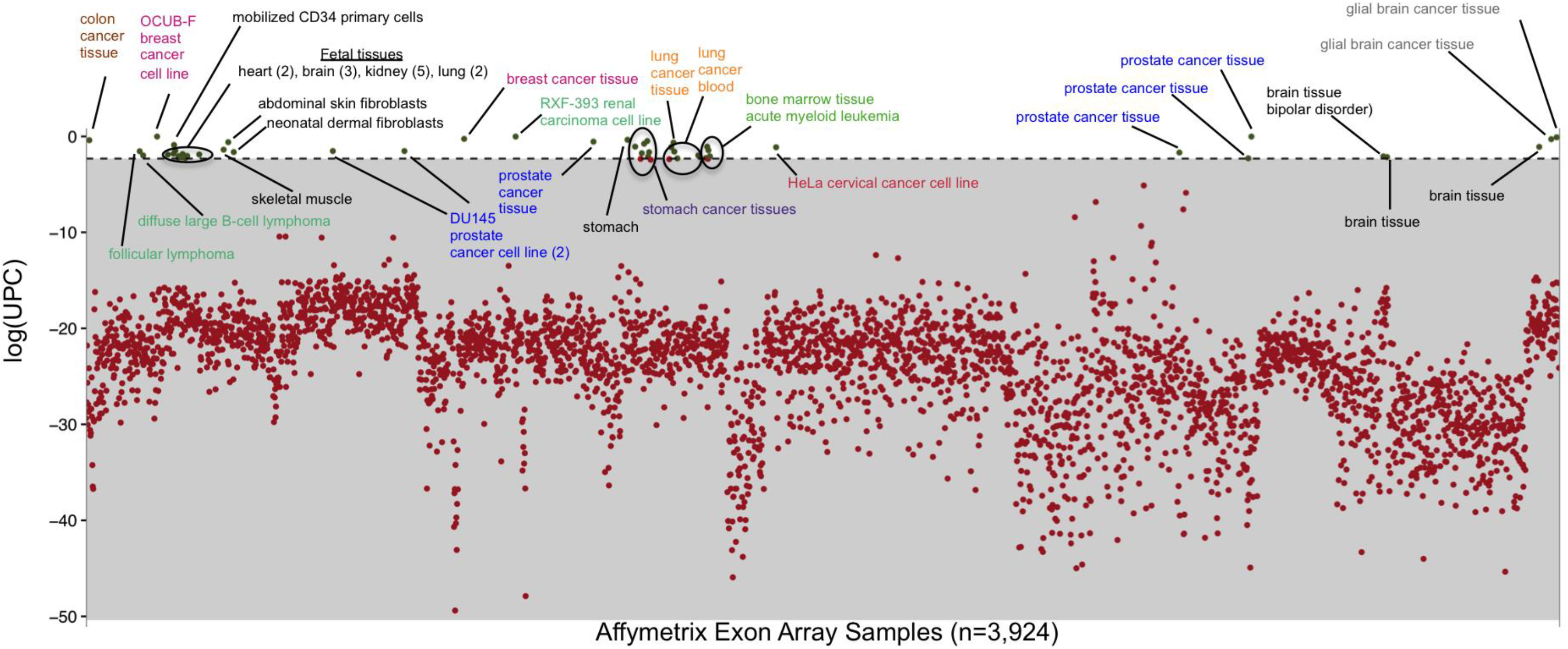
Scatterplot of *GHSROS* Universal exPression Code values in publicly available exon array datasets. The scatter plot shows the log of Universal exPression Code (UPC) values, an estimate on whether a gene is actively transcribed in exon array samples. The dotted horizontal line separates samples with a UPC ≥ 0.1.

**Supplementary Fig. S2.**
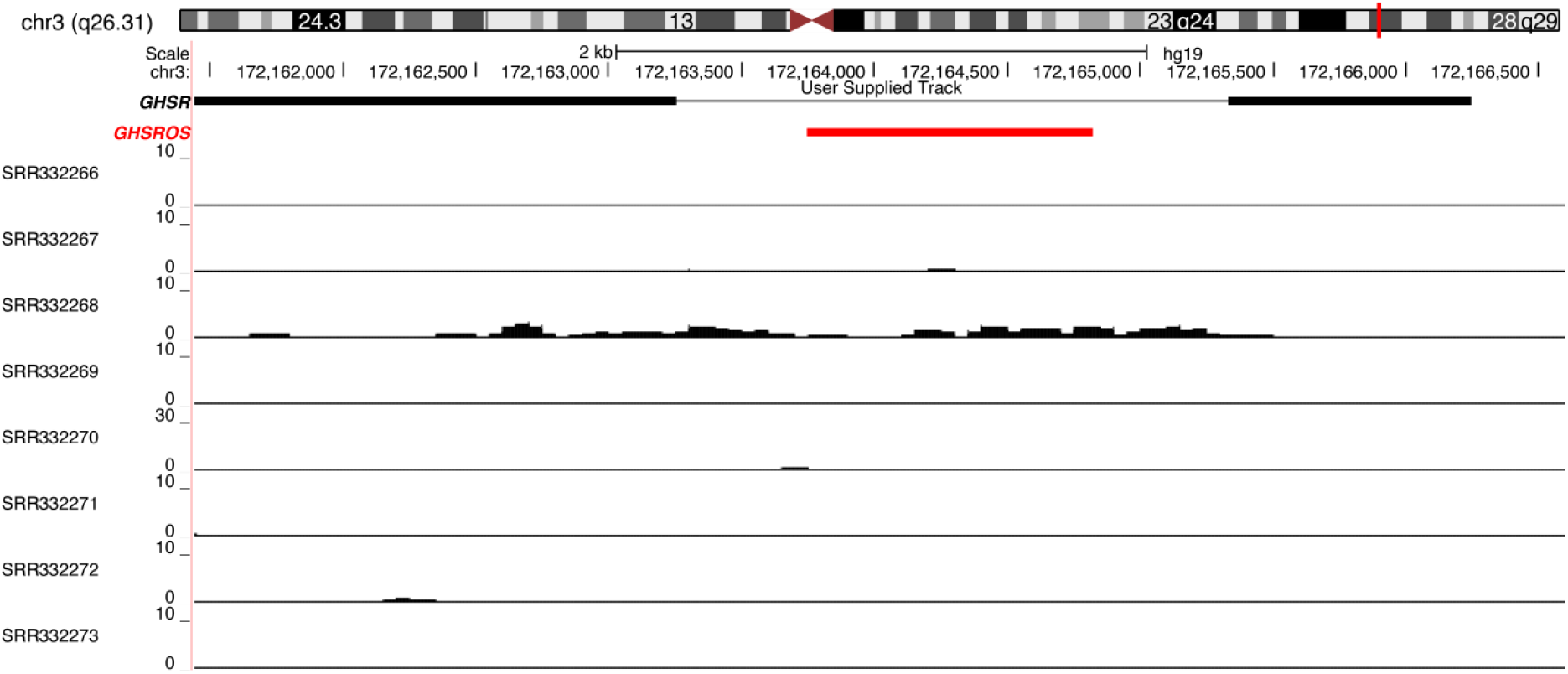
UCSC genome browser visualization of *GHSR*/*GHSROS* locus expression in castration-resistant prostate cancer. *GHSR* exons (black), antisense *GHSROS* exon (red). SRR332266 to SRR332273 denote NCBI Sequence Read Archive (SRA) database accession numbers. The *y*-axis represents read counts normalized to sequencing depth.

**Supplementary Fig. S3.**
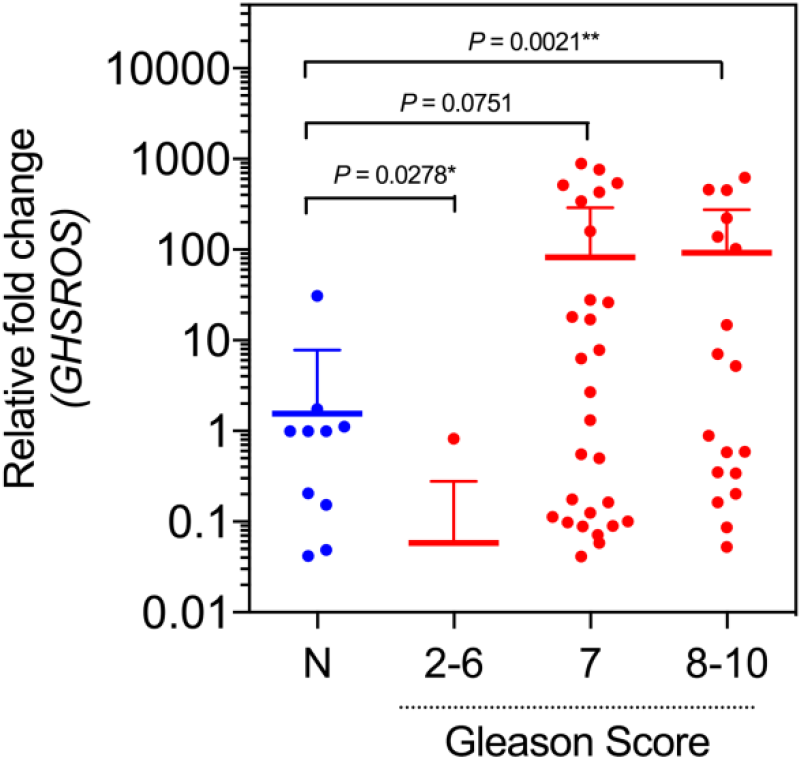
*GHSROS* expression in OriGene TissueScan Prostate Cancer Tissue qPCR panels stratified by Gleason score. \**P* ≤ 0.05, \*\**P* ≤ 0.01, Mann-Whitney-Wilcoxon test. Expression was normalized to the housekeeping gene *RPL32* and relative to a normal prostate sample.

**Supplementary Fig. S4.**
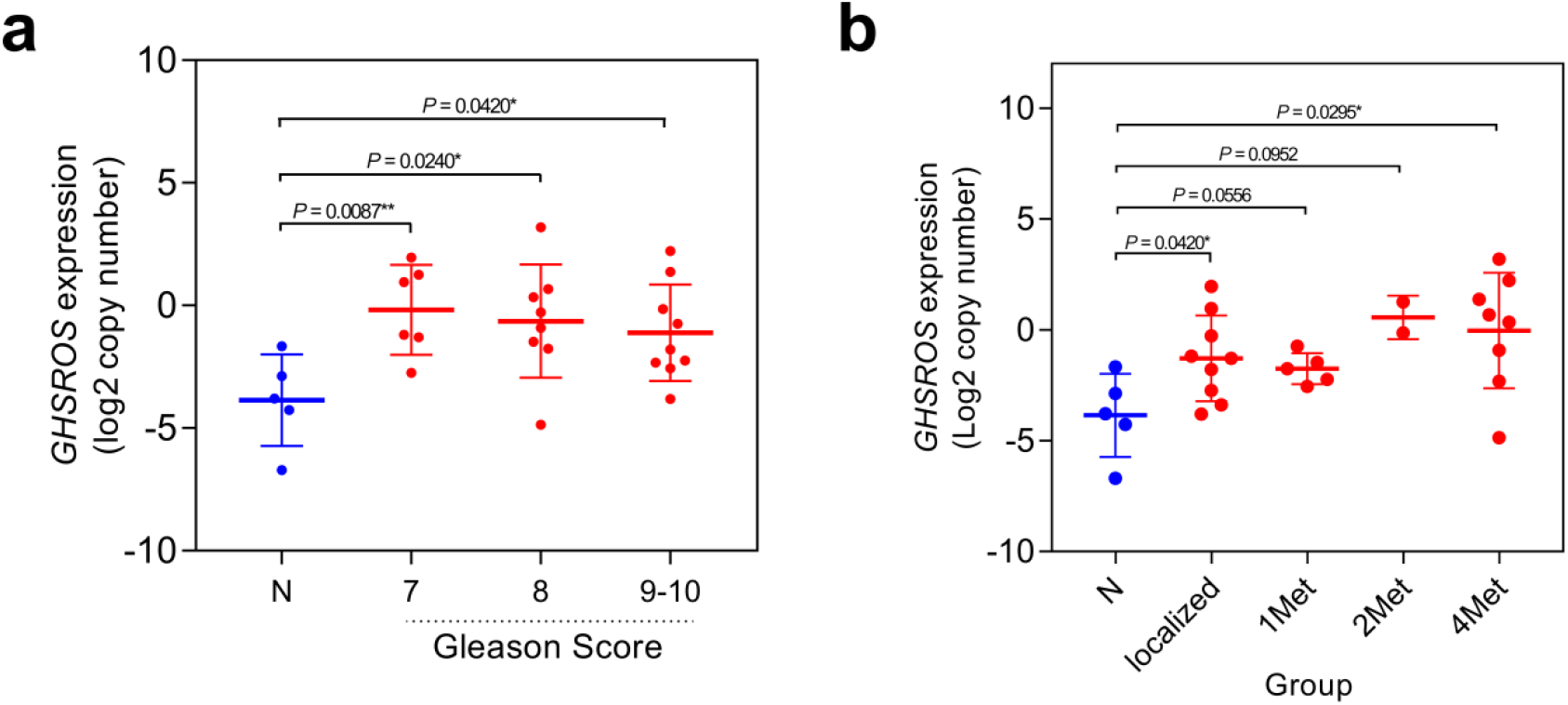
*GHSROS* expression in the Andalusian Biobank prostate tissue cohort. *GHSROS* expression in the Andalusian Biobank prostate tissue cohort stratified by (**a)** Gleason score and **(b)** number of metastatic sites. 1 Met denotes one and ≥ 2 Met two or more metastatic sites. Absolute expression levels were determined by qRT-PCR and adjusted by a normalization factor calculated from the expression levels of three housekeeping genes (*HPRT*, *ACTB*, and *GAPDH*). N denotes normal prostate. \**P* ≤ 0.05, \*\**P* ≤ 0.01, Mann-Whitney-Wilcoxon test.

**Supplementary Fig. S5.**
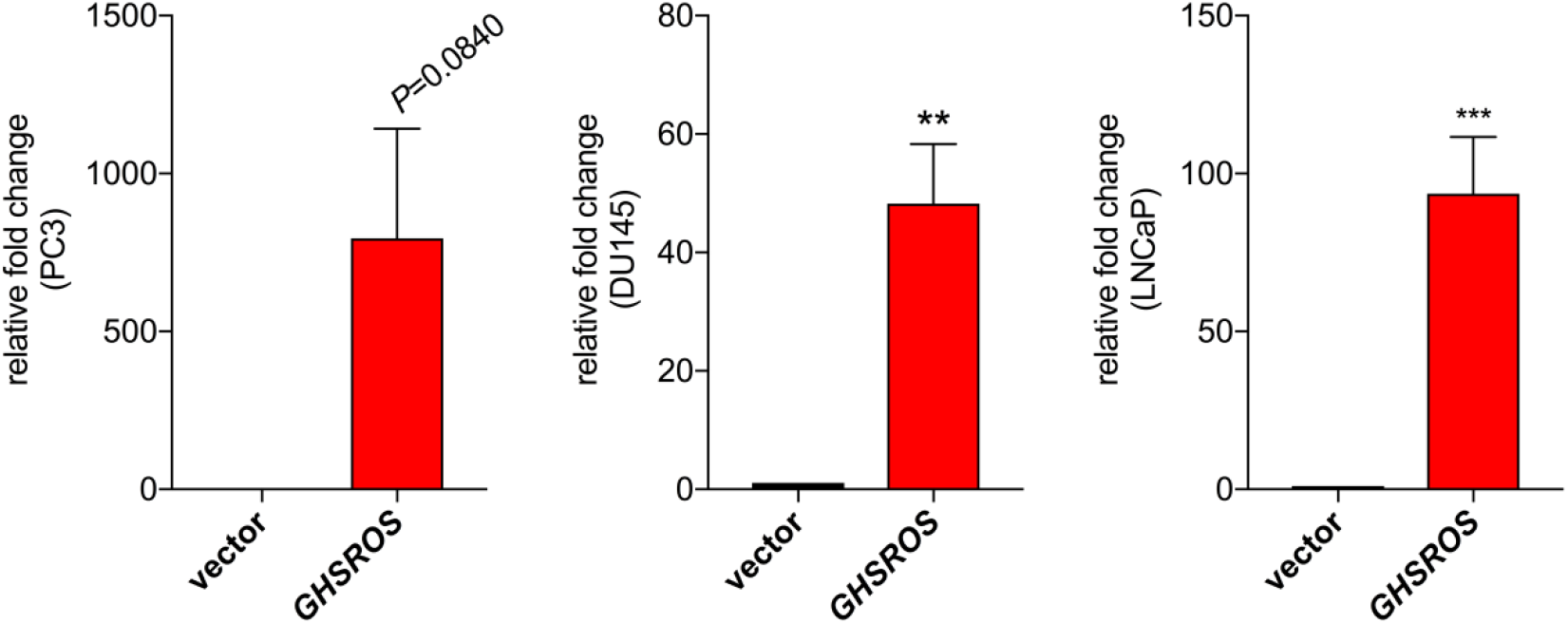
Confirmation of *GHSROS* overexpression in prostate cancer-derived cell lines. Bar graphs show qRT-PCR quantification of the relative expression levels of *GHSROS* when overexpressed in prostate-derived (PC3, DU145, and LNCaP) cancer cell lines. Expression was normalized to the housekeeping gene *RPL32* using the comparative 2^-ΔΔCt^ method of quantification. Results are relative to the respective vector control. Mean ± s.e.m., *n*=3, \*\**P* ≤ 0.01, \*\*\**P* ≤ 0.001, Student’s *t*-test.

**Supplementary Fig. S6.**
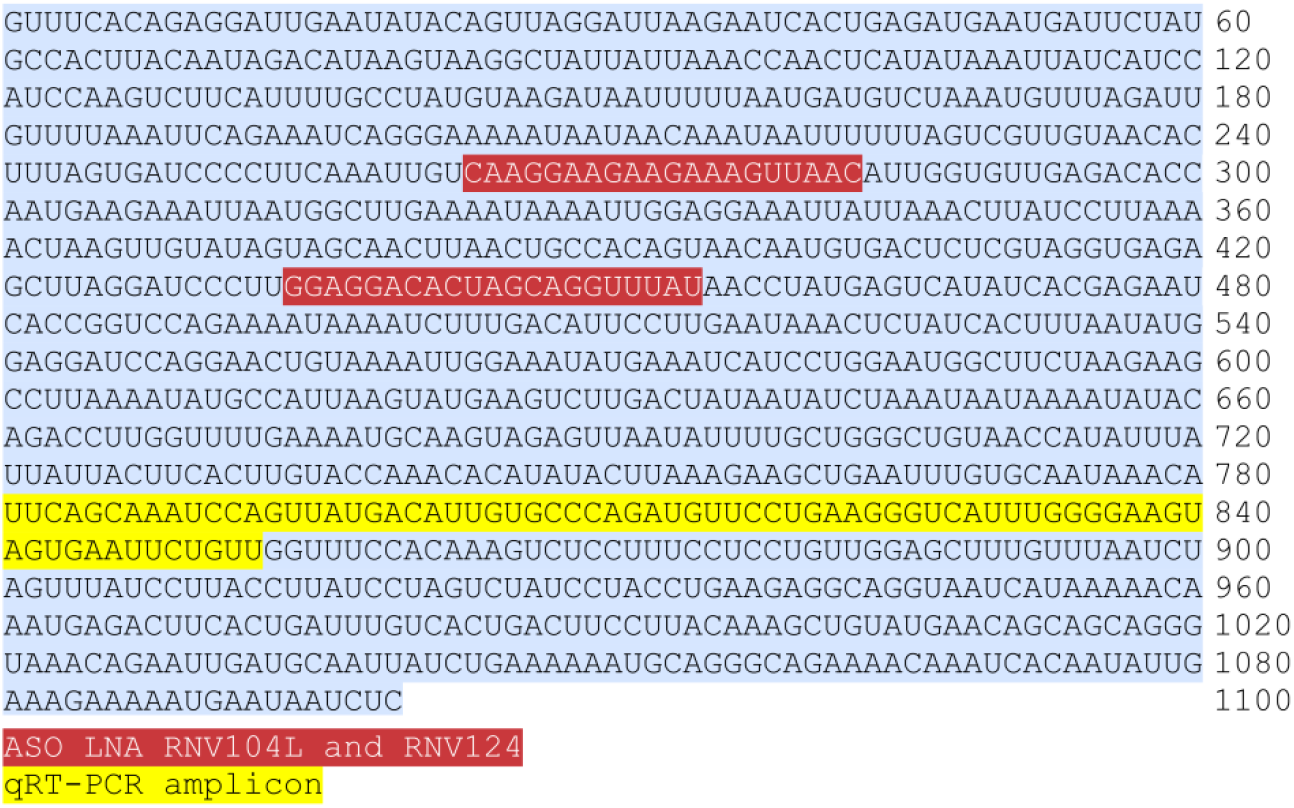
Sequence of the lncRNA *GHSROS* and regions targeted by oligonucleotides. The 1.1kb *GHSROS* sequence. Locked nucleic antisense oligonucleotide (LNA-ASO) locations are highlighted in red and the qRT-PCR amplicon in yellow.

**Supplementary Fig. S7.**
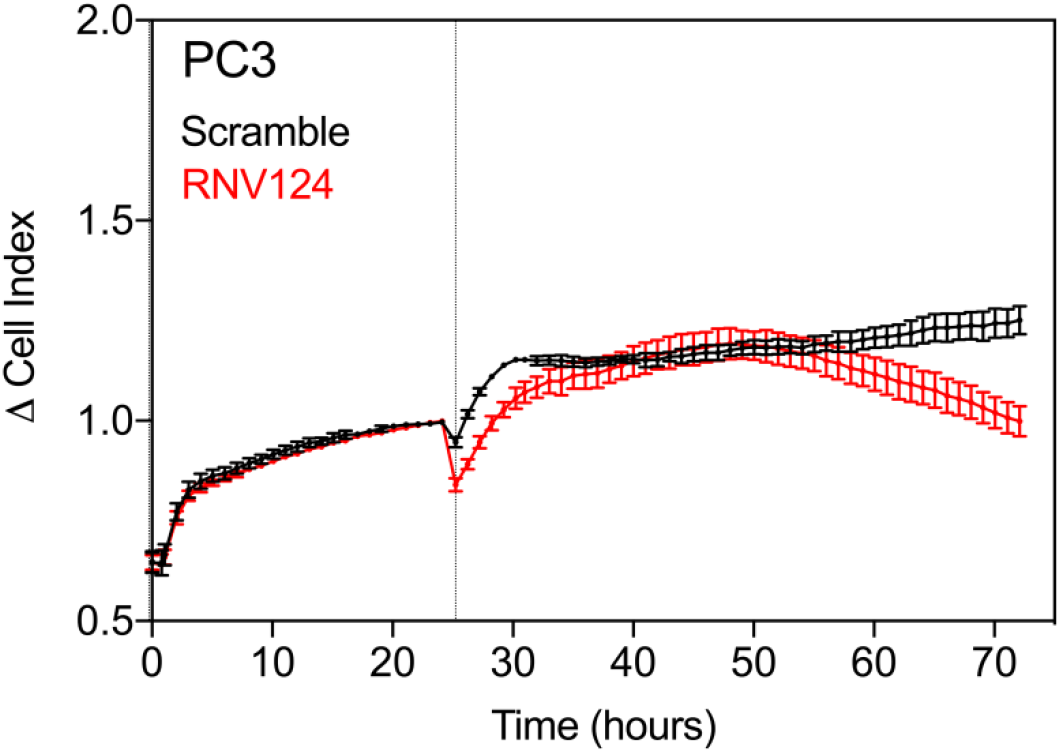
*GHSROS* knockdown attenuates PC3 cell survival upon serum starvation. Cells were transfected with the LNA-ASO RNV124 and grown for 24 hours (indicated by a vertical dotted line) prior to serum starvation. Results are relative to scrambled control. Mean ± s.e.m., *n*=3. At 48 hours after serum starvation, (72 hours after cells transfected as indicated on the x-axis), there was a significant difference in survival (*P* = 0.049, Student’s *t*-test).

**Supplementary Fig. S8.**
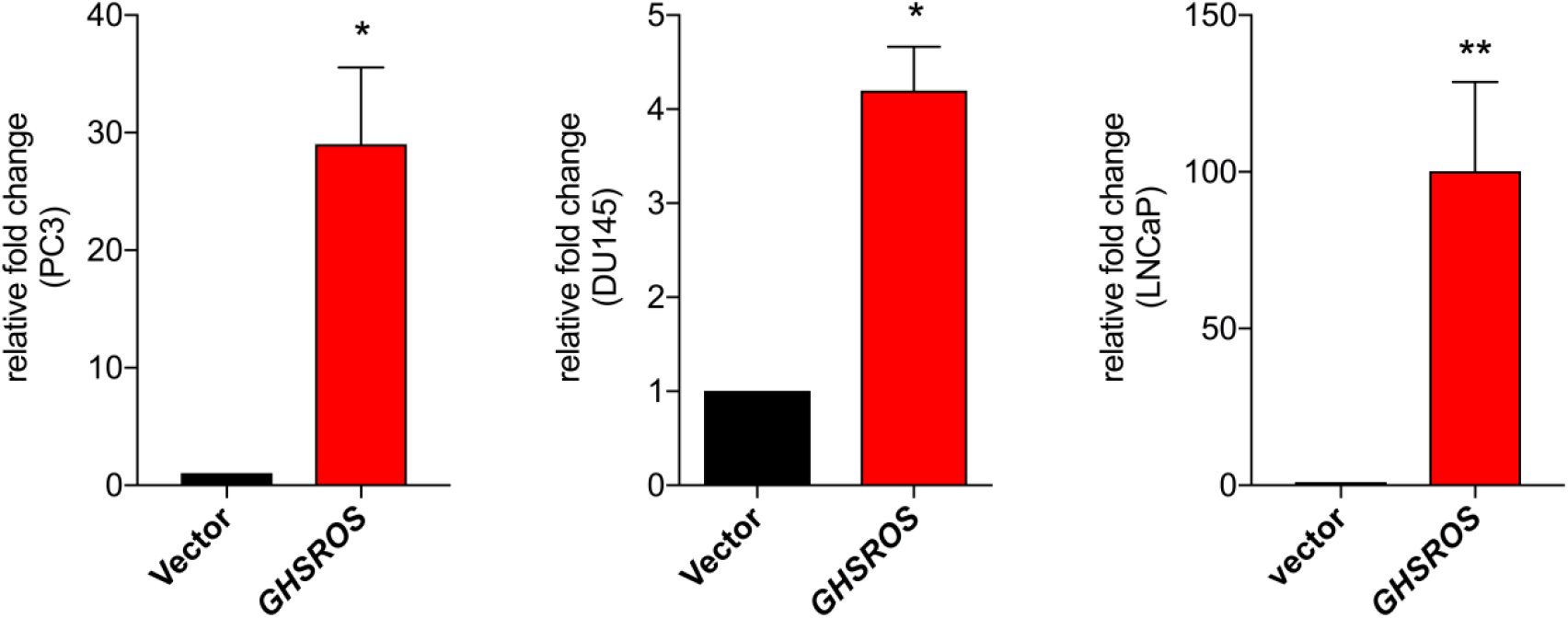
Validation of *GHSROS* overexpression in PC3, DU145, and LNCaP tumor xenografts by qRT-PCR. Expression changes were measured from excised PC3 (*n*=2 vector, *n*=3 *GHSROS*), DU145 (*n*=2 vector, *n*=3 *GHSROS*), and LNCaP xenografts (*n*=8 vector, *n*=5 *GHSROS*) at *in vivo* endpoint. Expression was normalized to the housekeeping gene *RPL32*. Results are relative to the respective vector control. Mean ± s.e.m., *n*=3, \**P* ≤ 0.05, \*\**P* ≤ 0.01, Student’s *t*-test.

**Supplementary Fig. S9.**
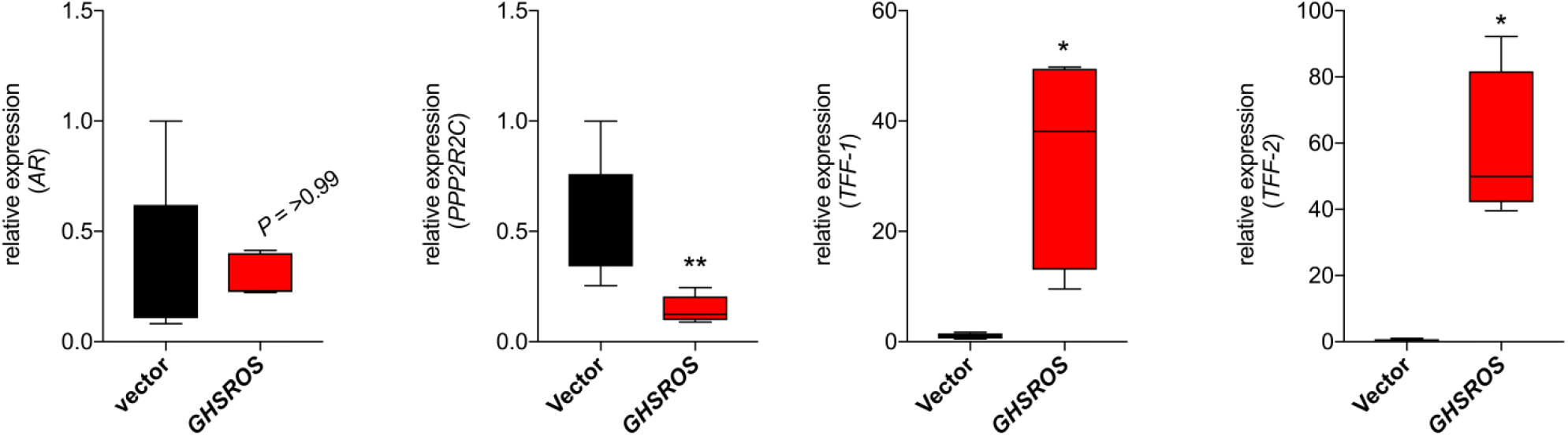
Expression of selected genes in LNCaP tumor xenografts overexpressing *GHSROS*. Expression changes were measured by qRT-PCR from excised LNCaP (*n*=4-8 vector, *n*=4-5 *GHSROS*) at *in vivo* endpoint. Expression was normalized to the housekeeping gene *RPL32*. Results are relative to the respective vector control. Mean, *n*=3, \**P* ≤ 0.05, \*\*\**P* ≤ 0.01, Student’s *t*-test.

**Supplementary Fig. S10.**
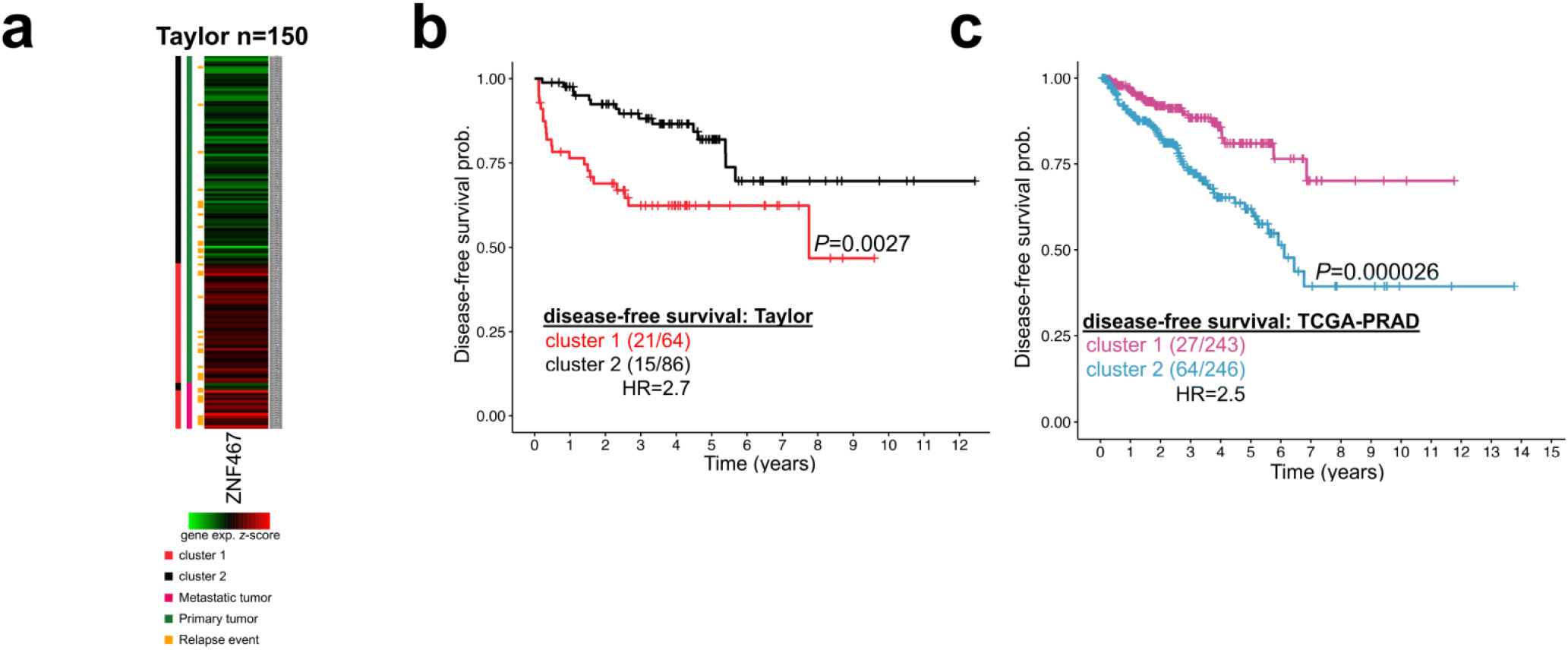
Zinc finger protein 467 (*ZNF467*), a gene induced by forced *GHSROS*-overexpression, is upregulated by metastatic tumors and associated with adverse relapse outcome. **(a)** Heat map of *ZNF467* expression in the Taylor cohort normalized to depict relative values within rows (samples) with high (red) and low expression (green). Vertical bars show patient grouping by *k*-means clustering (cluster 1, red; cluster 2, black), tumor type (primary, green; metastatic pink), and relapse status (relapse event, orange). **(b)** Kaplan-Meier analyses of *ZNF467* in the Taylor cohort. Patients were stratified by *k*-means clustering, as described in (a). **(c)** Kaplan-Meier analyses of *ZNF467* in the TCGA-PRAD cohort of 489 localized prostate tumors. Patients were stratified by *k*-means clustering (cluster 1, purple; cluster 2, turquoise).

**Supplementary Fig. S11.**
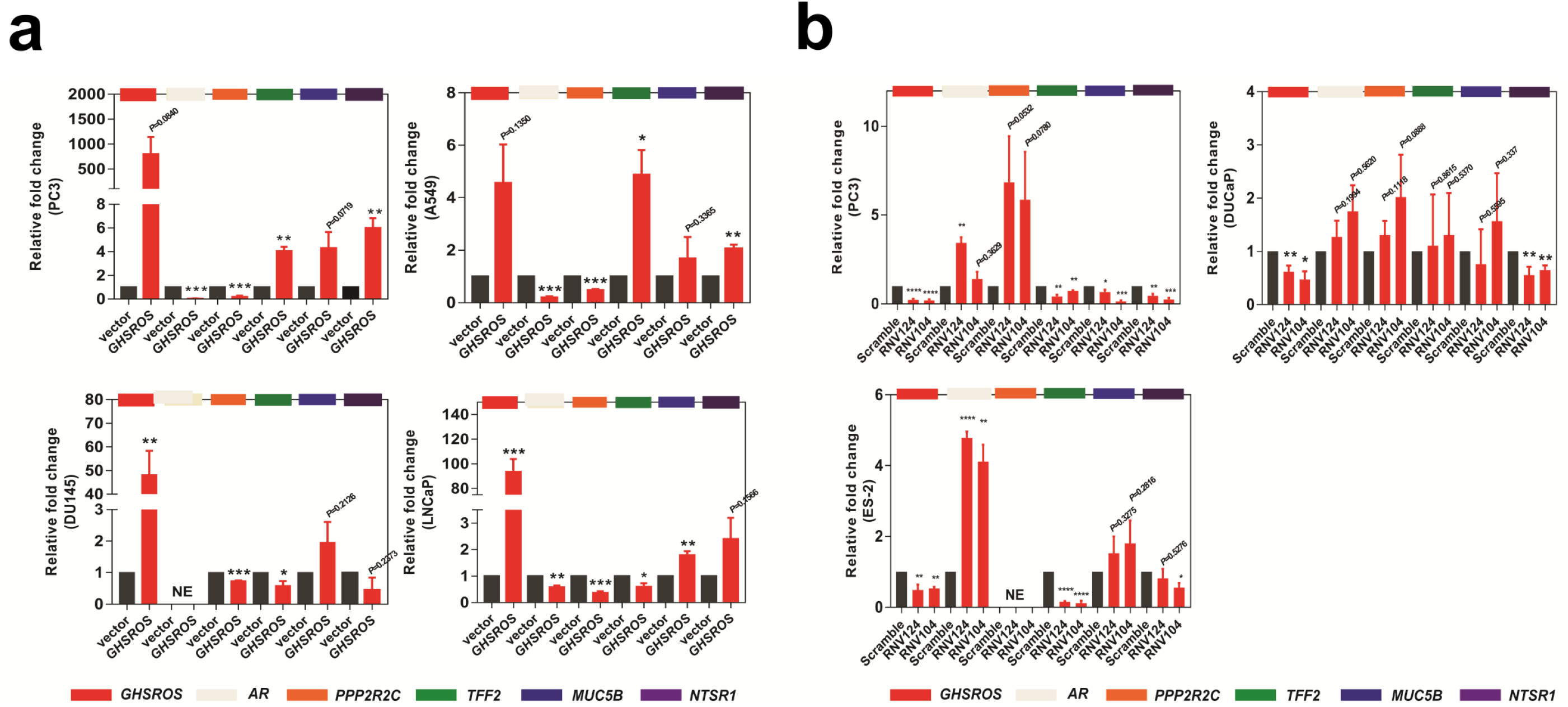
Effects of *GHSROS* perturbation in cultured cells assessed by qRT-PCR. **(a)** qRT-PCR validation of 5 genes regulated by *GHSROS*. Expression was normalized to the housekeeping gene RPL32. Results are relative to the respective vector control. Coloured bars indicate individual genes. Genes that were not expressed represented as no expression (NE). Mean ± s.e.m., *n*=3, \**P* ≤ 0.05, \*\**P* ≤ 0.01, \*\*\**P* ≤ 0.001, Student’s *t*-test. **(b)** qRT-PCR validation of regulated genes following knockdown of *GHSROS* by transfection with LNA-ASOs for 48 hours. Expression was normalized to the housekeeping gene *RPL32*. Results are relative to the respective scrambled control. Annotated as in (a).

**Supplementary Fig. S12.**
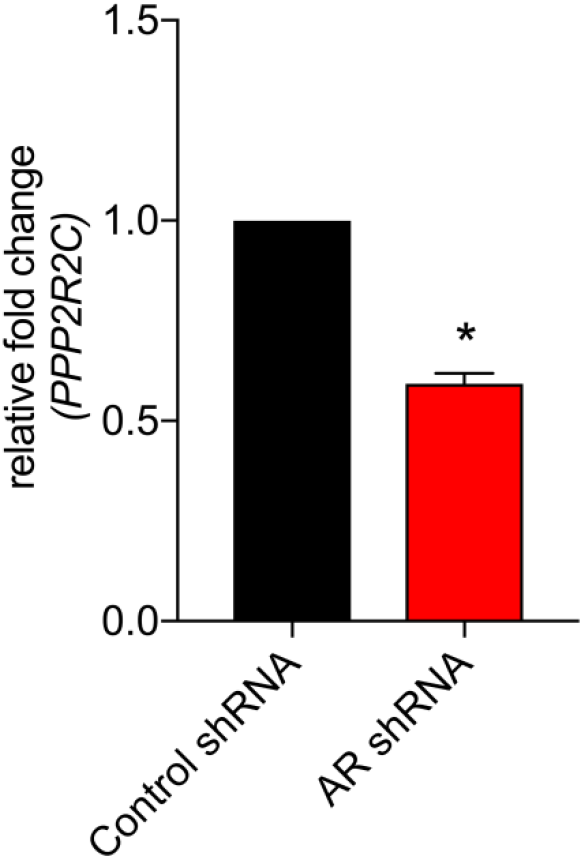
Effect of androgen receptor (AR) perturbation in LP50 prostate cancer cells on *PPP2R2C* expression. Assessed by microarray (NCBI GEO accession no. GSE22483). Mean ± s.e.m. *n*=2, \**Q* ≤ 0.05, moderated *t*-test.

**Supplementary Table S1.** Correlation between *GHSROS* expression and clinicopathological parameters in OriGene TissueScan Prostate Cancer Tissue qPCR panels. Six samples were excluded due to missing clinical information. Relative *GHSROS* expression in tumors (T) stratified by clinical stage and Gleason score was compared to a normal prostate sample (N). *P*-values were calculated using the Mann-Whitney-Wilcoxon test. NA = not applicable, PD = other prostatic diseases.

| Clinicopathological parameters | Sample number (*n*) | *P*-value |
| --- | --- | --- |
| Age at diagnosis (mean ± SD) | 62.2 ± 7.80 | |
| N/ T | 24/88 | 0.1413 |
| PD/ T | 31/88 | 0.8001 |
| N/ PD | 24/31 | 0.0691 |
| **Clinical stage** |  |  |
| N (normal prostate) | 24 |  |
| I | 0 | NA |
| II | 47 | 0.311 |
| III | 33 | 0.0855 |
| IV | 3 | 0.0185 |
| **Gleason score** |  |  |
| N (normal prostate) | 24 |  |
| 2-6 | 15 | 0.0278 |
| 7 | 47 | 0.0751 |
| 8-10 | 25 | 0.00210 |

**Supplementary Table S2.** Correlation between *GHSROS* expression and clinicopathological parameters in the Andalusian Biobank prostate tissue cohort. Absolute levels of *GHSROS* expression in tumors were stratified by Gleason score and the number of metastatic tumor sites were compared to normal prostate (N). Tumors positive or negative for extraprostatic extension and perineural infiltration were compared to each other. *P*-values were calculated using the Mann-Whitney-Wilcoxon test. NA = not applicable.

| Clinicopathological parameters | Sample number (*n*) | *P*-value |
| --- | --- | --- |
| **Age at diagnosis (mean ± SD)** | 73.7 ± 9.81 |  |
| **Gleason score** |  |  |
| N (normal prostate) | 8 |  |
| 7 | 6 | 0.00870 |
| 8 | 9 | 0.0240 |
| 9-10 | 13 | 0.0420 |
| **Number of metastatic sites** |  |  |
| N (normal prostate) | 8 |  |
| primary prostate tumor: 0/localized | 10 | 0.0420 |
| primary prostate tumor: 1 metastatic site | 6 | 0.0556 |
| primary prostate tumor: ≥2 metastatic sites | 12 | 0.0127 |
| **Extraprostatic extension** |  |  |
| - | 16 | 0.379 |
| + | 11 |  |
| **Perineural infiltration** |  |  |
| - | 8 | 0.415 |
| + | 20 |  |

**Supplementary Table S3.**
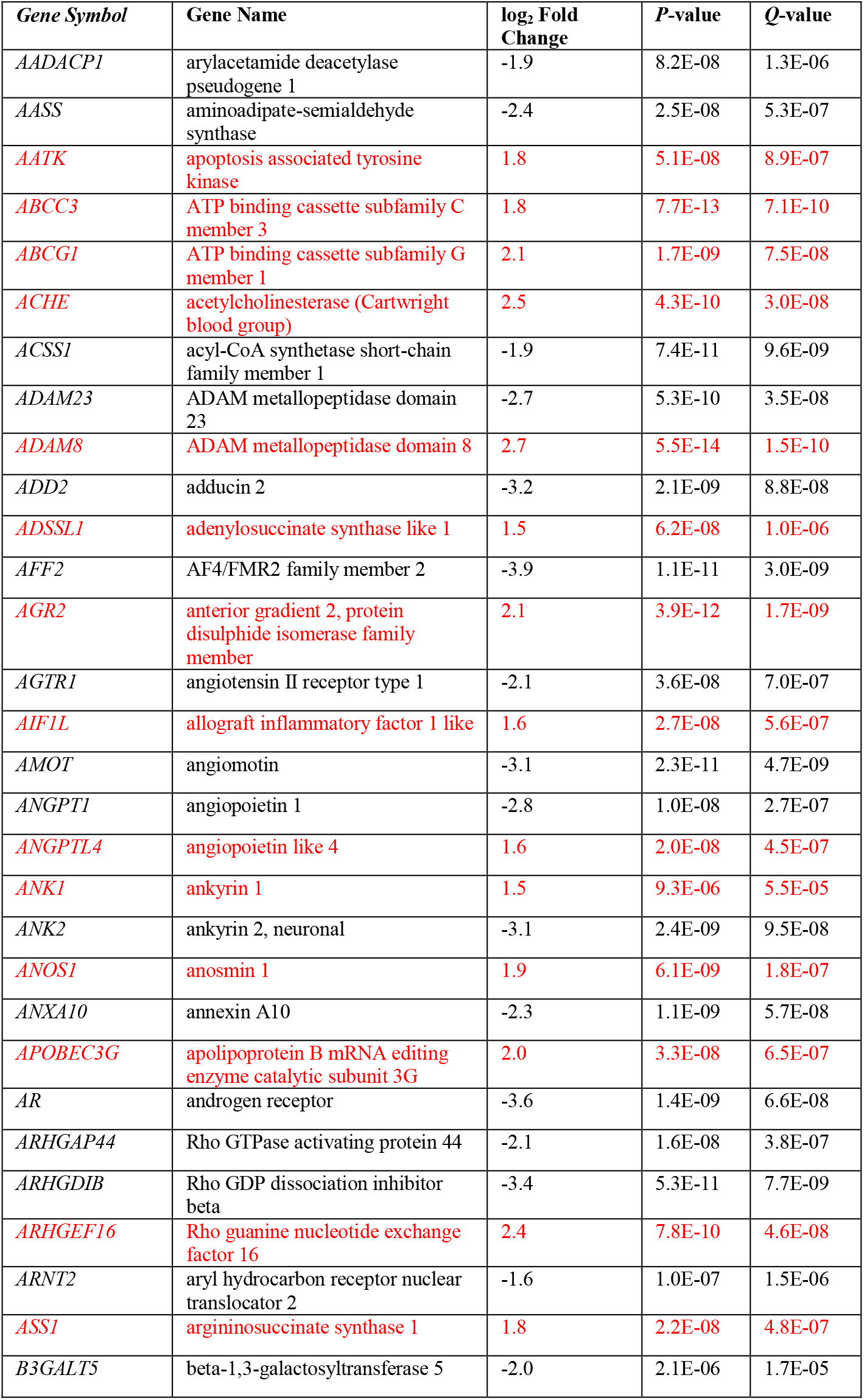
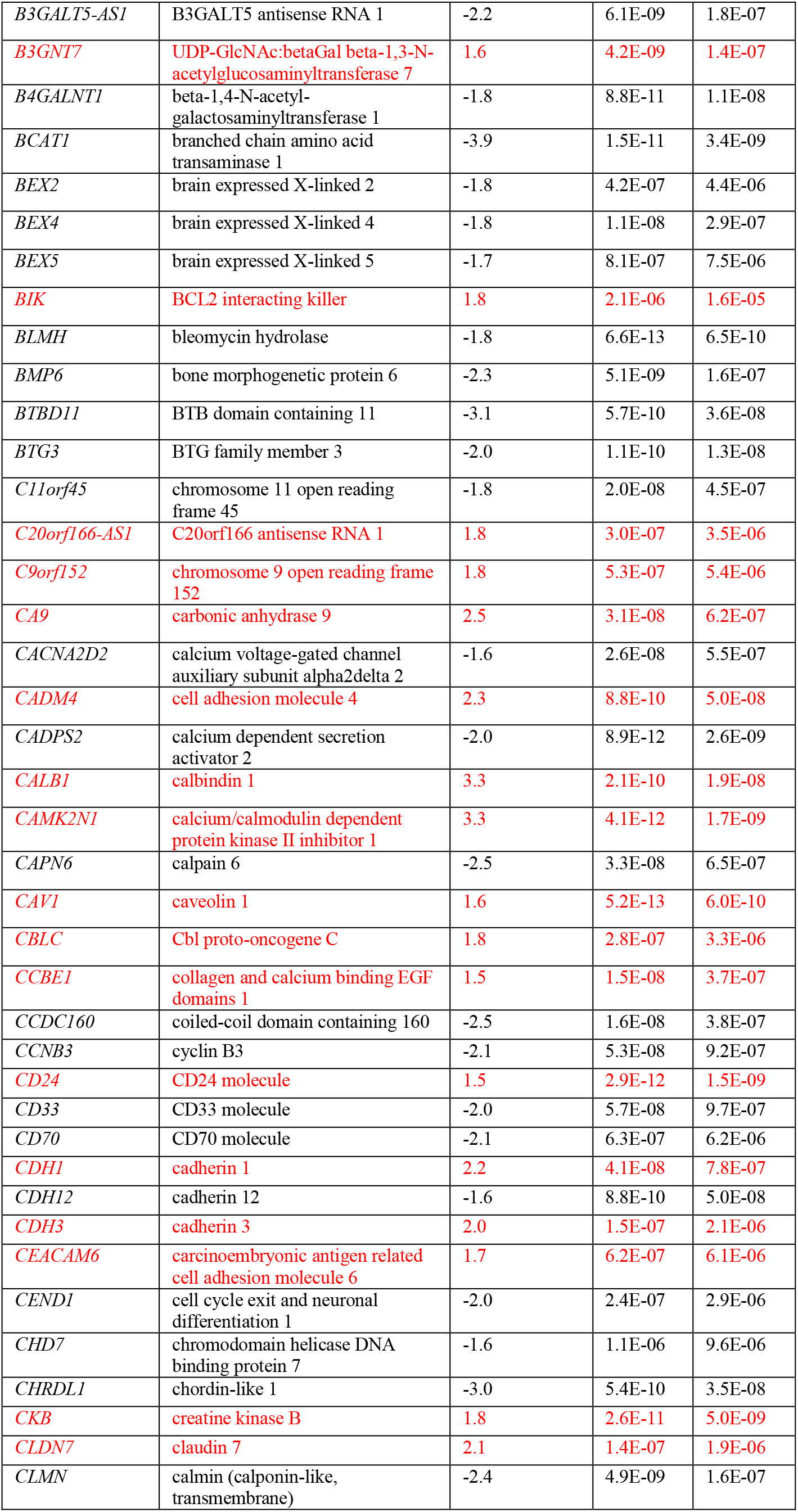
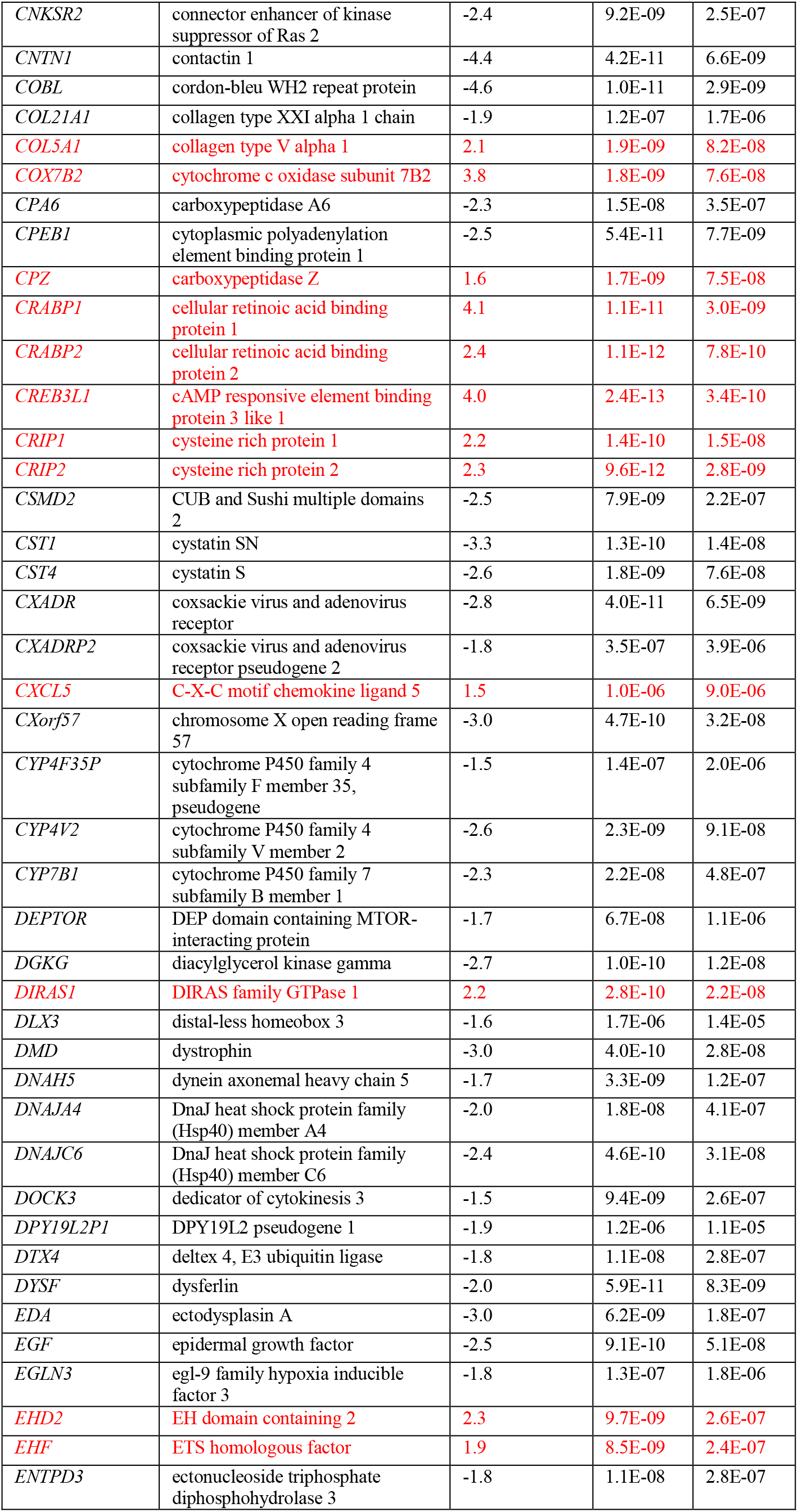
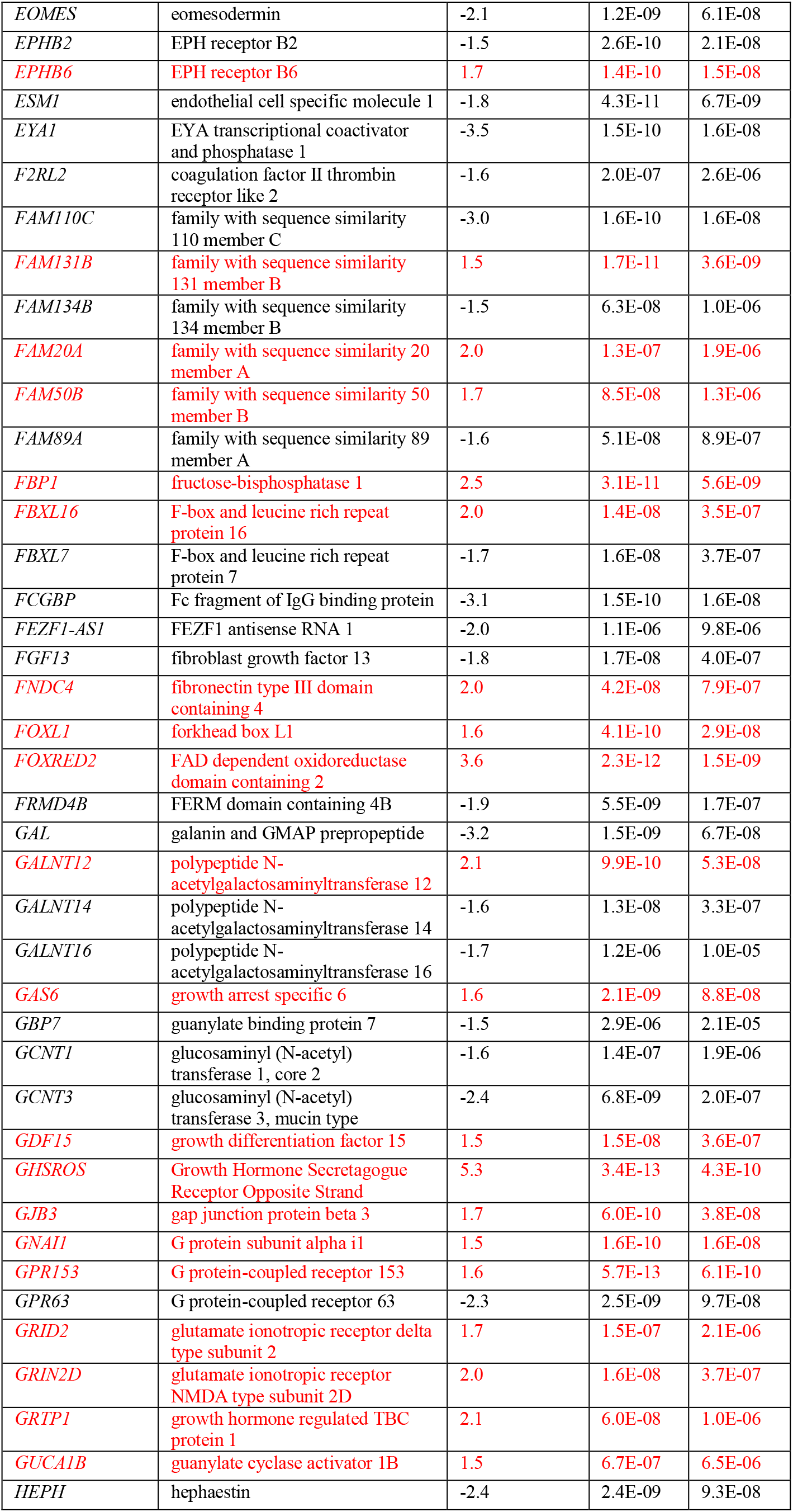
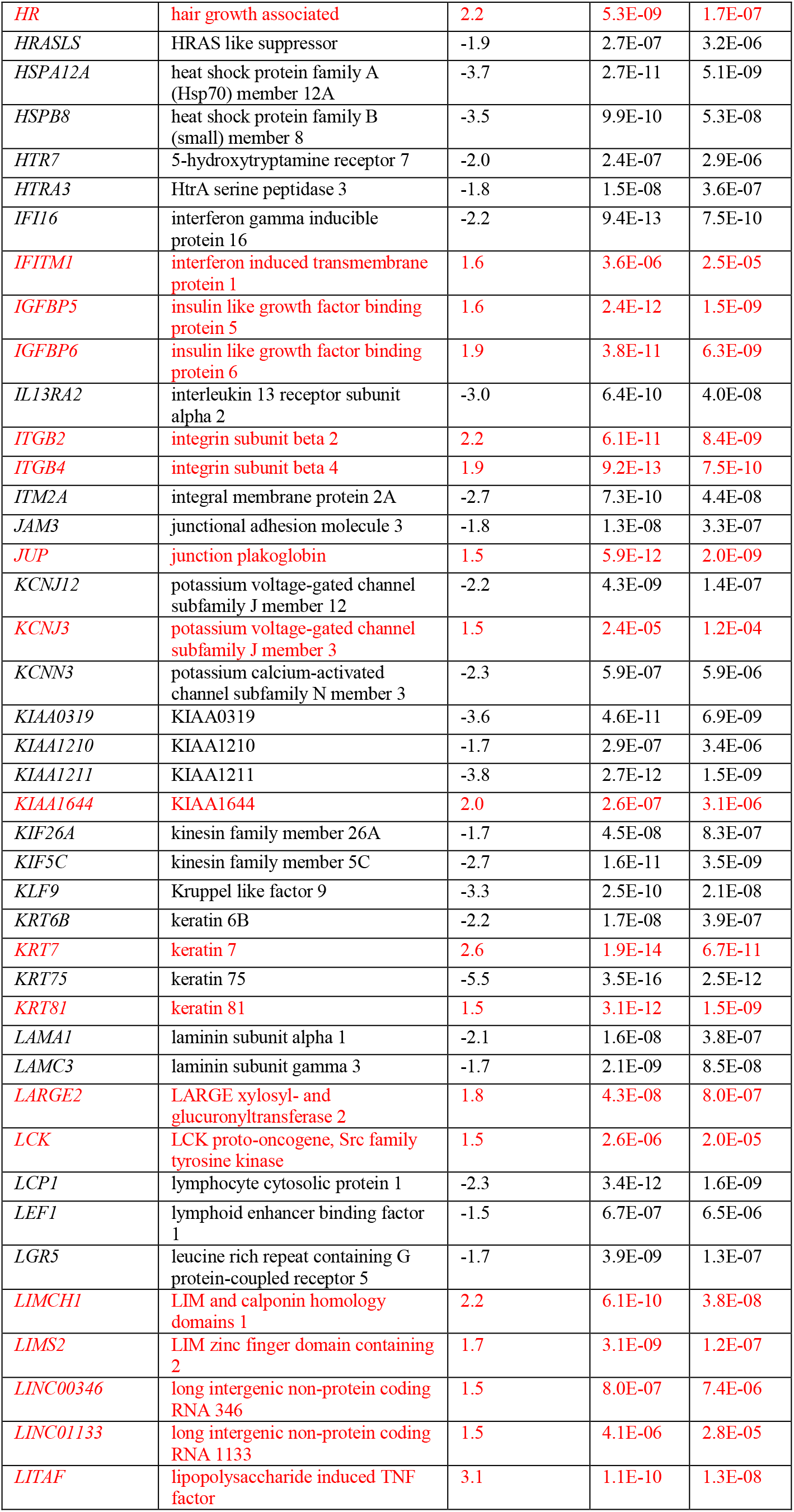
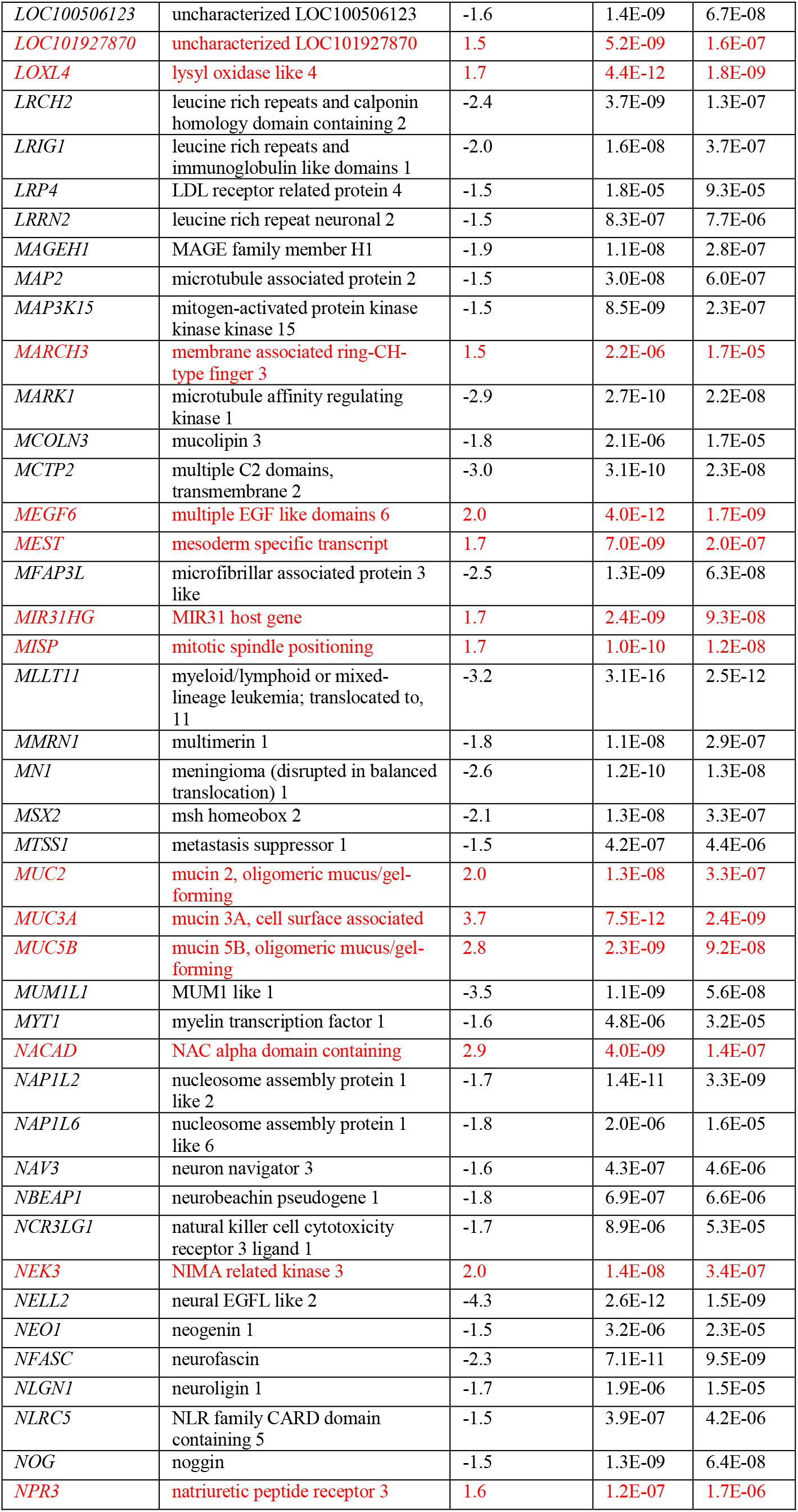
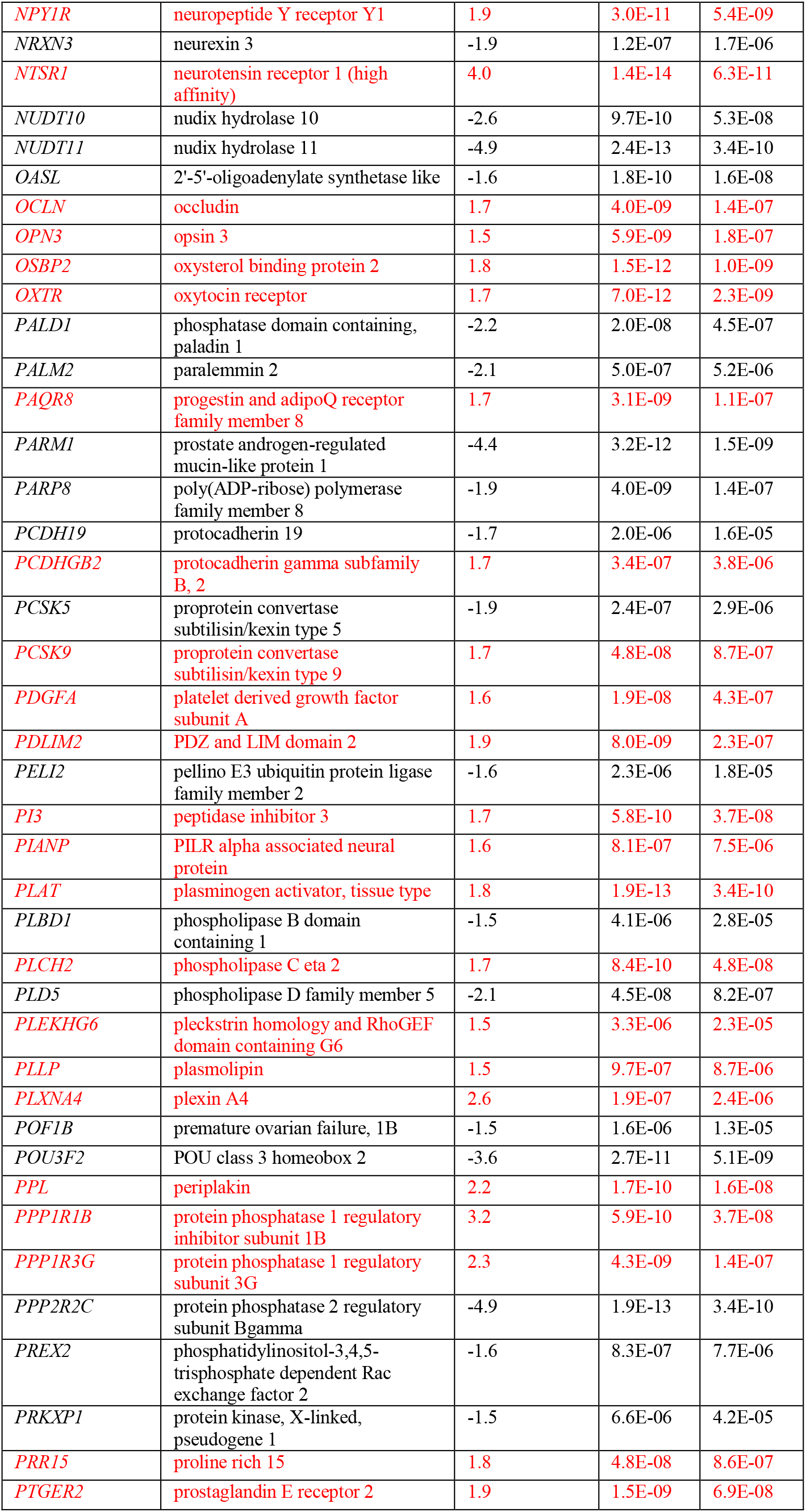
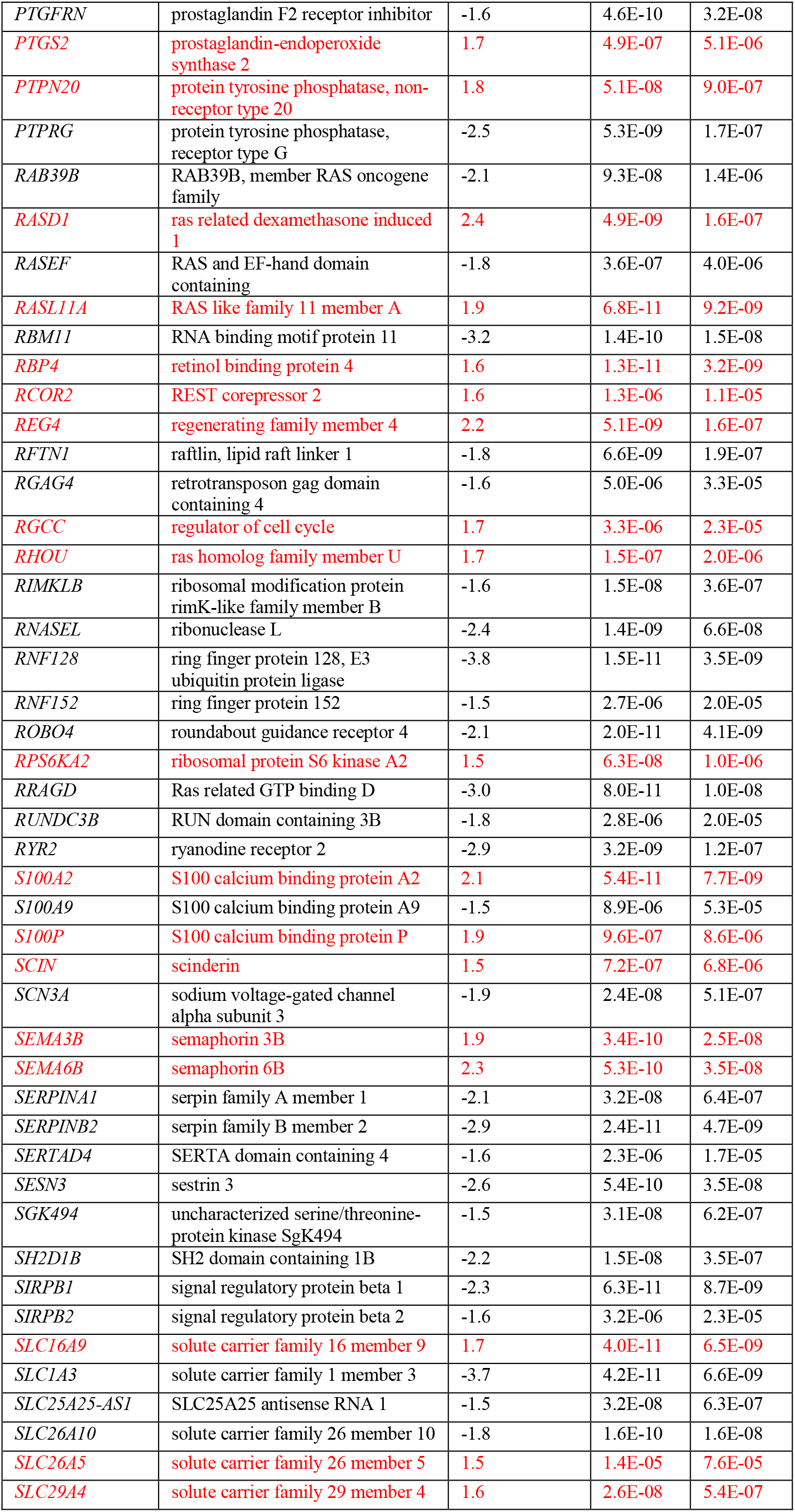
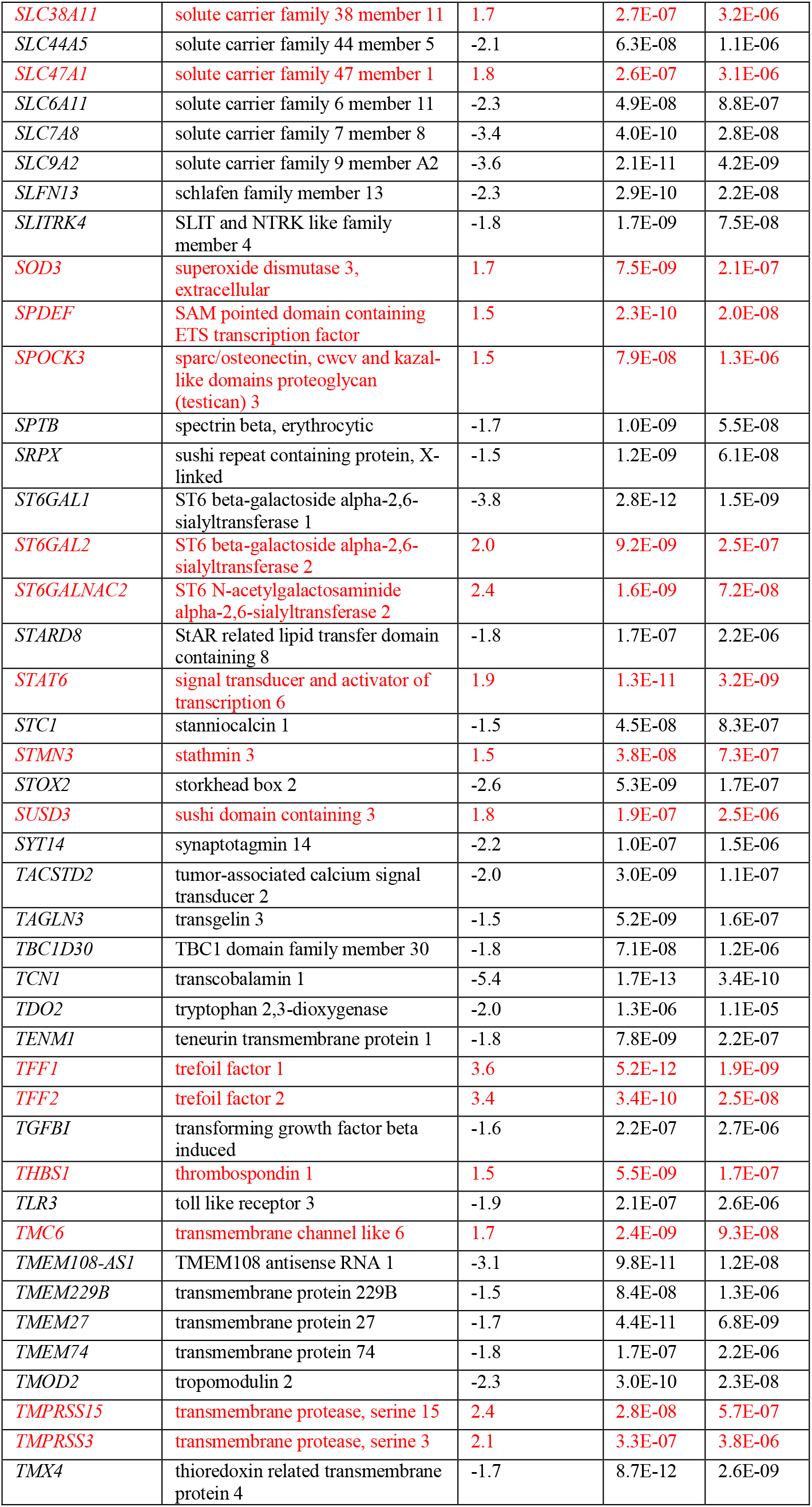
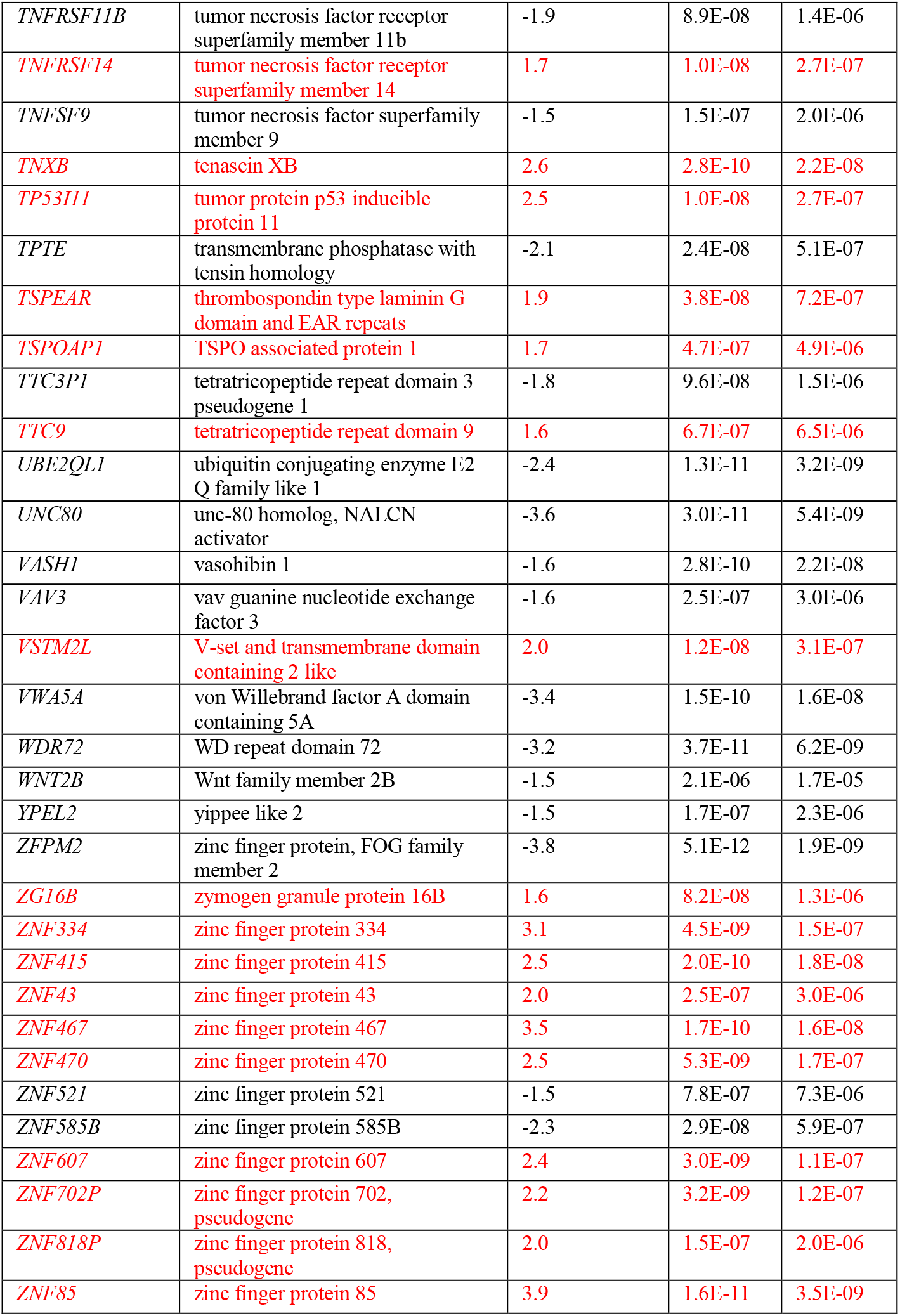
Differentially expressed genes in PC3-GHSROS cells. compared to empty vector control. Red: higher expression in PC3-GHSROS cells; Black: lower expression in PC3-GHSROS cells. Fold-changes are log_2_ transformed; *Q*-value denotes the false discovery rate (FDR; Benjamini-Hochberg)-adjusted *P*-value (cutoff ≤ 0.05).

**Supplementary Table S4.**
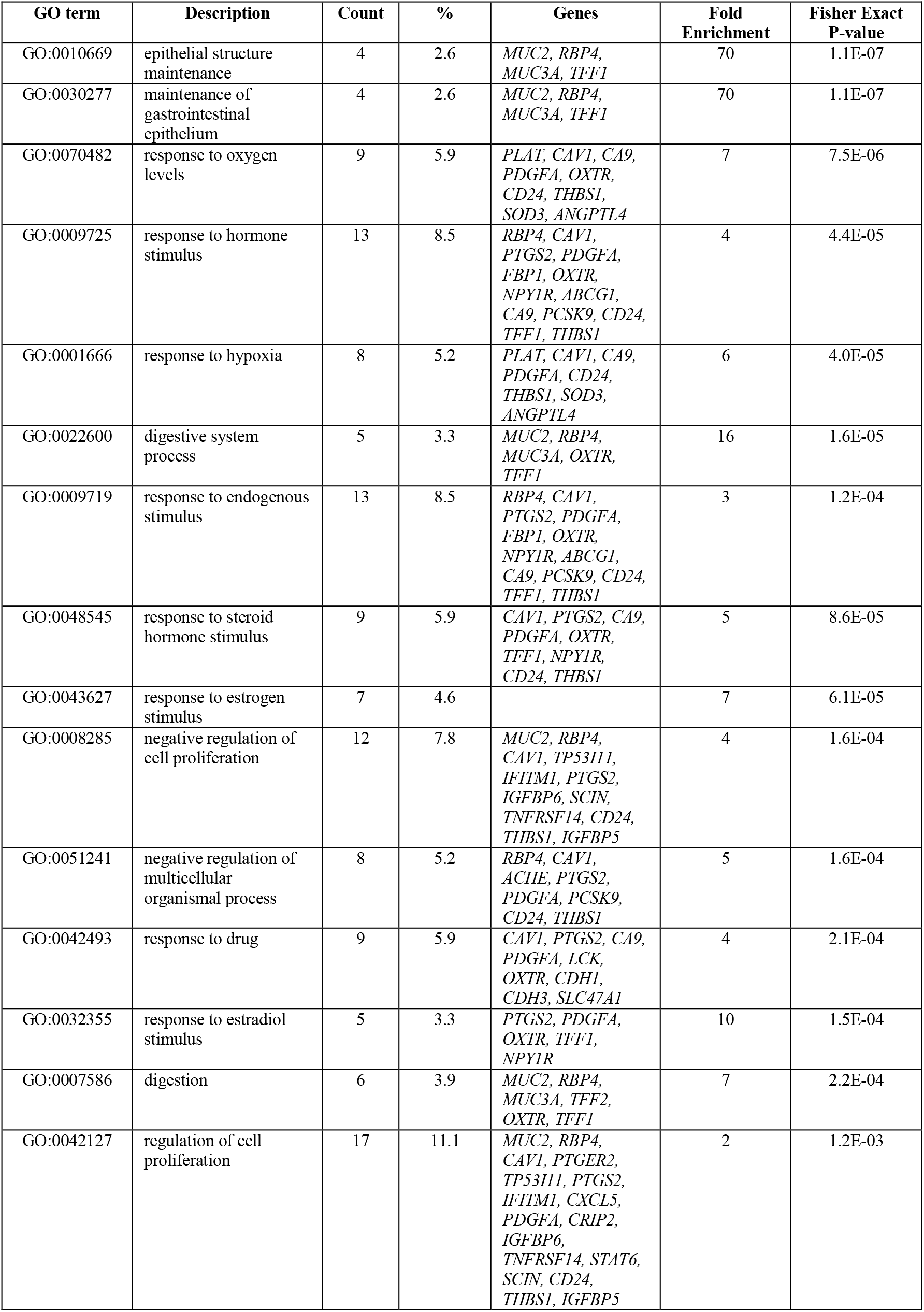
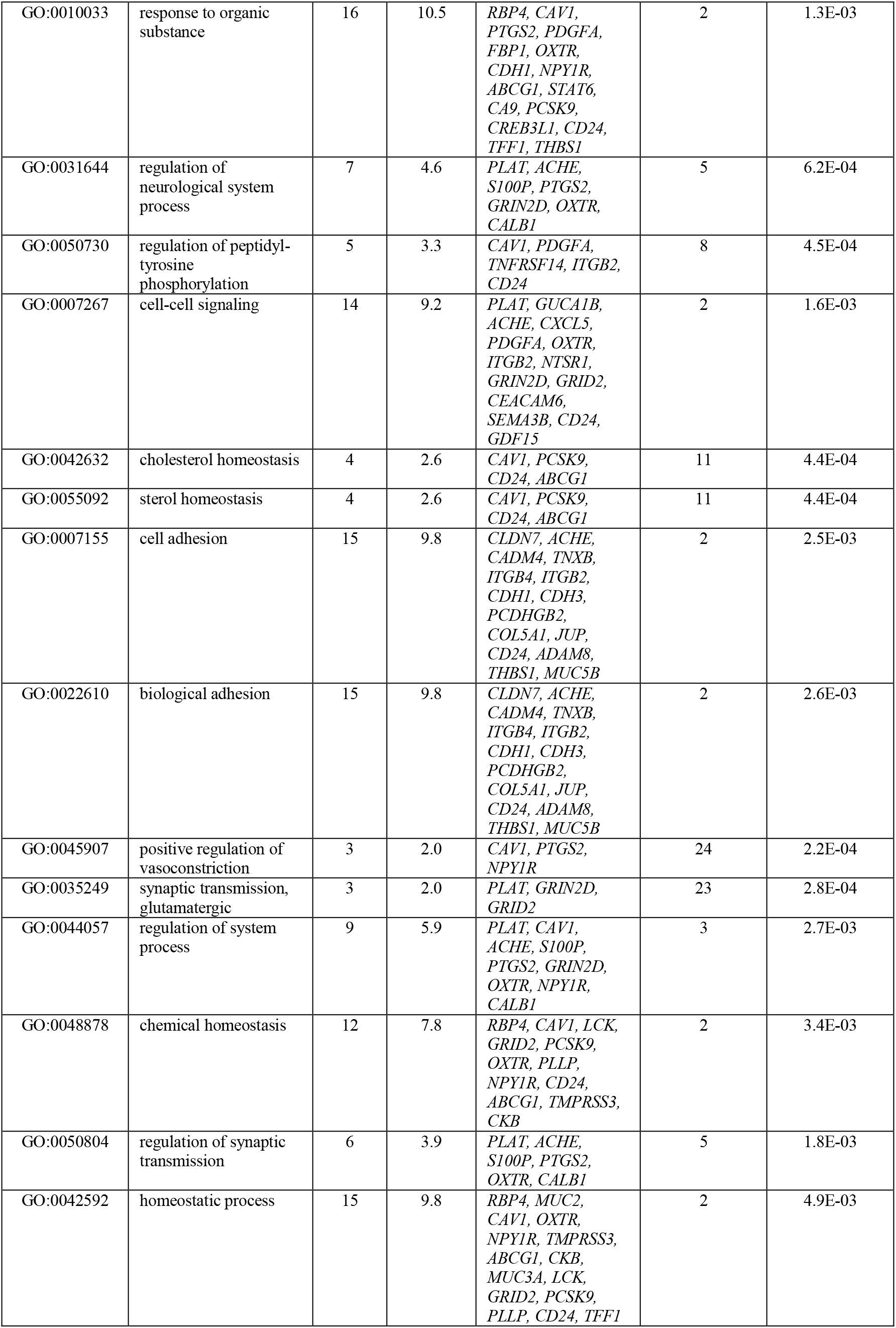
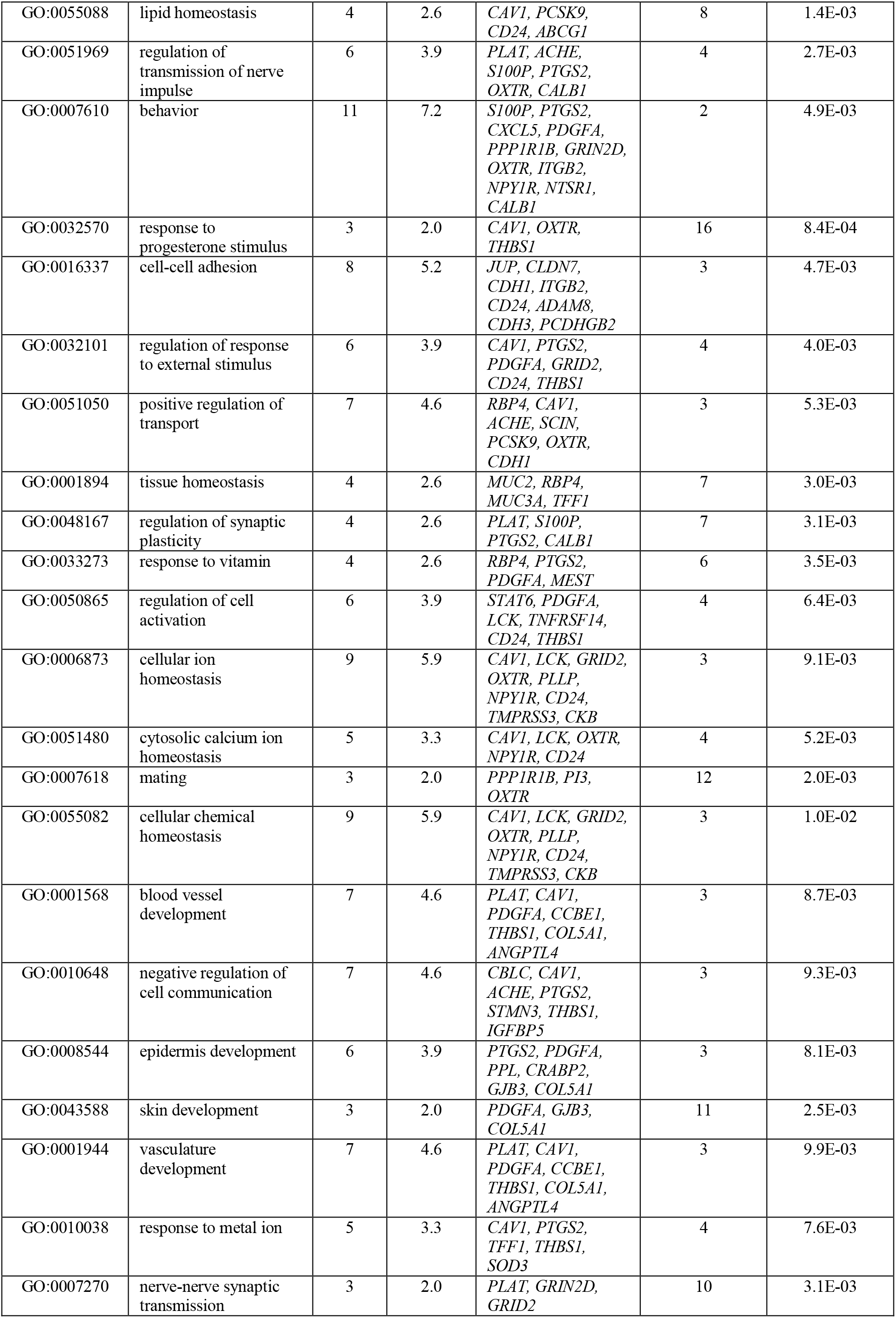
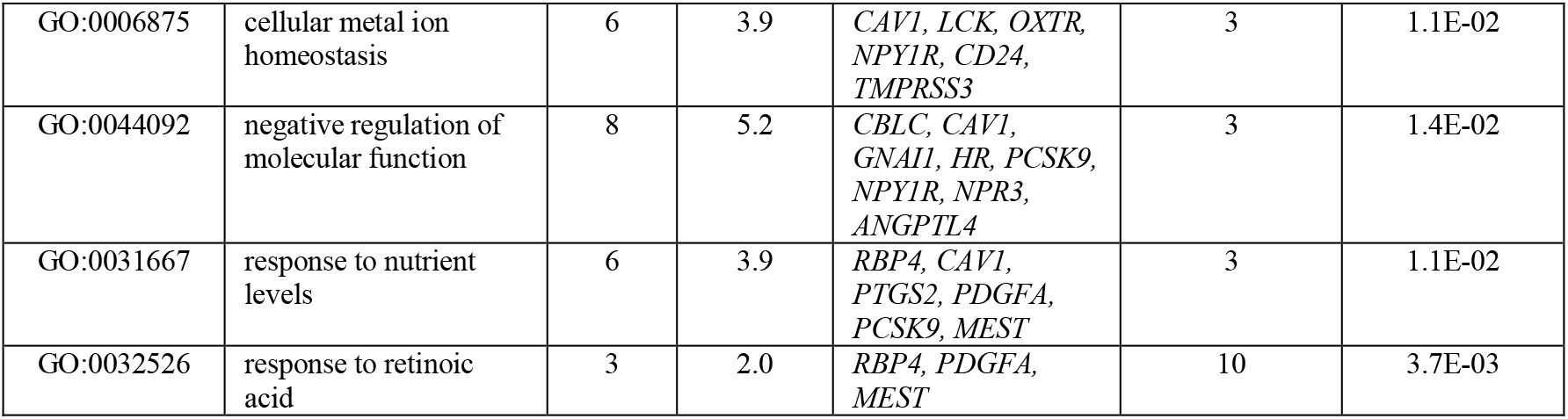
Enrichment for GO terms in the category ‘biological process’ for genes upregulated in PC3-GHSROS cells (compared to empty-vector control). *P* ≤ 0.01, Fisher’s exact test.

**Supplementary Table S5.**
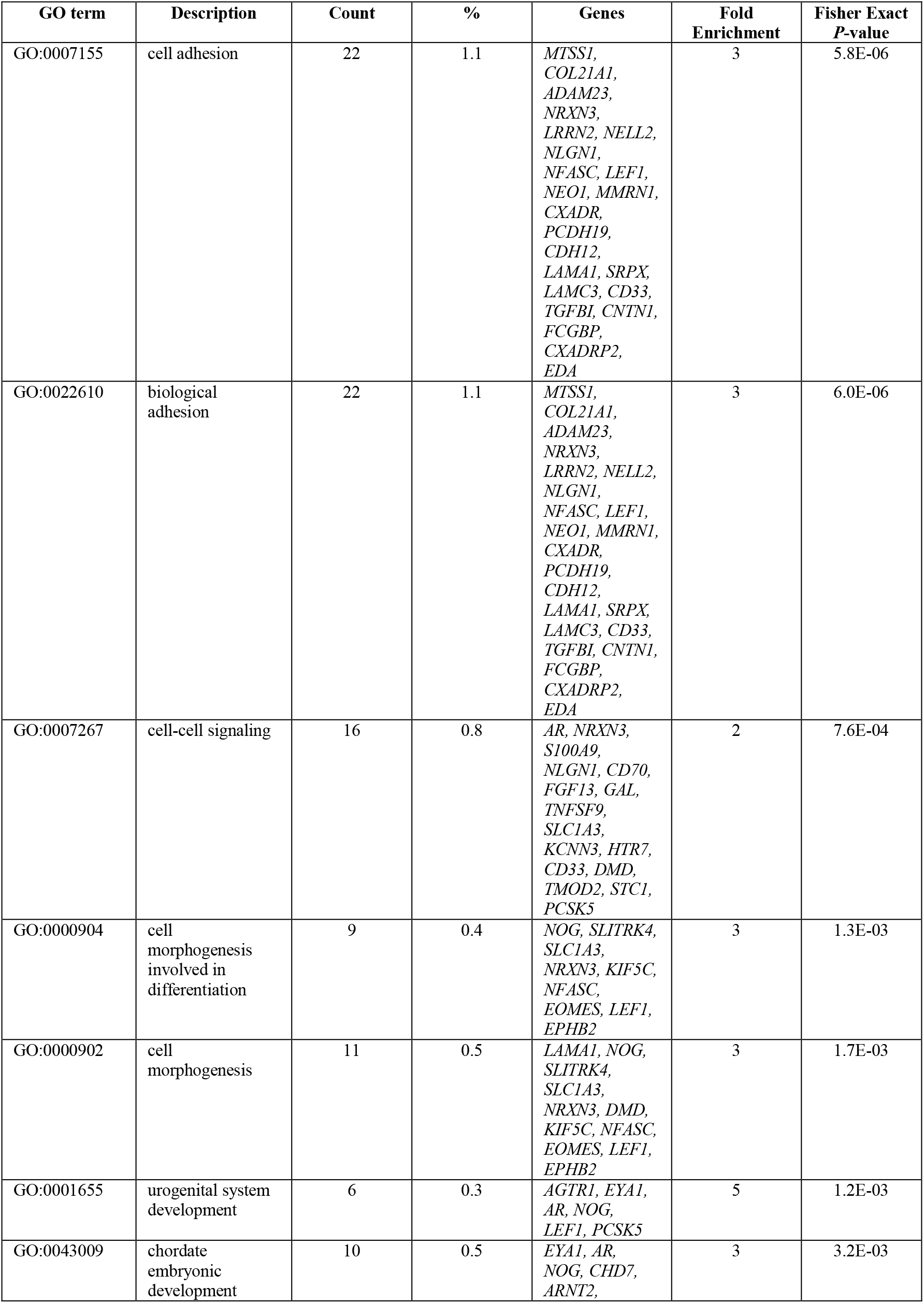
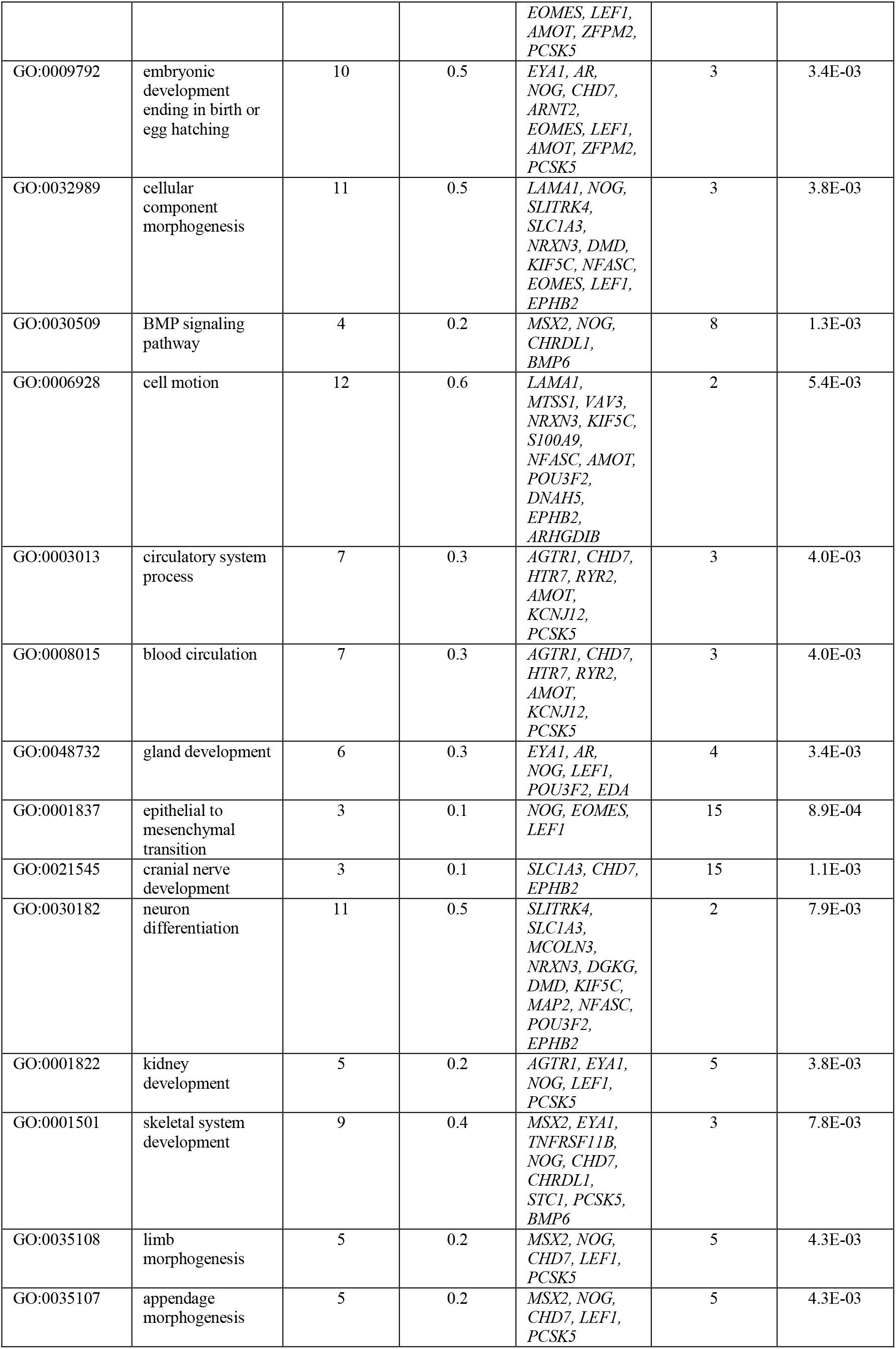
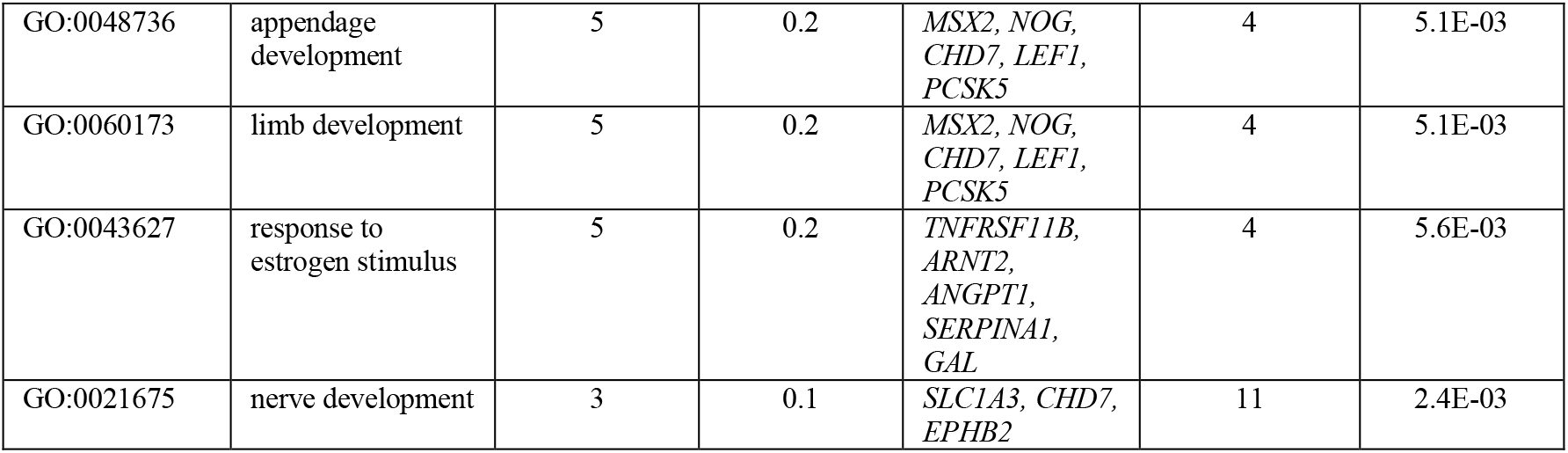
Enrichment for GO terms in the category ‘biological process’ for genes downregulated in PC3-GHSROS cells (compared to empty-vector control). *P* ≤ 0.01, Fisher’s exact test.

**Supplementary Table S6.**
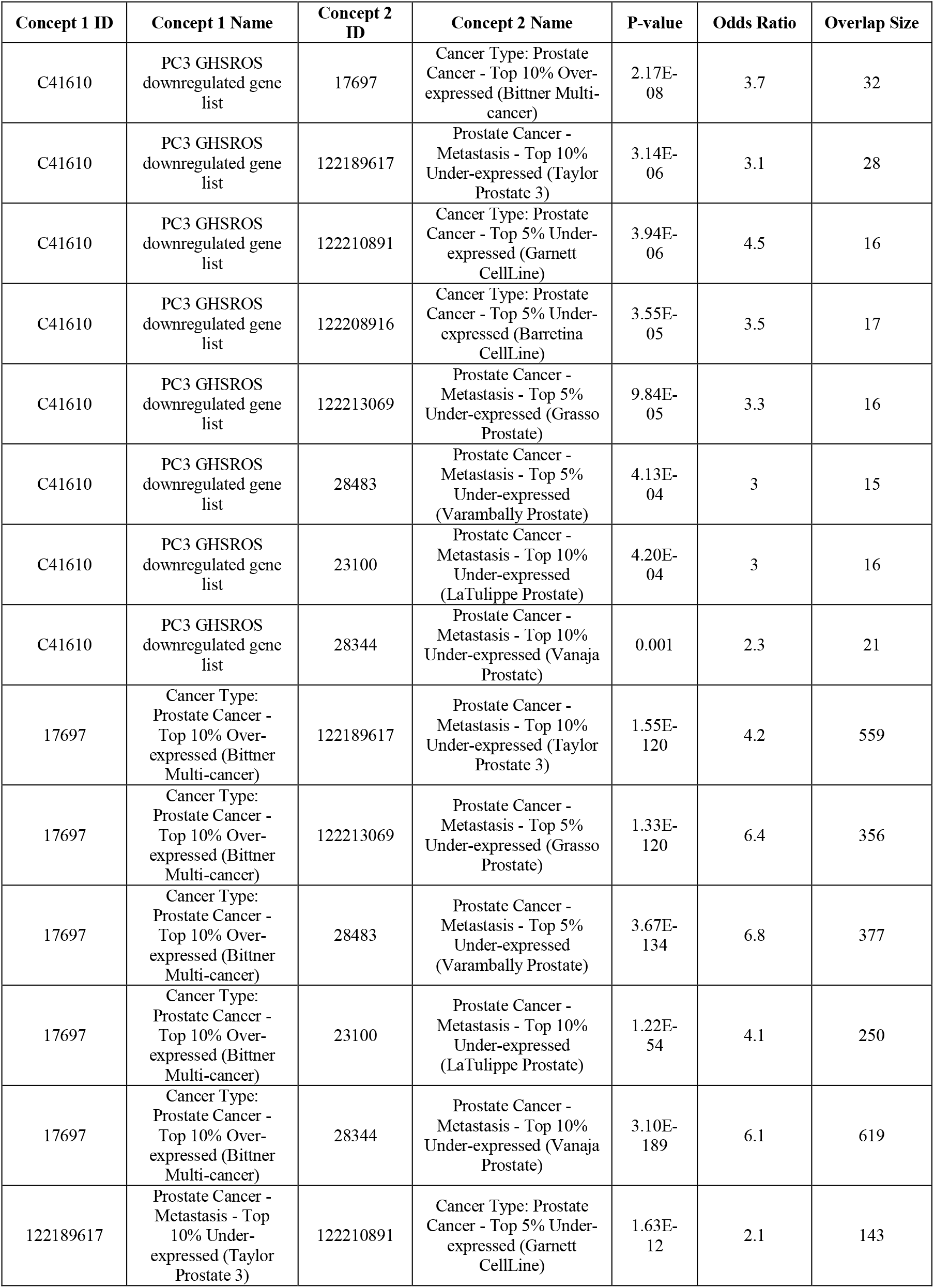
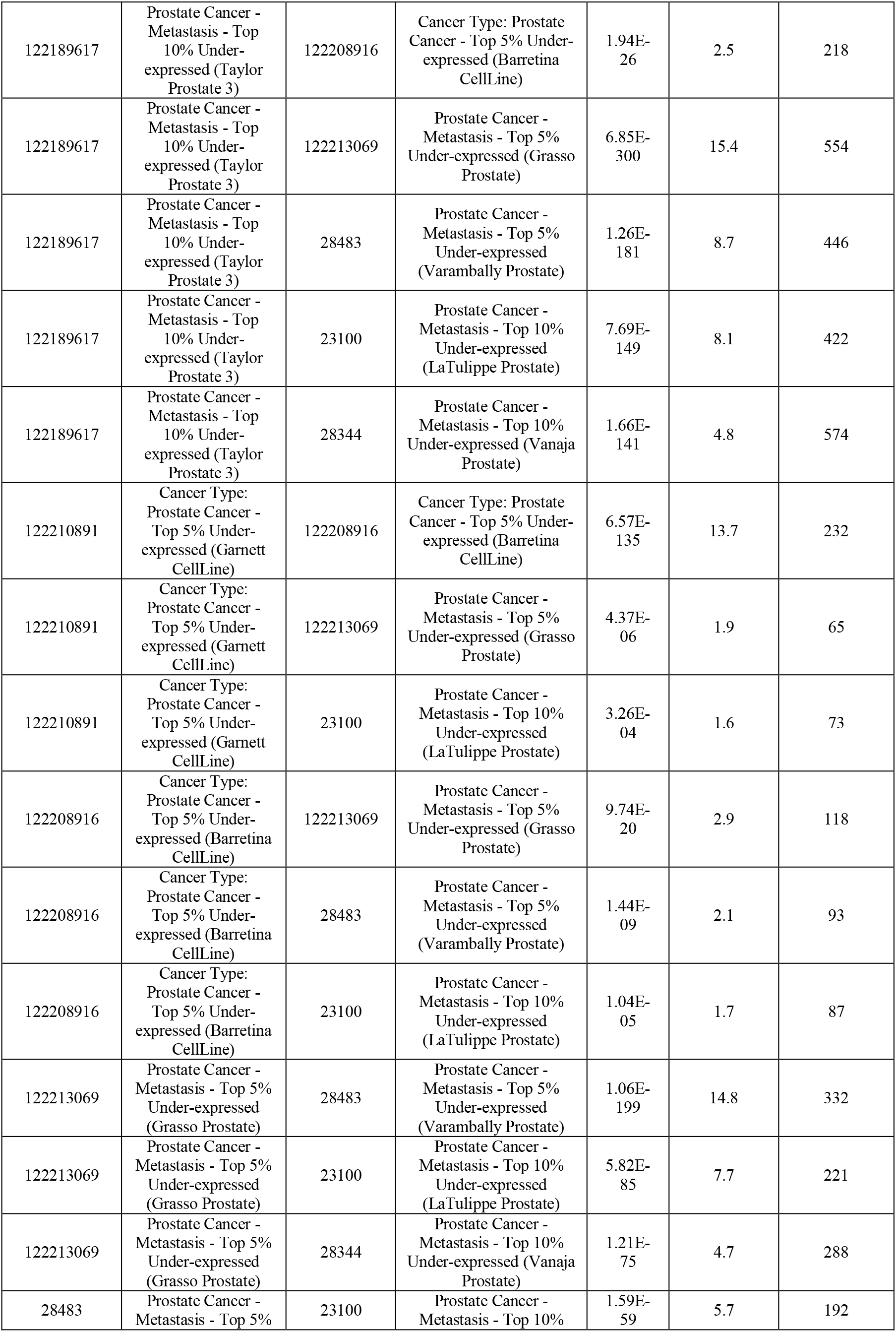
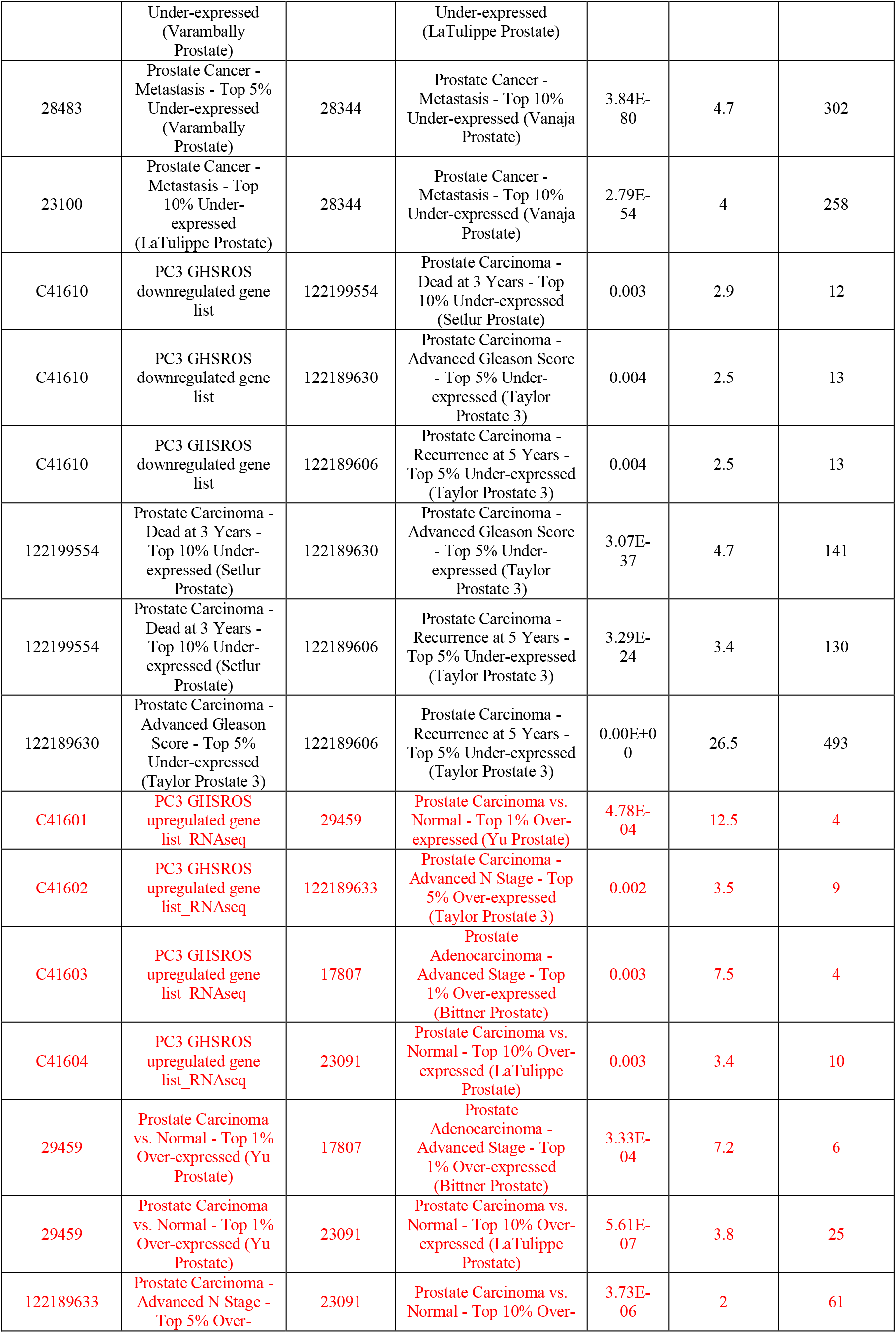
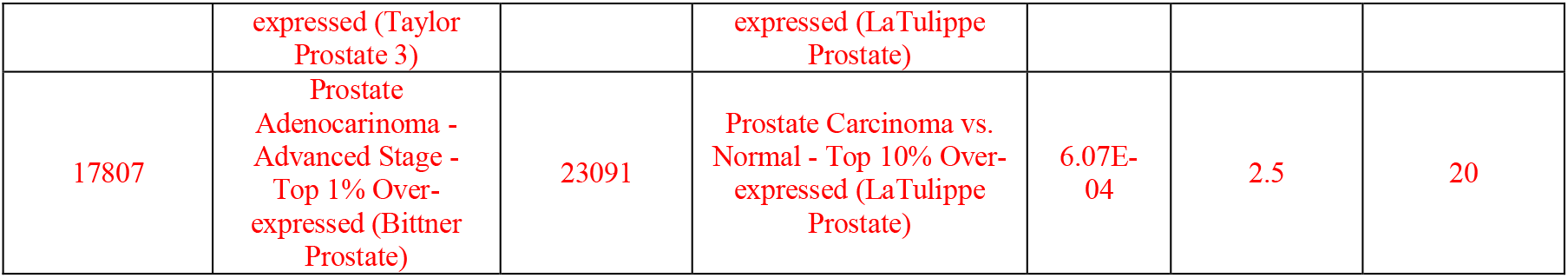
Oncomine concepts analysis of positively and negatively correlated PC3-*GHSROS* gene signature. Red: positively correlated gene signature; Black: negatively correlated gene signature. *P* ≤ 0.01, Fisher’s exact test.

**Supplementary Table S7.**
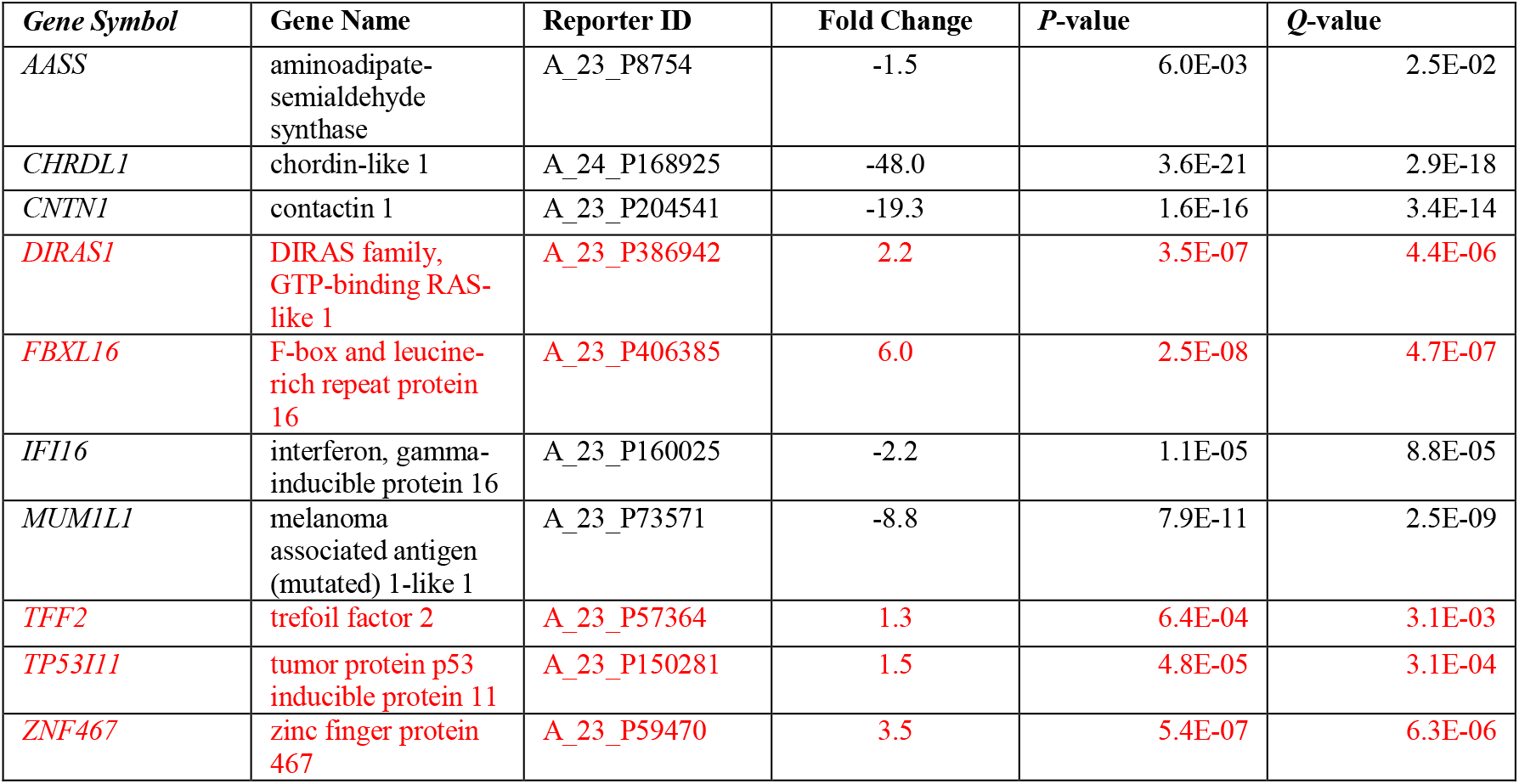
Differentially expressed genes in PC3-GHSROS and LNCaP-GHSROS cells compared to the Grasso Oncomine dataset. The Grasso dataset includes 59 localized and 35 metastatic prostate tumors. Red: higher expression in metastatic tumors; Black: lower expression in metastatic tumors. Fold-changes are log_2_ transformed; *Q*-value denotes the false discovery rate (FDR; Benjamini-Hochberg)-adjusted *P*-value.

**Supplementary Table S8.**
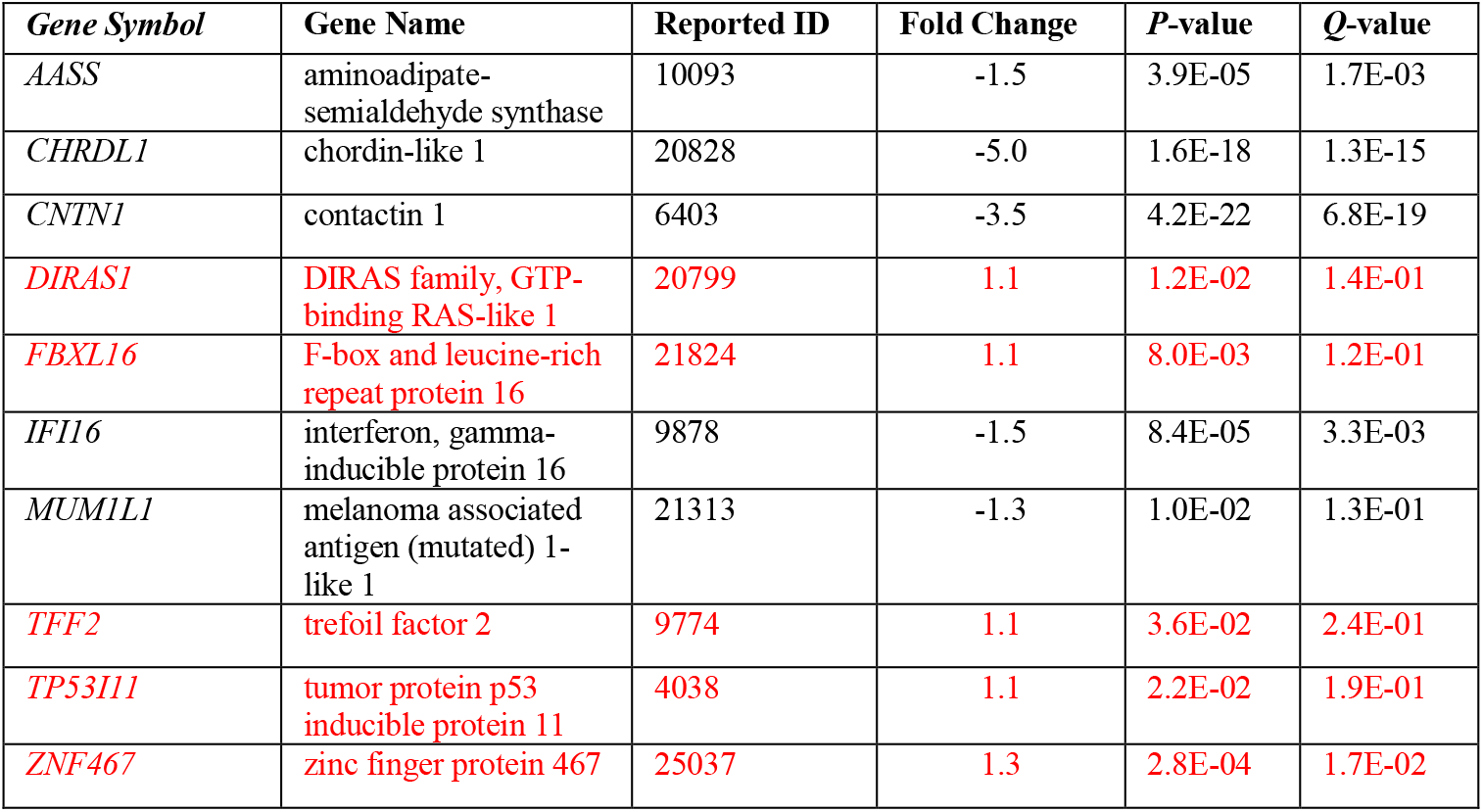
Differentially expressed genes in PC3-GHSROS and LNCaP-GHSROS cells compared to the Taylor Oncomine dataset. The Taylor dataset includes 123 localized and 35 metastatic prostate tumors. Red: higher expression in metastatic tumors; Black: lower expression in metastatic tumors. Fold-changes are log_2_ transformed; *Q*-value denotes the false discovery rate (FDR; Benjamini-Hochberg)-adjusted *P*-value.

**Supplementary Table S9.**
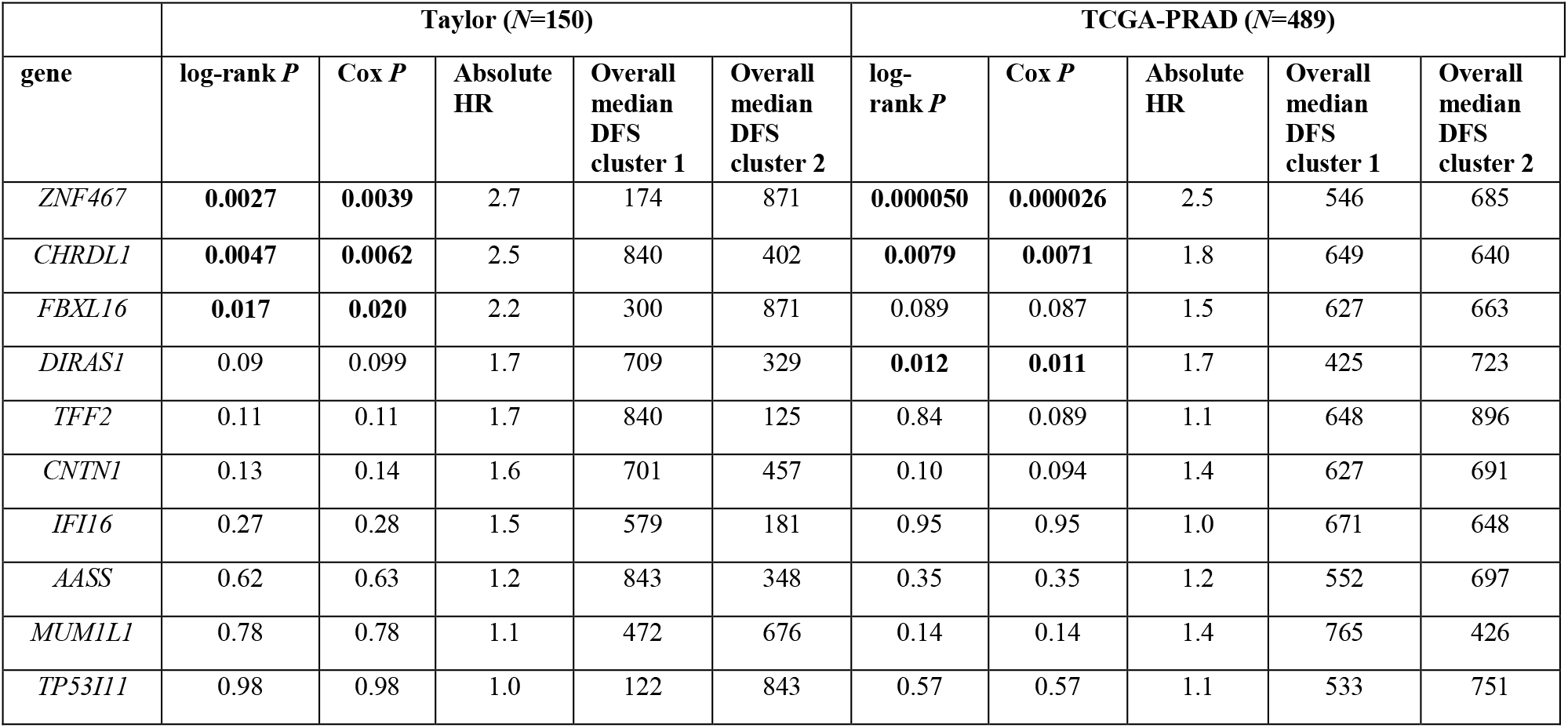
Disease-free survival (DFS) analysis of differentially expressed genes (in PC3-GHSROS cells, LNCaP-GHSROS cells and clinical metastatic tumors) in human datasets. Patients, in the Taylor (*N*=150; *n*=123 localized and *n*=27 metastatic tumors) and TCGA-PRAD (*N*=489; localized tumors) datasets, were stratified into two groups by *k*-means clustering of gene expression (*k*=2). The log-rank test, was used to assign statistical significance, with *P* ≤ 0.05 considered significant (shown in bold). The Cox *P*-value and absolute hazard ratio (HR) between *k*-means cluster 1 and 2 for each gene are indicated. Overall median disease-free survival (DFS) in days are indicated for each cluster.

**Supplementary Table S10.**
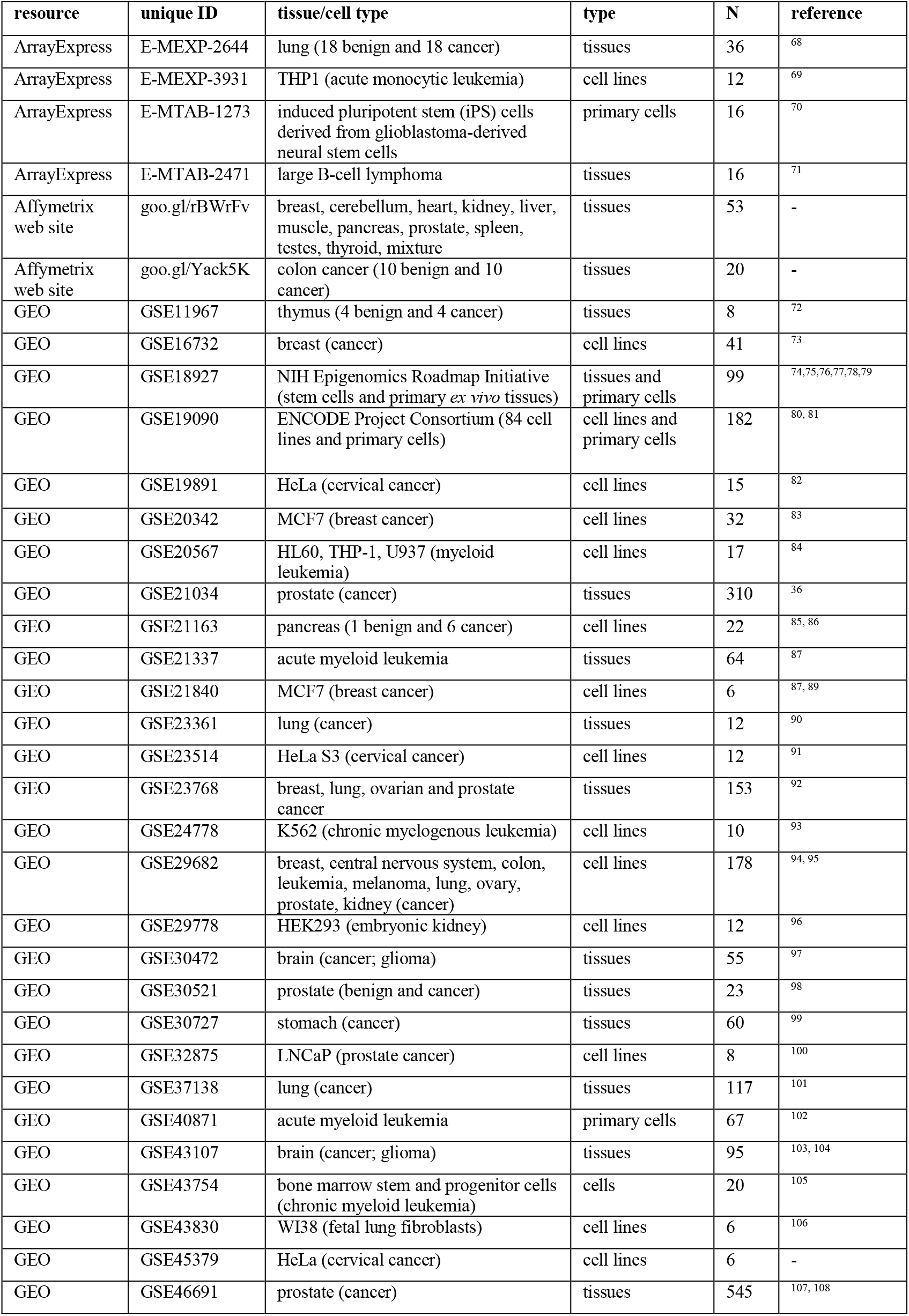
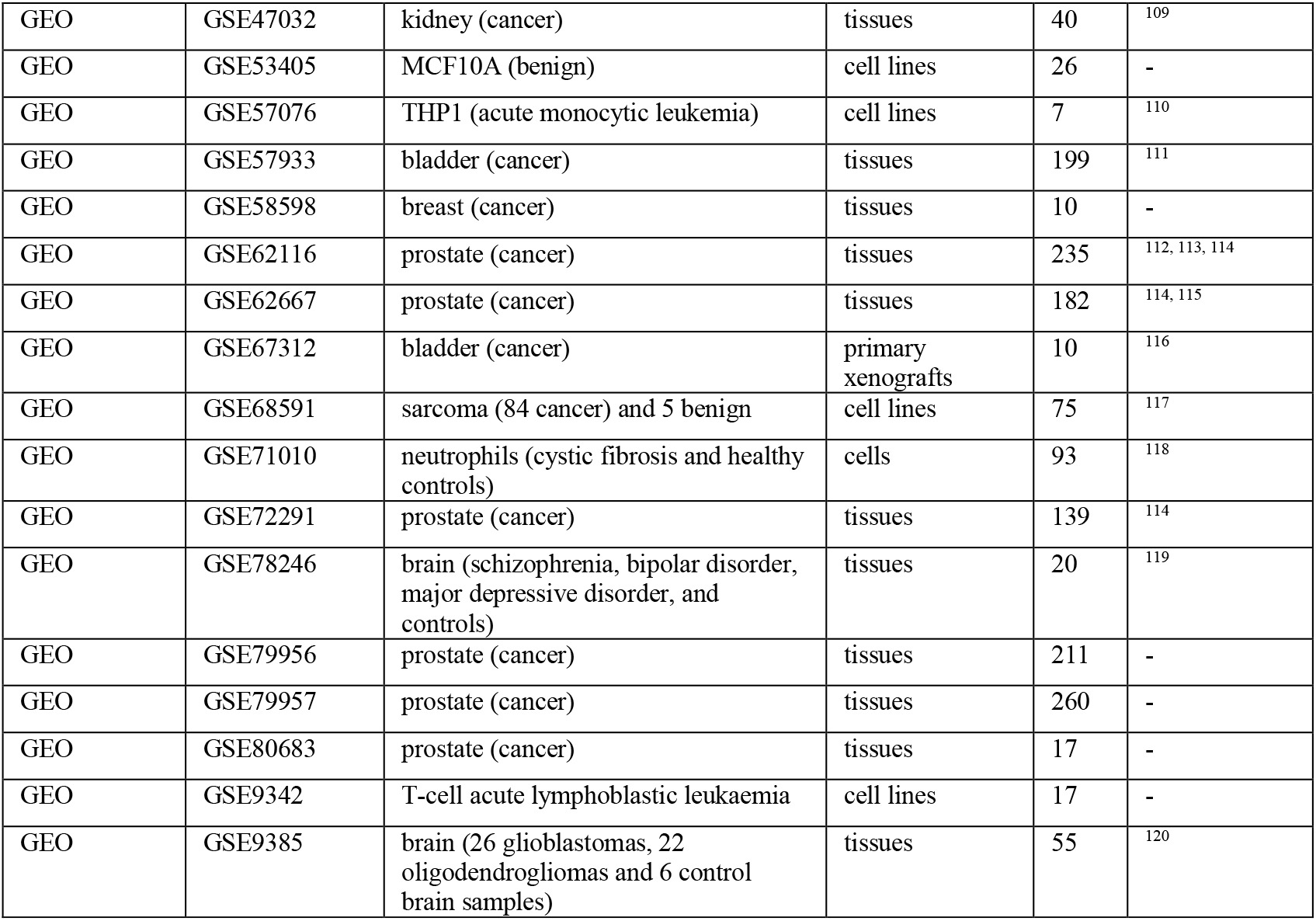
Overview of human Affymetrix exon array datasets interrogated.

| resource | unique ID | tissue/cell type | type | N | reference |
| --- | --- | --- | --- | --- | --- |
| ArrayExpress | E-MEXP-2644 | lung (18 benign and 18 cancer) | tissues | 36 | <sup>68</sup> |
| ArrayExpress | E-MEXP-3931 | THP1 (acute monocytic leukemia) | cell lines | 12 | <sup>69</sup> |
| ArrayExpress | E-MTAB-1273 | induced pluripotent stem (iPS) cells derived from glioblastoma-derived neural stem cells | primary cells | 16 | <sup>70</sup> |
| ArrayExpress | E-MTAB-2471 | large B-cell lymphoma | tissues | 16 | <sup>71</sup> |
| Affymetrix web site | goo.gl/rBWrFv | breast, cerebellum, heart, kidney, liver, muscle, pancreas, prostate, spleen, testes, thyroid, mixture | tissues | 53 | - |
| Affymetrix web site | goo.gl/Yack5K | colon cancer (10 benign and 10 cancer) | tissues | 20 | - |
| GEO | GSE11967 | thymus (4 benign and 4 cancer) | tissues | 8 | <sup>72</sup> |
| GEO | GSE16732 | breast (cancer) | cell lines | 41 | <sup>73</sup> |
| GEO | GSE18927 | NIH Epigenomics Roadmap Initiative (stem cells and primary *ex vivo* tissues) | tissues and primary cells | 99 | <sup>74,75,76,77,78,79</sup> |
| GEO | GSE19090 | ENCODE Project Consortium (84 cell lines and primary cells) | cell lines and primary cells | 182 | <sup>80,81</sup> |
| GEO | GSE19891 | HeLa (cervical cancer) | cell lines | 15 | <sup>82</sup> |
| GEO | GSE20342 | MCF7 (breast cancer) | cell lines | 32 | <sup>83</sup> |
| GEO | GSE20567 | HL60, THP-1, U937 (myeloid leukemia) | cell lines | 17 | <sup>84</sup> |
| GEO | GSE21034 | prostate (cancer) | tissues | 310 | <sup>36</sup> |
| GEO | GSE21163 | pancreas (1 benign and 6 cancer) | cell lines | 22 | <sup>85,86</sup> |
| GEO | GSE21337 | acute myeloid leukemia | tissues | 64 | <sup>87</sup> |
| GEO | GSE21840 | MCF7 (breast cancer) | cell lines | 6 | <sup>87,89</sup> |
| GEO | GSE23361 | lung (cancer) | tissues | 12 | <sup>90</sup> |
| GEO | GSE23514 | HeLa S3 (cervical cancer) | cell lines | 12 | <sup>91</sup> |
| GEO | GSE23768 | breast, lung, ovarian and prostate cancer | tissues | 153 | <sup>92</sup> |
| GEO | GSE24778 | K562 (chronic myelogenous leukemia) | cell lines | 10 | <sup>93</sup> |
| GEO | GSE29682 | breast, central nervous system, colon, leukemia, melanoma, lung, ovary, prostate, kidney (cancer) | cell lines | 178 | <sup>94,95</sup> |
| GEO | GSE29778 | HEK293 (embryonic kidney) | cell lines | 12 | <sup>96</sup> |
| GEO | GSE30472 | brain (cancer; glioma) | tissues | 55 | <sup>97</sup> |
| GEO | GSE30521 | prostate (benign and cancer) | tissues | 23 | <sup>98</sup> |
| GEO | GSE30727 | stomach (cancer) | tissues | 60 | <sup>99</sup> |
| GEO | GSE32875 | LNCaP (prostate cancer) | cell lines | 8 | <sup>100</sup> |
| GEO | GSE37138 | lung (cancer) | tissues | 117 | <sup>101</sup> |
| GEO | GSE40871 | acute myeloid leukemia | primary cells | 67 | <sup>102</sup> |
| GEO | GSE43107 | brain (cancer; glioma) | tissues | 95 | <sup>103,104</sup> |
| GEO | GSE43754 | bone marrow stem and progenitor cells (chronic myeloid leukemia) | cells | 20 | <sup>105</sup> |
| GEO | GSE43830 | WI38 (fetal lung fibroblasts) | cell lines | 6 | <sup>106</sup> |
| GEO | GSE45379 | HeLa (cervical cancer) | cell lines | 6 | - |
| GEO | GSE46691 | prostate (cancer) | tissues | 545 | <sup>107,108</sup> |
| GEO | GSE47032 | kidney (cancer) | tissues | 40 | <sup>109</sup> |
| GEO | GSE53405 | MCF10A (benign) | cell lines | 26 | - |
| GEO | GSE57076 | THP1 (acute monocytic leukemia) | cell lines | 7 | <sup>110</sup> |
| GEO | GSE57933 | bladder (cancer) | tissues | 199 | <sup>111</sup> |
| GEO | GSE58598 | breast (cancer) | tissues | 10 | - |
| GEO | GSE62116 | prostate (cancer) | tissues | 235 | <sup>112, 113, 114</sup> |
| GEO | GSE62667 | prostate (cancer) | tissues | 182 | <sup>114, 115</sup> |
| GEO | GSE67312 | bladder (cancer) | primary xenografts | 10 | <sup>116</sup> |
| GEO | GSE68591 | sarcoma (84 cancer) and 5 benign | cell lines | 75 | <sup>117</sup> |
| GEO | GSE71010 | neutrophils (cystic fibrosis and healthy controls) | cells | 93 | <sup>118</sup> |
| GEO | GSE72291 | prostate (cancer) | tissues | 139 | <sup>114</sup> |
| GEO | GSE78246 | brain (schizophrenia, bipolar disorder, major depressive disorder, and controls) | tissues | 20 | <sup>119</sup> |
| GEO | GSE79956 | prostate (cancer) | tissues | 211 | - |
| GEO | GSE79957 | prostate (cancer) | tissues | 260 | - |
| GEO | GSE80683 | prostate (cancer) | tissues | 17 | - |
| GEO | GSE9342 | T-cell acute lymphoblastic leukaemia | cell lines | 17 | - |
| GEO | GSE9385 | brain (26 glioblastomas, 22 oligodendrogliomas and 6 control brain samples) | tissues | 55 | <sup>120</sup> |

**Supplementary Table S11.**
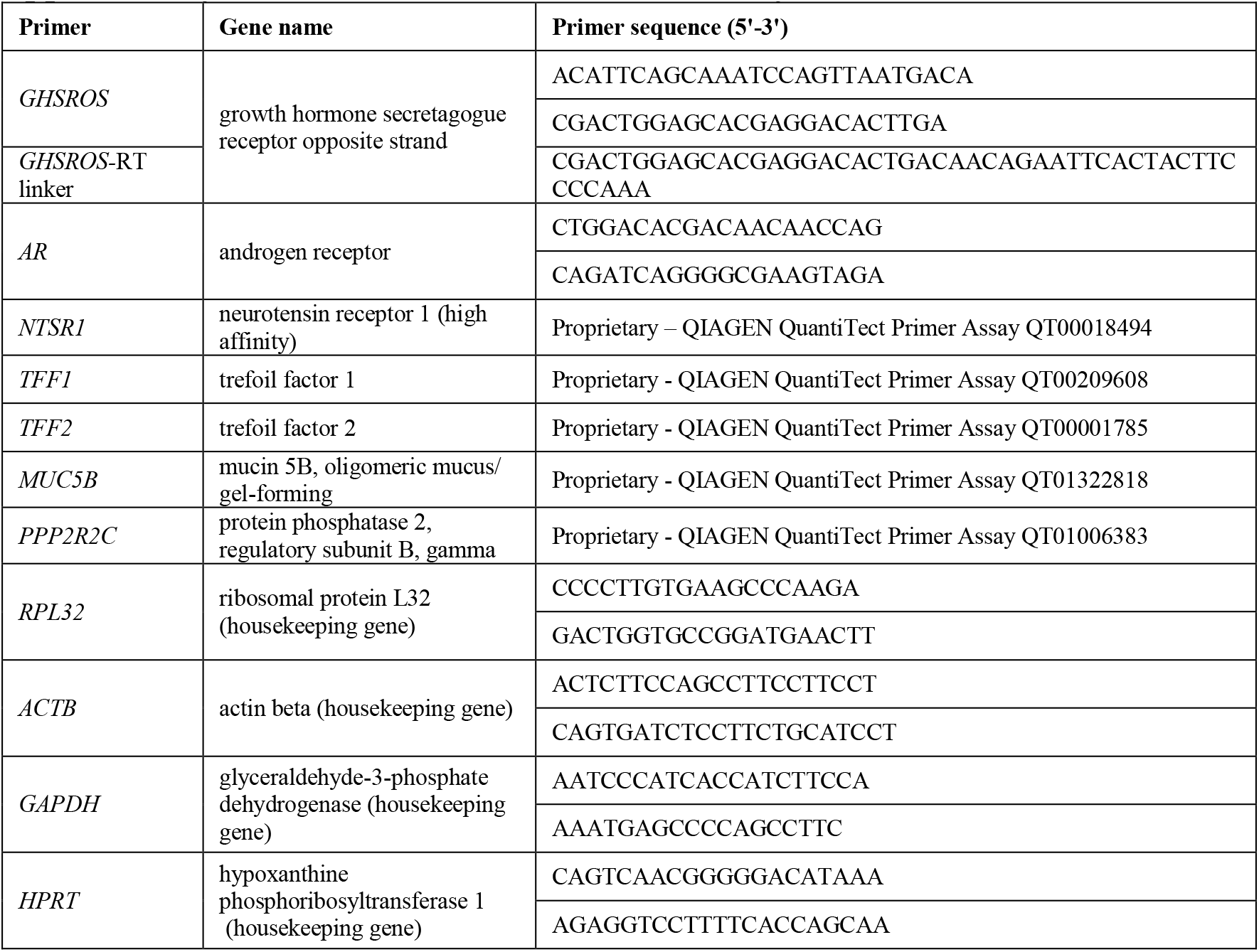
Primers used in this study.

**Supplementary Dataset 1. Differentially expressed genes in LNCaP-GHSROS cells. Compared to empty vector control**. Red: higher expression in LNCaP-GHSROS cells; Black: lower expression in LNCaP-GHSROS cells. Fold-changes are log_2_ transformed; *Q*-value denotes the false discovery rate (FDR; Benjamini-Hochberg)-adjusted *P*-value (cutoff ≤ 0.05).

(provided in a separate file).

**Supplementary Dataset 2. Enrichment for GO terms in the category ‘biological process’ for genes upregulated in LNCaP-GHSROS cells (compared to empty-vector control).** *P* ≤ 0.01, Fisher’s exact test.

(provided in a separate file).

**Supplementary Dataset 3. Enrichment for GO terms in the category ‘biological process’ for genes downregulated in LNCaP-GHSROS cells (compared to empty-vector control).** *P* ≤ 0.01, Fisher’s exact test.

(provided in a separate file).

## SUPPLEMENTARY METHODS

### Identification of *GHSROS* transcription in exon array datasets

To assess *GHSROS* expression, we interrogated Affymetrix GeneChip Exon 1.0 ST arrays, strand-specific oligonucleotide microarrays with probes for known and predicted exons (hereafter termed exon arrays). Exon arrays are comparable to RNA-seq in experiments aimed at assessing exon expression (*i.e.* gene isoforms) and suitable for experiments where the exon of interest is known^121, 122^. In the Exon 1.0 ST array, known (genes and ESTs) and putative exons are combined to form ‘transcript clusters’, with each exon defined as a probe set (typically, a set of 2-4 probes). By combining all probe sets, the expression of a transcript cluster (known or putative gene) can be measured (see https://goo.gl/4RSTG3). To identify probe set(s) corresponding to *GHSROS*, we downloaded the Exon 1.0 ST probe annotation file from NCBI (NCBI Gene Expression Omnibus (GEO) accession no. GPL5188). Full-length *GHSROS* (1.1 kb) was aligned to the human genome (NCBI36/hg18; March 2006 assembly) to generate genomic coordinates compatible with the probe file (chr3:173,646,439-173,647,538). Next, the probe annotation file (GPL5188) was interrogated to reveal probe sets spanning *GHSROS* by entering the following command in a UNIX terminal window:

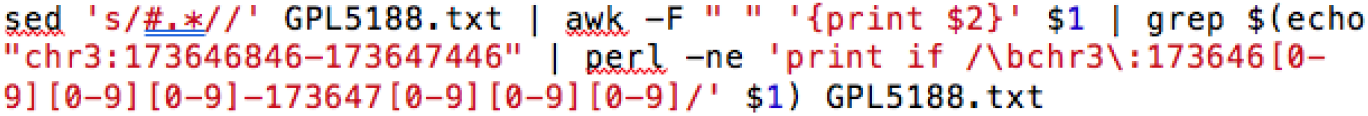

This revealed a probe set, 2652604, consisting of 4 probes complementary to *GHSROS*.

Cell and tissue exon array data were downloaded from NCBI GEO^123^, EBI ArrayExpress^124^ and the Affymetrix web site (see Supplementary Table S10). GEO datasets were bulk-downloaded using v3.6.2.117442 of the Aspera Connect Linux software (Aspera, Emeryville, CA, USA). In total, 3,924 samples were downloaded, corresponding to ∼46% of all exon array data deposited in the NCBI GEO database. Arrays (individual CEL files) were normalized (output on a log_2_ scale, centered at 0) using the *SCAN* function in the R package ‘SCAN.UPC’^125, 126^. SCAN normalizes each array (sample) individually by removing background noise (probe- and array-specific) data from within the array. Next, arrays were interrogated using the *UPC* function in ‘SCAN.UPC’. UPC outputs standardized expression values (UPC value), ranging from 0 to 1, which indicate whether a gene is actively transcribed in a sample of interest: higher values indicate that a gene is ‘active’^126^. UPC scores are platform-independent and allow cross-experimental and cross-platform integration.

### Evaluation of *GHSR/GHSROS* transcription in deep RNA-seq dataset

It has been estimated that reliable detection of low abundance transcripts in humans warrants very deep sequencing (> 200 million reads per sample^127^) – far beyond most current datasets. To illustrate, we considered the expression of *GHSR*/*GHSROS* in a comparable clinical dataset. Publicly available RNA-seq data (NCBI GEO accession no. GSE31528) from eight subjects with metastatic castration-resistant prostate cancer (bone marrow metastases)^128^ were interrogated. Briefly, total RNA-seq was performed on random-primed paired end read libraries, to ensure consistent transcript coverage^128, 129^, generating an average of 160M reads per sample. Paired-end FASTQ files were aligned to the human genome (UCSC build hg19) using the spliced-read mapper TopHat (v2.0.9)^130^ and reference gene annotations to guide the alignment. BigWig sequencing tracks for the UCSC genome browser^131, 132^ were obtained from TopHat-generated BAM files (indexed by samtools v1.2^133^) using a local instance of the *bamCoverage* command in deepTools v2.5.4^134^. BigWig files were visualized in the UCSC genome browser (hg19). A region with less than ∼10 supporting reads can be considered to have low coverage, rendering active transcription difficult to interpret^127, 135^.

### Cell culture, prostate cancer patient derived xenograft (PDX) models, and treatments

The cancer cell lines PC3 (ATCC CRL-1435), DU145 (ATCC HTB-81), LNCaP (ATCC CRL-1740), ES-2 (ATCC CRL-1978), A549 (ATCC CCL-185), and 22Rv1 (ATCC CRL-2505) were obtained from the American Type Culture Collection (ATCC, Rockville, MD, USA). The C4-2B^66^ and DUCaP^67^ prostate cancer cell lines, six LuCaP prostate derived xenograft (PDX) lines^17^, and the BM18 PDX cell line^18^ were available in our laboratory. All prostate cancer and ovarian cancer cell lines were maintained in Roswell Park Memorial Institute (RPMI) 1640 medium (RPMI-1640; Invitrogen, Carlsbad, CA) with 10 % Fetal Calf Serum (FCS, Thermo Fisher Scientific Australia, Scoresby, VIC, Australia), supplemented with 100 U/mL penicillin G and 100 ng/mL streptomycin (Invitrogen). The A549 lung cancer cell line was maintained in Dulbecco’s Modified Eagle Medium: Nutrient Mixture F-12 (DMEM/F12) medium (Invitrogen) with 10% FCS (Thermo Fisher Scientific Australia) supplemented with 100 U/mL penicillin G and 100 ng/mL streptomycin (Invitrogen). The non-tumorigenic RWPE-1 (ATCC CRL-11609) and the transformed, tumorigenic RWPE-2 (ATCC CRL-11610) prostate epithelium-derived cell lines were cultured in keratinocyte serum-free medium (Invitrogen) supplemented with 50 µg/mL bovine pituitary extract and 5 ng/mL epidermal growth factor (Invitrogen). All cell lines were passaged at 2- to 3-day intervals on reaching 70 % confluency using TrypLE Select (Invitrogen). Cell morphology and viability were monitored by microscopic observation and regular Mycoplasma testing was performed (Universal Mycoplasma Detection Kit, ATCC). For drug treatments, cells were treated with 10 µM enzalutamide (ENZ; Selleck Chemicals, Houston, TX, USA) or 10-100 nM docetaxel (DTX; Sigma Aldrich, St. Louis, MO, USA) for 96 (functional assays) or 48 hours (qRT-PCR) and compared to dimethyl sulfoxide (DMSO) (Sigma Aldrich, St. Louis, MO, USA) vehicle control.

### Production of *GHSROS* overexpressing cancer cell lines

Full-length *GHSROS* transcript was cloned into the *pTargeT* mammalian expression vector (Promega, Madison, WI). PC3, DU145, and A549 cell lines were transfected with *GHSROS-pTargeT* DNA, or vector alone (empty vector), (using Lipofectamine LTX, Invitrogen) according to the manufacturer’s instructions. Cells were incubated for 24 hours in LTX and selected with geneticin (100-1500 µg/mL G418, Invitrogen). As LNCaP prostate cancer cells were difficult to transfect using lipid-mediated transfection, we employed lentiviral transduction. Briefly, *pReceiver-Lv105* vectors, expressing full length *GHSROS,* or empty control vectors, were obtained from GeneCopoeia (Rockville, MD). For stable overexpression, LNCaP cells were seeded at 50-60% confluency and transduced with *GHSROS,* or empty vector control lentiviral constructs in the presence of 8 µg/ml polybrene (Sigma Aldrich). Following a 48-hour incubation period, transduced cells were selected with 1 µg/mL puromycin (Invitrogen). *GHSROS* expression was confirmed approximately 3 weeks after selection by qRT-PCR, every 2-3 weeks, and before every functional experiment (see Supplementary Fig. S5).

### RNA extraction, reverse transcription and quantitative reverse transcription Polymerase Chain Reaction (qRT-PCR)

Total RNA was extracted from cell pellets using an RNeasy Plus Mini Kit (QIAGEN, Hilden, Germany) with a genomic DNA (gDNA) Eliminator spin column. To remove contaminating genomic DNA, 1 µg RNA was DNase treated prior to cDNA synthesis with Superscript III (Invitrogen). qRT-PCR was performed using the AB7500 FAST sequence detection thermal cycler (Applied Biosystems, Foster City, CA), or the ViiA Real-Time PCR system (Applied Biosystems) with SYBR Green PCR Master Mix (QIAGEN) using primers listed in Supplementary Table S11. A negative control (water instead of template) was used in each real-time plate for each primer set. All real-time experiments were performed in triplicate. Baseline and threshold values (C_t_) were obtained using ABI 7500 Prism and the relative expression of mRNA was calculated using the comparative 2^-ΔΔCt^ method^136^. Expression was normalized to the housekeeping gene ribosomal protein L32 (*RPL32*). Statistical analyses were performed using GraphPad Prism v.6.01 software (GraphPad Software, Inc., San Diego, CA). Student’s *t*-test or Mann-Whitney-Wilcoxon tests were used to assess the statistical significance of all the direct comparisons.

### *GHSROS* qRT-PCR interrogation of human tissue specimens

To survey the expression of *GHSROS* in cancer, we initially interrogated a TissueScan Cancer Survey Tissue qPCR panel (CSRT102; OriGene, Rockville, MD, USA); cDNA arrayed on multi-well PCR plates. Some of the samples are normal non-malignant tissue samples, making it possible to compare expression in tumor versus normal tissue. For each cancer type, data were expressed as mean fold change using the comparative 2^-ΔΔCt^ method against a non-malignant control tissue. Normalized to β-actin (*ACTB*).

To further investigate the expression of *GHSROS* in prostate cancer TissueScan Prostate Cancer Tissue qPCR panels (HPRT101, HPRT102, and HPRT103) were obtained from OriGene. The cDNA panels contained of a total of 24 normal prostate-derived samples, 31 abnormal prostate samples (defined as lesions), and 88 prostate tumor samples. These panels were examined by qRT-PCR, using the method described above, except that the housekeeping gene ribosomal protein L32 (*RPL32*) was employed.

An independent cohort was obtained from the Andalusian Biobank (Servicio Andaluz de Salud, Spain). It consisted of tissue from 28 patients with clinical high-grade prostate cancer (10 localized and 18 metastatic tumors) and 8 normal prostate tissue samples. RT-PCR was performed using Brilliant III SYBR Green Master Mix and a Stratagene Mx3000p instrument (both from Agilent, La Jolla, CA, USA), as previously described^137^. Briefly, samples were on the same plate were analysed with a standard curve to estimate mRNA copy number (tenfold dilutions of synthetic cDNA template for each transcript). No-RNA controls were carried out for all primer pairs. To control for variations in the amount of RNA used, and the efficiency of the reverse-transcription reaction, the expression level (copy number) of each transcript was adjusted by a normalization factor (NF) obtained from the expression of three housekeeping genes (*ACTB*, *HPRT*, and *GAPDH*) using the geNorm algorithm^138^. Primers used are listed in Supplementary Table S11.

### Locked Nucleic Acid-Antisense Oligonucleotides (LNA-ASO)

Two distinct LNA ASOs, RNV104L and RNV124, complementary to different regions of *GHSROS* (see Supplementary Fig. S6), were designed in-house and synthesized commercially (Exiqon, Vedbæk, Denmark). The ASOs contained two consecutive LNA nucleotides at the 5’-end and three consecutive LNA nucleotides at the 3’-end – in line with gapmer design principles. RNV104L contained LNA nucleotides at positions 2, 3, 16, 17, and 18; RNV124 at positions 3, 4, 18, 19, and 20. The LNA ASO sequences were as follows: scrambled control sequence: 5’-GC<u>TT</u>CGACTCGTAATCA<u>CCT</u>A-3’; RNV124 (underlined bases denote LNA nucleotides): 5’-AT<u>AA</u>ACCTGCTAGTGTC<u>CTC</u>C-3’; RNV104L: 5’-G<u>TT</u>AACTTTCTTCTT<u>CCT</u>TG-3’. Lyophilized oligonucleotides were resuspended in ultrapure H2O (Invitrogen) and stored as a 100 µM stock solution at −20°C. Briefly, LNA-ASOs were diluted to 20 µM in OptiMEM I Reduced Serum Medium (Invitrogen) and cultured cells were transfected according to the manufacturer’s instructions. Cultured cells were incubated at 37°C in 5% CO2 for 4 hours, before 500 µl growth medium, containing 30% FCS, was added to the serum-free medium. The cells were transfected for 24-72 hours and *GHSROS* levels assessed by qRT-PCR.

### RNA secondary structure prediction

The ViennaRNA web server was employed^139^ to predict the secondary structure of *GHSROS* and its minimum free energy^140, 141^.

### Cell proliferation assays

Proliferation assays were performed using an xCELLigence real-time cell analyzer (RTCA) DP instrument (ACEA Biosciences, San Diego, CA). This system employs sensor impedance technology to quantify the status of the cell using a unit-less parameter termed the cell index (CI). The CI represents the status of the cell based on the measured relative changes in electrical impedance that occur in the presence and absence of cells in the wells (generated by the software, according to the formula CI = (Z_i_ – Z_0_)/15 Ω, where Z_i_ is the impedance at an individual point of time during the experiment and Z_0_ the impedance at the start of the experiment). Impedance is measured at three different frequencies (10, 25 or 50 kHz). Briefly, 5 × 10^3^ cells were trypsinized and seeded into a 96 well plate (E-plate) and grown for 48 hours in 150 µl growth media. Cell index was measured every 15 minutes and all experiments were performed in triplicate, with at least three independent repeats. Because cells did not attach well to the gold microelectrodes of the xCELLigence instrument, LNCaP proliferation was quantified by measuring the cleavage of WST-1 (Roche, Basel, Switzerland). Briefly, 5 × 10^4^ cells/ well were seeded in 96-well plates (BD Biosciences, Franklin Lakes, NJ) and propagated for 72 hours in complete medium. To determine cell number, absorbance was measured using the FLUOstar Omega spectrophotometer (BMG, Ortenberg, Germany) at 440 nm using a reference wavelength of 600 nm. All proliferation experiments were performed independently three times, with 8 replicates each.

### Cell Viability Assay

LNCaP and PC3 vector or *GHSROS* over-expressing cells (5000 cells/well) were seeded in 96-well plates (BD Biosciences) and propagated overnight in complete medium. LNCaP cells were treated with standard doses of test compounds in both charcoal stripped FCS (CSS) or 2% FCS. PC3 cells were treated with increasing doses of docetaxel in 2% FCS. After a 96-hour period cell viability was measured using a WST-1 cell proliferation assay (Roche, Nonnenwald, Penzberg, Germany) according to the manufacturer’s instructions. All viability experiments were performed independently three times, with 4 replicates each.

### Cell Migration assays

Migration assays were performed using an xCELLigence RTCA DP instrument (ACEA Biosciences). Briefly, 5 × 10^4^ cells/well were seeded on the top chamber in 150 µl serum-free media. The lower chamber contained 160 µl media with 10% FCS as a chemo-attractant. Cell index was measured every 15 minutes for 24 hours to indicate the rate of cell migration to the lower chamber. All experiments were performed in triplicate with at least 3 independent repeats. Because cells did not attach well to the gold microelectrodes of the xCELLigence instrument, LNCaPs migration was assessed using a transwell assay. Briefly, 6 × 10^5^ cells were suspended in serum-free medium and added to the upper chamber of inserts coated with a polycarbonate membrane (8 µm pore size; BD Biosciences). Cells in 12-well plates were allowed to migrate for 24 h in response to a chemoattractant (10% FBS) in the lower chamber. After 24 h, cells remaining in the upper chamber were removed. Cells that had migrated to the lower surface of the membrane were fixed with methanol (100%) and stained with 1% crystal violet. Acetic acid (10%, v/v) was used to extract the crystal violet and absorbance was measured at 595 nm. Each experiment consisted of three replicates and was repeated independently three times.

### Mouse subcutaneous *in vivo* xenograft models

All mouse studies were carried out with approval from the University of Queensland and the Queensland University of Technology Animal Ethics Committees. PC3-GHSROS, PC3-vector, DU145-GHSROS, DU145-vector, LNCaP-GHSROS, and LNCaP-vector cell lines were injected subcutaneously into the flank of 4-5-week-old male NSG mice^142^ (obtained from Animal Resource Centre, Murdoch, WA, Australia). Cells were injected in a 1:1 ratio with growth factor-reduced Matrigel (Thermo Fisher) (*n*=8-10 per cell line) and tumors measured twice weekly with digital calipers (ProSciTech, Kirwan, QLD, Australia). Neither randomization nor blinding for animal use was performed because we commercially obtained these mice with the same genetic background. Animals were euthanized once tumor volume reached 1,000 mm^3^, or at other ethical endpoints. At the experimental endpoint, the primary tumor was resected, divided in half, snap frozen and stored at −80°C.

### Histology and immunohistochemistry

For histological analysis, cryosections (6-10 μm thick) were prepared using a Leica CM1850 cryotome (Wetzlar, Germany). Sections were collected onto warm, charged Menzel Superfrost slides (Thermo Fisher), fixed in ice-cold 100% acetone, air dried and stored at −80 °C. For immunohistochemistry, tissues were fixed in paraformaldehyde and dehydrated through a graded series of ethanol and xylene, before being embedded in paraffin. Sections (5µm) were mounted on to glass Menzel Superfrost slides ThermoFisher Scientific). Immunohistochemistry was performed using antibodies for the proliferation marker Ki67 (rabbit anti-human Ki67, Abcam, Cambridge, UK) and for the infiltration of murine blood vessels using rabbit anti-murine CD31 antibody (Abcam). Tissue sections were incubated with HRP-polymer conjugates (SuperPicture, Thermo Fisher Scientific), and incubated with the chromagen diaminobenzidine (DAB) (Dako, Glostrup, Denmark), as per manufacturer’s specifications. Slides were counterstained with Mayer’s hematoxylin, dehydrated, and mounted with coverslips using D.P.X neutral mounting medium (Sigma-Aldrich). All sections were counterstained with Mayer’s hematoxylin (Sigma Aldrich) and mounted with coverslips using D.P.X with Colourfast (Fronine, ThermoFisher Scientific).

### RNA sequencing of PC3-GHSROS cells

RNA was extracted from *in vitro* cultured PC3-GHSROS cells and controls, as outlined in the manuscript body. RNA purity was analysed using an Agilent 2100 Bioanalyzer, and RNA with an RNA Integrity Number (RIN) above 7 used for RNA-seq. Strand-specific RNA-sequencing (RNA-seq) was performed by Macrogen, South Korea. A TruSeq stranded mRNA library (Illumina) was constructed and RNA sequencing performed (50 million reads) on a HiSeq 2000 instrument (Illumina) with 100bp paired end reads. Pre-processing of raw FASTQ reads, including elimination of contamination adapters, was performed with scythe v0.994 (https://github.com/vsbuffalo/scythe). Paired-end human FASTQ files were aligned to the human genome, UCSC build hg19 using the spliced-read mapper TopHat (v2.0.9)^130^ and reference gene annotations to guide the alignment.

Raw gene counts were computed from TopHat-generated BAM files using featureCounts v1.4.5-p1^143^, counting coding sequence (CDS) features of the UCSC hg19 gene annotation file (gtf). FeatureCounts output files were analysed using the R programming language (v.3.2.2). Briefly, raw counts were normalized by Trimmed Mean of M-values (TMM) correction^144, 145^. Library size-normalized read counts (per million; CPM) were subjected to the voom function (variance modelling at the observation-level) in limma v3.22.1 (Linear Models for Microarray Data)^146, 147^, with trend=TRUE for the eBayes function and correction for multiple testing (Benjamini-Hochberg false discovery rate of cut-off, *Q*-value, set at 0.05). Genes with at least a 1.5 log_2_ fold-change difference in expression between PC3-GHSROS and PC3-vector (empty vector) cells were defined as differentially expressed. Although validation is not required, as RNA-seq gives very accurate measurements of relative expression across a broad dynamic range^148^, selected differentially regulated genes were validated using quantitative reverse-transcription PCR (qRT-PCR) (see manuscript body and table S11).

Detailed gene annotations were obtained by querying Ensembl with the R/Bioconductor package ‘biomaRt’^149^. Gene Ontology (GO) term analyses were performed using DAVID (Database for Annotation, Visualization and Integrated Discovery)^150^. Briefly, to test for enrichment we interrogated DAVID’s GO FAT database with genes differentially expressed in PC3-*GHSROS* cells. The DAVID functional annotation tool categorizes GO terms and calculates an ‘enrichment score’ or EASE score (a modified Fisher’s exact test-derived *P*-value). Categories with smaller *P*-values (*P* ≤ 0.01) and larger fold-enrichments (≥ 2.0) were considered interesting and most likely to convey biological meaning^150^.

To perform Oncomine meta-analysis, genes differentially expressed in PC3-GHSROS were separated into ‘over-expressed’ and ‘under-expressed’ gene sets. The Oncomine database^27^ was interrogated by importing these genes, and enriched concepts were generated and ordered by *P*-values (calculated using Fisher’s exact test). Only datasets with an odds ratio ≥ 3.0 and a *P-*value ≤ 0.01 were retained. The datasets were exported as nodes and edges for network visualization in Cytoscape^151^ (v3.4.0). The network layout and node position were generated using the Force-Directed Layout algorithm^152^, with odds ratio as the leading parameter for the edge weight. Using our custom concept generated lists, we next sought to assess the differential expression of our gene lists in two prostate cancer microarray datasets: Grasso^35^ (59 localized and 35 metastatic prostate tumors) and Taylor^36^ (123 localized and 27 metastatic prostate tumors). Differentially expressed genes were ranked and results exported as fold change (log_2_ transformed, median centered). Data was filtered for significance with *P*-value set at ≤ 0.05 and Benjamini-Hochberg false discovery rate (FDR) *Q*-value^153^ at ≤ 0.25; a threshold deemed suitable to find biologically relevant transcriptional signatures^154, 155^.

### RNA sequencing of LNCaP-GHSROS cells

RNA was extracted from LNCaP-GHSROS xenograft tumors and controls (empty vector control lentiviral constructs), as outlined in the manuscript body. RNA purity was analysed using an Agilent 2100 Bioanalyzer, and RNA with an RNA Integrity Number (RIN) above 7 used for RNA-seq. Strand-specific RNA-seq was performed by the South Australian Health and Medical Research Institute (SAHMRI, Adelaide, SA, Australia). A TruSeq stranded mRNA library (Illumina) was constructed and RNA sequencing performed (35 million reads) on a Nextseq 500 instrument (Illumina) with 75bp single end reads. Pre-processing of raw FASTQ reads, including elimination of contamination adapters, was performed with scythe v0.994 (https://github.com/vsbuffalo/scythe). Human (xenograft tumor; the graft) and mouse (the host) RNA-seq reads were separated using Xenome^156^ on the trimmed FASTQ files, leaving ∼20M human reads. Reads were aligned to the human genome and processed as described for PC3-GHSROS cells above. Genes differentially expressed in LNCaP-GHSROS cells (cutoff set at log_2_ 1.5-fold-change and *Q* ≤ 0.05) were imported into the GSEA (Gene Set Enrichment Analysis) program^157^.

### LP50 prostate cancer cell line *AR* knockdown microarray

Publicly available Affymetrix HG-U133 Plus 2.0 microarray data (NCBI GEO accession no. GSE22483) from a substrain of the LNCaP cell line: androgen-independent late passage LNCaP cells (LP50) was interrogated. This cell line was subjected to androgen receptor (*AR*) knockdown by shRNA^64^. The array (*n*=2, of AR shRNA and scrambled control) was normalized to housekeeping genes using the Affymetrix Gene Chip Operating System v1.4^64^. Prior to differential expression analysis, the probe set was pre-filtered, using the R statistical programming language, as follows: probes with mean expression values in the lowest 20^th^ percentile of the array was removed. Differential expression was determined by the R package ‘limma’^139^ and probes with a Benjamini-Hochberg adjusted *P*-value (*Q*; BH-FDR) ≤ 0.05 considered significant. Gene annotations were obtained using the R/Bioconductor packages ‘Biobase’^158^ and ‘GEOquery’^159^.

### Survival analysis

Two datasets were interrogated: Taylor^36^(123 localized and 27 metastatic prostate tumors) and TCGA-PRAD from The Cancer Genomics Atlas (TCGA) consortium, which contains tumors from patients with moderate- (∼39% Gleason 6 and 3 + 4) and high- (∼61% Gleason 4+3 and Gleason 8-10) risk localized prostate carcinoma^37^. Briefly, in the case of TCGA-PRAD, the UCSC Xena Browser^160^ was used to obtain normalized gene expression values, represented as log_2_(normalized counts+1), from the ‘TCGA TARGET GTeX’ dataset consisting of ∼12,000 tissue samples from 31 cancers^161^. To obtain up-to-date overall survival (OS) and disease-free survival (DFS) information, we manually queried cBioPortal for Cancer Genomics^162, 163^ (last accessed 05.08.16).

We performed non-hierarchical *k*-means clustering^164^ to partition patients into groups with similar gene expression patterns^165^. The following 10 genes obtained by Oncomine meta-analysis (see above) were assessed: *AASS*, *CHRDL1*, *CNTN1*, *DIRAS1*, *FBXL16*, *IFI16*, *MUM1L1*, *TP53I11*, *TFF2*, and *ZNF467.* Clustering was performed using the *kmeans* function in the R package ‘stats’ with two clusters/groups (*k=2*) and the best cluster pair after 500 runs (*nstart=500*) was retained^166^. Kaplan-Meier survival analysis^167^ was performed with the R package ‘survival’^168^, fitting survival curves (*survfit*) and computing log-rank *P*-values using the *survdiff* function, with *rho=0* (equivalent to the method employed by UCSC Xena; see https://goo.gl/4knf62). Survival curves were plotted when survival was significantly different between two groups (log-rank *P* ≤ 0.05). We used the *coxph* function in the R package ‘survival’ to test the prognostic significance of genes (that is: we implemented the Cox proportional hazard model to analyze the association of gene expression with patient survival)^169, 170^, with *P* ≤ 0.05 (Wald test) considered significant. Because there is a single categorical covariate (*k*-means cluster; group), the *P*-values from the log-rank and the Cox regression tests are comparable. We considered groups (clusters) that had fewer than 10 samples with a recorded event unreliable.

A scaled heat map (unsupervised hierarchical clustering by Euclidean distance) was generated in R using heatmap.3 (available at https://goo.gl/Yd9aTY) and a custom R script.

### Statistical analyses

Data values were expressed as mean ± s.e.m. of at least two independent experiments and evaluated using Student’s *t*-test for unpaired samples, or otherwise specified. Mean differences were considered significant when *P* ≤ 0.05. *Q*-values denote multiple testing correction (Benjamini-Hochberg) adjusted *P*-values^153^. Normalized high-throughput gene expression data were analyzed using LIMMA, employing a modified version of the Student’s *t*-test (moderated *t*-test) where the standard errors are reduced toward a common value using an empirical Bayesian model robust for datasets with few biological replicates^147^. Statistical analyses were performed using GraphPad Prism v.6.01 software (GraphPad Software, Inc., San Diego, CA), or the R statistical programming language.

### Code

Selected R code is available in a repository at https://github.com/sciseim/GHSROS_MS. Additional R and bash scripts can be obtained by contacting the corresponding authors.

## REFERENCES

1. Mattick J.S., Rinn J.L. Discovery and annotation of long noncoding RNAs. Nat Struct Mol Biol 2015; 22: 5–7.

2. Huarte M. The emerging role of lncRNAs in cancer. Nat Med 2015; 21: 1253–1261.

3. Whiteside E.J. et al. Identification of a long non-coding RNA gene, growth hormone secretagogue receptor opposite strand, which stimulates cell migration in non-small cell lung cancer cell lines. Int J Oncol 2013; 43: 566–574.

4. Saxonov S., Berg P., Brutlag D.L. A genome-wide analysis of CpG dinucleotides in the human genome distinguishes two distinct classes of promoters. Proc Natl Acad Sci U S A 2006; 103: 1412–1417.

5. Derrien T. et al. The GENCODE v7 catalog of human long noncoding RNAs: analysis of their gene structure, evolution, and expression. Genome Res 2012; 22: 1775–1789.

6. Fitzmaurice C. et al. The Global Burden of Cancer 2013. JAMA Oncol 2015; 1: 505–527.

7. Tosoian J.J., Antonarakis E.S. Molecular heterogeneity of localized prostate cancer: more different than alike. Transl Cancer Res 2017; 6: S47–S50.

8. Shoag J., Barbieri C.E. Clinical variability and molecular heterogeneity in prostate cancer. Asian J Androl 2016; 18: 543–548.

9. Dagogo-Jack I., Shaw A.T. Tumour heterogeneity and resistance to cancer therapies. Nat Rev Clin Oncol 2018; 15: 81–94.

10. Mouraviev V. et al. Clinical prospects of long noncoding RNAs as novel biomarkers and therapeutic targets in prostate cancer. Prostate Cancer Prostatic Dis 2016; 19: 14–20.

11. Ruiz-Orera J., Messeguer X., Subirana J.A., Alba M.M. Long non-coding RNAs as a source of new peptides. eLife 2014; 3: e03523.

12. Kutter C. et al. Rapid turnover of long noncoding RNAs and the evolution of gene expression. PLoS Genet 2012; 8: e1002841.

13. Cabili M.N. et al. Integrative annotation of human large intergenic noncoding RNAs reveals global properties and specific subclasses. Genes Dev 2011; 25: 1915–1927.

14. Necsulea A. et al. The evolution of lncRNA repertoires and expression patterns in tetrapods. Nature 2014; 505: 635–640.

15. Wang K.C. et al. A long noncoding RNA maintains active chromatin to coordinate homeotic gene expression. Nature 2011; 472: 120–124.

16. Geisler S., Coller J. RNA in unexpected places: long non-coding RNA functions in diverse cellular contexts. Nat Rev Mol Cell Biol 2013; 14: 699–712.

17. Nguyen H.M. et al. LuCaP Prostate Cancer Patient-Derived Xenografts Reflect the Molecular Heterogeneity of Advanced Disease and Serve as Models for Evaluating Cancer Therapeutics. Prostate 2017; 77: 654–671.

18. McCulloch D.R., Opeskin K., Thompson E.W., Williams E.D. BM18: A novel androgen-dependent human prostate cancer xenograft model derived from a bone metastasis. Prostate 2005; 65: 35–43.

19. Komura K. et al. Resistance to docetaxel in prostate cancer is associated with androgen receptor activation and loss of KDM5D expression. Proc Natl Acad Sci U S A 2016; 113: 6259–6264.

20. Drake C.G., Sharma P., Gerritsen W. Metastatic castration-resistant prostate cancer: new therapies, novel combination strategies and implications for immunotherapy. Oncogene 2014; 33: 5053–5064.

21. Sonpavde G., Wang C.G., Galsky M.D., Oh W.K, Armstrong A.J. Cytotoxic chemotherapy in the contemporary management of metastatic castration-resistant prostate cancer (mCRPC). BJU Int 2015; 116: 17–29.

22. Chandrasekar T., Yang J.C., Gao A.C., Evans C.P. Mechanisms of resistance in castration-resistant prostate cancer (CRPC). Transl Androl Urol 2015; 4: 365–380.

23. Lee H.Y. et al. Clinical predictor of survival following docetaxel-based chemotherapy. Oncol Lett 2014; 8: 1788–1792.

24. Tran C. et al. Development of a second-generation antiandrogen for treatment of advanced prostate cancer. Science 2009; 324: 787–790.

25. Lin D. et al. Next generation patient-derived prostate cancer xenograft models. Asian J Androl 2014; 16: 407–412.

26. Hanahan D., Weinberg R.A. Hallmarks of cancer: the next generation. Cell 2011; 144: 646–674.

27. Rhodes D.R. et al. Oncomine 3.0: genes, pathways, and networks in a collection of 18,000 cancer gene expression profiles. Neoplasia 2007; 9: 166–180.

28. Seim I., Jeffery P.L., Thomas P.B., Nelson C.C., Chopin L.K. Whole-genome sequence of the metastatic PC3 and LNCaP human prostate cancer cell lines. G3 (Bethesda) 2017; 7: 1731-1741.

29. Szklarczyk D. et al. The STRING database in 2017: quality-controlled protein-protein association networks, made broadly accessible. Nucleic Acids Res 2017; 45: D362–D368.

30. Putzke A.P. et al. Metastatic progression of prostate cancer and e-cadherin regulation by ZEB1 and SRC family kinases. Am J Pathol 2011; 179: 400–410.

31. Legrier M.E. et al. Mucinous differentiation features associated with hormonal escape in a human prostate cancer xenograft. Br J Cancer 2004; 90: 720–727.

32. Vestergaard E.M., Borre M., Poulsen S.S., Nexo E., Torring N. Plasma levels of trefoil factors are increased in patients with advanced prostate cancer. Clin Cancer Res 2006; 12: 807–812.

33. Kani K. et al. Anterior gradient 2 (AGR2): blood-based biomarker elevated in metastatic prostate cancer associated with the neuroendocrine phenotype. Prostate 2013; 73: 306–315.

34. Zweitzig D.R., Smirnov D.A., Connelly M.C., Terstappen L.W., O’Hara S.M., Moran E. Physiological stress induces the metastasis marker AGR2 in breast cancer cells. Mol Cell Biochem 2007; 306: 255–260.

35. Grasso C.S. et al. The mutational landscape of lethal castration-resistant prostate cancer. Nature 2012; 487: 239–243.

36. Taylor B.S et al. Integrative genomic profiling of human prostate cancer. Cancer Cell 2010; 18: 11–22.

37. Cancer Genome Atlas Research Network. The Molecular Taxonomy of Primary Prostate Cancer. Cell 2015; 163: 1011–1025.

38. Cyr-Depauw C. et al. Chordin-Like 1 suppresses bone morphogenetic protein 4-induced breast cancer cell migration and invasion. Mol Cell Biol 2016; 36: 1509–1525.

39. Chandran U.R., Dhir R., Ma C., Michalopoulos G., Becich M., Gilbertson J. Differences in gene expression in prostate cancer, normal appearing prostate tissue adjacent to cancer and prostate tissue from cancer free organ donors. BMC Cancer 2005; 5: 45.

40. Castro M.A. et al. Regulators of genetic risk of breast cancer identified by integrative network analysis. Nat Genet 2016; 48: 12–21.

41. Jhun M.A. et al. Gene expression signature of Gleason score is associated with prostate cancer outcomes in a radical prostatectomy cohort. Oncotarget 2017; 8: 43035–43047.

42. Zhu L., Hu Z., Liu J., Gao J., Lin B. Gene expression profile analysis identifies metastasis and chemoresistance-associated genes in epithelial ovarian carcinoma cells. Med Oncol 2015; 32: 426.

43. Davies G.F. et al. TFPI1 mediates resistance to doxorubicin in breast cancer cells by inducing a hypoxic-like response. PLoS One 2014; 9: e84611.

44. Bluemn E.G. et al. PPP2R2C loss promotes castration-resistance and is associated with increased prostate cancer-specific mortality. Mol Cancer Res 2013; 11: 568–578.

45. Gonzalez-Alonso P., Cristobal I., Manso R., Madoz-Gurpide J., Garcia-Foncillas J., Rojo F.. PP2A inhibition as a novel therapeutic target in castration-resistant prostate cancer. Tumour Biol 2015; 36: 5753–5755.

46. Zhu H., Zhu X., Zheng L., Hu X., Sun L., Zhu X. The role of the androgen receptor in ovarian cancer carcinogenesis and its clinical implications. Oncotarget 2017; 8: 29395–29405.

47. Harlos C., Musto G., Lambert P., Ahmed R., Pitz M.W. Androgen pathway manipulation and survival in patients with lung cancer. Horm Cancer 2015; 6: 120–127.

48. Liu S.J. et al. CRISPRi-based genome-scale identification of functional long noncoding RNA loci in human cells. Science 2017; 355.

49. Chiu H.S. et al. Pan-Cancer Analysis of lncRNA Regulation Supports Their Targeting of Cancer Genes in Each Tumor Context. Cell Rep 2018; 3: 297–312.

50. Cabanski C.R. et al. Pan-cancer transcriptome analysis reveals long noncoding RNAs with conserved function. RNA Biol 2015; 12: 628–642.

51. Puente J., Grande E., Medina A., Maroto P., Lainez N., Arranz J.A. Docetaxel in prostate cancer: a familiar face as the new standard in a hormone-sensitive setting. Ther Adv Med Oncol 2017; 9: 307–318.

52. Belkhiri A., Zhu S., El-Rifai W. DARPP-32: from neurotransmission to cancer. Oncotarget 2016; 7: 17631–17640.

53. Prensner J.R. et al. The long noncoding RNA SChLAP1 promotes aggressive prostate cancer and antagonizes the SWI/SNF complex. Nat Genet 2013; 45: 1392–1398.

54. Ferraldeschi R., Welti J., Luo J., Attard G., de Bono J.S. Targeting the androgen receptor pathway in castration-resistant prostate cancer: progresses and prospects. Oncogene 2015; 34: 1745–1757.

55. Wyatt A.W., Gleave M.E. Targeting the adaptive molecular landscape of castration-resistant prostate cancer. EMBO Mol Med 2015; 7: 878–894.

56. Bluemn E.G. et al. Androgen receptor pathway-independent prostate cancer is sustained through FGF signaling. Cancer Cell 2017; 32: 474–489.

57. Peng D. et al. Integrated molecular analysis reveals complex interactions between genomic and epigenomic alterations in esophageal adenocarcinomas. Sci Rep 2017; 7: 40729.

58. Bi D., Ning H., Liu S., Que X., Ding K. miR-1301 promotes prostate cancer proliferation through directly targeting PPP2R2C. Biomed Pharmacother 2016; 81: 25-30.

59. Yan L., Cai K., Liang J., Liu H., Liu Y., Gui J. Interaction between miR-572 and PPP2R2C, and their effects on the proliferation, migration, and invasion of nasopharyngeal carcinoma (NPC) cells. Biochem Cell Biol 2017; 95: 578–584.

60. Wu A.H. et al. miR-572 prompted cell proliferation of human ovarian cancer cells by suppressing PPP2R2C expression. Biomed Pharmacother 2016; 77: 92–97.

61. Fan Y.L., Chen L., Wang J., Yao Q., Wan J.Q. Over expression of PPP2R2C inhibits human glioma cells growth through the suppression of mTOR pathway. FEBS Lett 2013; 587: 3892–3897.

62. Sengupta R. et al. CXCR4 activation defines a new subgroup of Sonic hedgehog-driven medulloblastoma. Cancer Res 2012; 72: 122–132.

63. Auvergne R.M. et al. Transcriptional differences between normal and glioma-derived glial progenitor cells identify a core set of dysregulated genes. Cell Rep 2013; 3: 2127–2141.

64. Gonit M. et al. Hormone depletion-insensitivity of prostate cancer cells is supported by the AR without binding to classical response elements. Mol Endocrinol 2011; 25: 621–634.

65. Stein C.A., Castanotto D. FDA-Approved Oligonucleotide Therapies in 2017. Mol Ther 2017; 25: 1069–1075.

66. Thalmann G.N. et al. Androgen-independent cancer progression and bone metastasis in the LNCaP model of human prostate cancer. Cancer Res 1994; 54: 2577–2581.

67. Lee Y.G., Korenchuk S., Lehr J., Whitney S., Vessela R., Pienta K.J. Establishment and characterization of a new human prostatic cancer cell line: DuCaP. In Vivo 2001; 15: 157–162.

## SUPPLEMENTARY REFERENCES

68. Langer W. et al. Exon array analysis using re-defined probe sets results in reliable identification of alternatively spliced genes in non-small cell lung cancer. BMC Genomics 2010; 11: 676.

69. Folkard D.L. et al. Suppression of LPS-induced transcription and cytokine secretion by the dietary isothiocyanate sulforaphane. Mol Nutr Food Res 2014; 58: 2286–2296.

70. Stricker S.H. et al. Widespread resetting of DNA methylation in glioblastoma-initiating cells suppresses malignant cellular behavior in a lineage-dependent manner. Genes Dev 2013; 27: 654–669.

71. Riihijarvi S. et al. Prognostic influence of macrophages in patients with diffuse large B-cell lymphoma: a correlative study from a Nordic phase II trial. Haematologica 2015; 100: 238–245.

72. Soreq L. et al. Identifying alternative hyper-splicing signatures in MG-thymoma by exon arrays. PLoS One 2008; 3: e2392.

73. Riaz M. et al. Low-risk susceptibility alleles in 40 human breast cancer cell lines. BMC Cancer 2009; 9: 236.

74. Bernstein B.E. et al. The NIH Roadmap Epigenomics Mapping Consortium. Nat Biotechnol 2010; 28: 1045–1048.

75. Maurano M.T. et al. Systematic localization of common disease-associated variation in regulatory DNA. Science 2012; 337: 1190–1195.

76. Neph S. et al. An expansive human regulatory lexicon encoded in transcription factor footprints. Nature 2012; 489: 83–90.

77. Polak P. et al. Cell-of-origin chromatin organization shapes the mutational landscape of cancer. Nature 2015; 518: 360–364.

78. Polak P. et al. Reduced local mutation density in regulatory DNA of cancer genomes is linked to DNA repair. Nat Biotechnol 2014; 32: 71–75.

79. Schultz M.D. et al. Human body epigenome maps reveal noncanonical DNA methylation variation. Nature 2015; 523: 212–216.

80. Hansen R.S. et al. Sequencing newly replicated DNA reveals widespread plasticity in human replication timing. Proc Natl Acad Sci U S A 2010; 107: 139–144.

81. Thurman R.E. et al. The accessible chromatin landscape of the human genome. Nature 2012; 489: 75–82.

82. Younis I. et al. Rapid-response splicing reporter screens identify differential regulators of constitutive and alternative splicing. Mol Cell Biol 2010; 30: 1718–1728.

83. Dutertre M. et al. Estrogen regulation and physiopathologic significance of alternative promoters in breast cancer. Cancer Res 2010; 70: 3760–3770.

84. Kitamura H. et al. Ubiquitin-specific protease 2-69 in macrophages potentially modulates metainflammation. FASEB J 2013; 27: 4940–4953.

85. Omura N. et al. Cyclooxygenase-deficient pancreatic cancer cells use exogenous sources of prostaglandins. Mol Cancer Res 2010; 8: 821–832.

86. Vincent A. et al. Genome-wide analysis of promoter methylation associated with gene expression profile in pancreatic adenocarcinoma. Clin Cancer Res 2011; 17: 4341–4354.

87. Risueno A. et al. A robust estimation of exon expression to identify alternative spliced genes applied to human tissues and cancer samples. BMC Genomics 2014; 15: 879.

88. Dutertre M. et al. A recently evolved class of alternative 3’-terminal exons involved in cell cycle regulation by topoisomerase inhibitors. Nat Commun 2014; 5: 3395.

89. Dutertre M. et al. Cotranscriptional exon skipping in the genotoxic stress response. Nat Struct Mol Biol 2010; 17: 1358–1366.

90. Calverley D.C. et al. Significant downregulation of platelet gene expression in metastatic lung cancer. Clin Transl Sci 2010; 3: 227–232.

91. Llorian M. et al. Position-dependent alternative splicing activity revealed by global profiling of alternative splicing events regulated by PTB. Nat Struct Mol Biol 2010; 17: 1114–1123.

92. Kan Z. et al. Diverse somatic mutation patterns and pathway alterations in human cancers. Nature 2010; 466: 869–873.

93. Pencovich N., Jaschek R., Tanay A., Groner Y. Dynamic combinatorial interactions of RUNX1 and cooperating partners regulates megakaryocytic differentiation in cell line models. Blood 2011; 117: e1–e14.

94. Kohn K.W., Zeeberg B.M., Reinhold W.C., Pommier Y. Gene expression correlations in human cancer cell lines define molecular interaction networks for epithelial phenotype. PLoS One 2014; 9: e99269.

95. Reinhold W.C. et al. Exon array analyses across the NCI-60 reveal potential regulation of TOP1 by transcription pausing at guanosine quartets in the first intron. Cancer Res 2010; 70: 2191–2203.

96. Mukherjee N. et al. Integrative regulatory mapping indicates that the RNA-binding protein HuR couples pre-mRNA processing and mRNA stability. Mol Cell 2011; 43: 327–339.

97. Gravendeel L.A. et al. Gene expression profiles of gliomas in formalin-fixed paraffin-embedded material. Br J Cancer 2012; 106: 538–545.

98. Agell L. et al. A 12-gene expression signature is associated with aggressive histological in prostate cancer: SEC14L1 and TCEB1 genes are potential markers of progression. Am J Pathol 2012; 181: 1585–1594.

99. Hong S.H. et al. Upregulation of adenylate cyclase 3 (ADCY3) increases the tumorigenic potential of cells by activating the CREB pathway. Oncotarget 2013; 4: 1791–1803.

100. Rajan P. et al. Identification of novel androgen-regulated pathways and mRNA isoforms through genome-wide exon-specific profiling of the LNCaP transcriptome. PLoS One 2011; 6: e29088.

101. Baty F. et al. EGFR exon-level biomarkers of the response to bevacizumab/erlotinib in non-small cell lung cancer. PLoS One 2013; 8: e72966.

102. Klco J.M. et al. Genomic impact of transient low-dose decitabine treatment on primary AML cells. Blood 2013; 121: 1633–1643.

103. Erdem-Eraslan L. et al. Intrinsic molecular subtypes of glioma are prognostic and predict benefit from adjuvant procarbazine, lomustine, and vincristine chemotherapy in combination with other prognostic factors in anaplastic oligodendroglial brain tumors: a report from EORTC study 26951. J Clin Oncol 2013; 31: 328–336.

104. Mack S,C. et al. Epigenomic alterations define lethal CIMP-positive ependymomas of infancy. Nature 2014; 506: 445–450.

105. Gerber J.M. et al. Genome-wide comparison of the transcriptomes of highly enriched normal and chronic myeloid leukemia stem and progenitor cell populations. Oncotarget 2013; 4: 715–728.

106. Tripathi V. et al. Long noncoding RNA MALAT1 controls cell cycle progression by regulating the expression of oncogenic transcription factor B-MYB. PLoS Genet 2013; 9: e1003368.

107. Erho N. et al. Discovery and validation of a prostate cancer genomic classifier that predicts early metastasis following radical prostatectomy. PLoS One 2013; 8: e66855.

108. Zhao S.G. et al. The Landscape of Prognostic Outlier Genes in High-Risk Prostate Cancer. Clin Cancer Res 2016; 22: 1777–1786.

109. Valletti A. et al. Genome-wide analysis of differentially expressed genes and splicing isoforms in clear cell renal cell carcinoma. PLoS One 2013; 8: e78452.

110. Silva-Martinez G.A. et al. Arachidonic and oleic acid exert distinct effects on the DNA methylome. Epigenetics 2016; 11: 321–334.

111. Mitra A.P. et al. Discovery and validation of novel expression signature for postcystectomy recurrence in high-risk bladder cancer. J Natl Cancer Inst 2014; 106.

112. Karnes R.J. et al. Validation of a genomic classifier that predicts metastasis following radical prostatectomy in an at risk patient population. J Urol 2013; 190: 2047–2053.

113. Tsai H. et al. Cyclin D1 Loss distinguishes prostatic small-cell carcinoma from most prostatic adenocarcinomas. Clin Cancer Res 2015; 21: 5619–5629.

114. Zhao S.G. et al. The Landscape of Prognostic Outlier Genes in High-Risk Prostate Cancer. Clin Cancer Res 2016; 22: 1777–1786.

115. Klein E.A. et al. A genomic classifier improves prediction of metastatic disease within 5 years after surgery in node-negative high-risk prostate cancer patients managed by radical prostatectomy without adjuvant therapy. Eur Urol 2015; 67: 778–786.

116. Jager W. et al. Patient-derived bladder cancer xenografts in the preclinical development of novel targeted therapies. Oncotarget 2015; 6: 21522–21532.

117. Teicher B.A. et al. Sarcoma cell line screen of oncology drugs and investigational agents identifies patterns associated with gene and microRNA expression. Mol Cancer Ther 2015; 14: 2452–2462.

118. Hu Z., Jiang K., Frank M.B., Chen Y., Jarvis J.N. Complexity and specificity of the neutrophil transcriptomes in juvenile idiopathic arthritis. Sci Rep 2016; 6: 27453.

119. Morgan L.Z. et al. Quantitative trait locus and brain expression of HLA-DPA1 offers evidence of shared immune alterations in psychiatric disorders. Microarrays (Basel) 2016; 5: 6.

120. French P.J. et al. Identification of differentially regulated splice variants and novel exons in glial brain tumors using exon expression arrays. Cancer Res 2007; 67: 5635–5642.

121. Dapas M., Kandpal M., Bi Y., Davuluri R.V. Comparative evaluation of isoform-level gene expression estimation algorithms for RNA-seq and exon-array platforms. Brief Bioinform 2017; 18: 260–269.

122. Griffith M. et al. Alternative expression analysis by RNA sequencing. Nat Methods 2010; 7: 843–847.

123. Barrett T. et al. NCBI GEO: archive for functional genomics datasets--update. Nucleic Acids Res 2013; 41: D991–995.

124. Brazma A. et al. ArrayExpress--a public repository for microarray gene expression data at the EBI. Nucleic Acids Res 2003; 31: 68–71.

125. Piccolo S.R. et al. A single-sample microarray normalization method to facilitate personalized-medicine workflows. Genomics 2012; 100: 337–344.

126. Piccolo S.R., Withers M.R., Francis O.E., Bild A.H., Johnson W.E. Multiplatform single-sample estimates of transcriptional activation. Proc Natl Acad Sci U S A 2013; 110: 17778–17783.

127. Tarazona S., Garcia-Alcalde F., Dopazo J., Ferrer A., Conesa A. Differential expression in RNA-seq: a matter of depth. Genome Res 2011; 21: 2213–2223.

128. Sowalsky AG. et al. Whole transcriptome sequencing reveals extensive unspliced mRNA in metastatic castration-resistant prostate cancer. Mol Cancer Res 2015; 13: 98–106.

129. Adiconis X. et al. Comparative analysis of RNA sequencing methods for degraded or low-input samples. Nat Methods 2013; 10: 623–629.

130. Kim D. et al. TopHat2: accurate alignment of transcriptomes in the presence of insertions, deletions and gene fusions. Genome Biol 2013; 14: R36.

131. Kent W.J. et al. The human genome browser at UCSC. Genome Res 2002; 12: 996–1006.

132. Karolchik D. et al. The UCSC Table Browser data retrieval tool. Nucleic Acids Res 2004; 32: D493–496.

133. Li H. et al. The Sequence Alignment/Map format and SAMtools. Bioinformatics 2009; 25: 2078–2079.

134. Ramirez F. et al. deepTools2: a next generation web server for deep-sequencing data analysis. Nucleic Acids Res 2016; 44: W160–W165.

135. Sims D., Sudbery I., Ilott N.E., Heger A., Ponting C.P. Sequencing depth and coverage: key considerations in genomic analyses. Nat Rev Genet 2014; 15: 121–132.

136. Livak K.J., Schmittgen T.D. Analysis of relative gene expression data using real-time quantitative PCR and the 2(-Delta Delta C(T)) Method. Methods 2001; 25: 402–408.

137. Hormaechea-Agulla D. et al. Ghrelin *O*-acyltransferase (GOAT) enzyme is overexpressed in prostate cancer, and its levels are associated with patient’s metabolic status: Potential value as a non-invasive biomarker. Cancer Lett 2016; 383: 125–134.

138. Vandesompele J. et al. Accurate normalization of real-time quantitative RT-PCR data by geometric averaging of multiple internal control genes. Genome Biol 2002; 3: research0034.1-research0034.11.

139. Gruber A.R., Bernhart S.H., Lorenz R. The ViennaRNA web services. Methods Mol Biol 2015; 1269: 307–326.

140. Wan Y., Kertesz M., Spitale R.C., Segal E., Chang H.Y. Understanding the transcriptome through RNA structure. Nat Rev Genet 2011; 12: 641–655.

141. Zuker M., Stiegler P. Optimal computer folding of large RNA sequences using thermodynamics and auxiliary information. Nucleic Acids Res 1981; 9: 133–148.

142. Shultz L.D. et al. Human lymphoid and myeloid cell development in NOD/LtSz-scid IL2R gamma null mice engrafted with mobilized human hemopoietic stem cells. J Immunol 2005; 174: 6477–6489.

143. Liao Y., Smyth G.K., Shi W. featureCounts: an efficient general purpose program for assigning sequence reads to genomic features. Bioinformatics 2014; 30: 923–930.

144. Robinson M.D., McCarthy D.J., Smyth G.K. edgeR: a Bioconductor package for differential expression analysis of digital gene expression data. Bioinformatics 2010; 26: 139–140.

145. Robinson M.D., Oshlack A. A scaling normalization method for differential expression analysis of RNA-seq data. Genome Biol 2010; 11: R25.

146. Law C.W., Chen Y., Shi W., Smyth G.K. voom: Precision weights unlock linear model analysis tools for RNA-seq read counts. Genome Biol 2014; 15: R29.

147. Ritchie M.E. et al. limma powers differential expression analyses for RNA-sequencing and microarray studies. Nucleic Acids Res 2015; 43: e47.

148. Wang E.T. et al. Alternative isoform regulation in human tissue transcriptomes. Nature 2008; 456: 470–476.

149. Durinck S., Spellman P.T., Birney E., Huber W. Mapping identifiers for the integration of genomic datasets with the R/Bioconductor package biomaRt. Nat Protoc 2009; 4: 1184–1191.

150. Huang da W., Sherman B.T., Lempicki R.A. Systematic and integrative analysis of large gene lists using DAVID bioinformatics resources. Nat Protoc 2009; 4: 44–57.

151. Shannon P. et al. Cytoscape: a software environment for integrated models of biomolecular interaction networks. Genome Res 2003; 13: 2498–2504.

152. Suderman M., Hallett M. Tools for visually exploring biological networks. Bioinformatics 2007; 23: 2651–2659.

153. Benjamini Y., Hochberg Y. Controlling the false discovery rate: a practical and powerful approach to multiple testing. J R Stat Soc Series B Stat Methodol 1995; 1: 289–300.

154. Luck S., Thurley K., Thaben P.F., Westermark P.O. Rhythmic degradation explains and unifies circadian transcriptome and proteome data. Cell Rep 2014; 9: 741–751.

155. Simola D.F. et al. A chromatin link to caste identity in the carpenter ant *Camponotus floridanus*. Genome Res 2013; 23: 486–496.

156. Conway T. et al. Xenome--a tool for classifying reads from xenograft samples. Bioinformatics 2012; 28: i172–178.

157. Subramanian A. et al. Gene set enrichment analysis: a knowledge-based approach for interpreting genome-wide expression profiles. Proc Natl Acad Sci U S A 2005; 102: 15545–15550.

158. Huber W. et al. Orchestrating high-throughput genomic analysis with Bioconductor. Nat Methods 2015; 12: 115–121.

159. Davis S., Meltzer P.S. GEOquery: a bridge between the Gene Expression Omnibus (GEO) and BioConductor. Bioinformatics 2007; 23: 1846–1847.

160. Goldman M. et al. The UCSC Cancer Genomics Browser: update 2015. Nucleic Acids Res 2015; 43: D812–817.

161. Vivian J. et al. Toil enables reproducible, open source, big biomedical data analyses. Nat Biotechnol 2017; 35: 314–316.

162. Cerami E. et al. The cBio cancer genomics portal: an open platform for exploring multidimensional cancer genomics data. Cancer Discov 2012; 2: 401–404.

163. Gao J. et al. Integrative analysis of complex cancer genomics and clinical profiles using the cBioPortal. Sci Signal 2013; 6: pl1.

164. Hartigan J.A., Wong M.A. Algorithm AS 136: A k-means clustering algorithm. J R Stat Soc Ser C Appl Stat 1979; 28: 100–108.

165. Quackenbush J. Computational analysis of microarray data. Nat Rev Genet 2001; 2: 418–427.

166. Weichselbaum R.R. et al. An interferon-related gene signature for DNA damage resistance is a predictive marker for chemotherapy and radiation for breast cancer. Proc Natl Acad Sci U S A 2008; 105: 18490–18495.

167. Rich J.T. et al. A practical guide to understanding Kaplan-Meier curves. Otolaryngol Head Neck Surg 2010; 143: 331–336.

168. Therneau T. A package for survival analysis in S, doi: 10.1093/survival/43.3.1 (2015).

169. Cox D.R. Regression models and life-tables. Breakthroughs in Statistics. 527–541 (Springer, 1992).

170. Efron B. The efficiency of Cox’s likelihood function for censored data. J Am Stat Assoc 1977; 72: 557–565.

